# Evolutionary Rewiring of Root Cell-Type Nitrogen Responses

**DOI:** 10.64898/2026.06.05.730053

**Authors:** Zhengao Di, Deeksha Singh, Ashleigh Lister, Yao Min Cai, Chao Bian, Mohana Talasila, Yuxuan Lan, Suzanne Henderson, Fiona Fraser, Tom Barker, Thomas Brabbs, Kendall Baker, Leah Catchpole, David Swarbreck, R. Glen Urhig, Iain Macaulay, Wilfried Haerty, Siobhan M. Brady, Nicola J. Patron

## Abstract

Nitrate induces widespread transcriptional reprogramming in plants, but how transcription factors coordinate these responses across cell types and evolve across species with divergent root patterning remains unclear. Using single-cell transcriptomics in *Arabidopsis thaliana* and *Solanum lycopersicum* (tomato) under varying nitrate conditions, we demonstrate that many transcriptional responses are cell type and developmental stage-specific. Arabidopsis NIN-LIKE PROTEIN 7 (NLP7) directly modulates both broadly expressed and cell-type-specific targets, but is not required to establish the cell type specific nature of these patterns. Comparative analysis reveals evolutionary shifts; key nitrogen programs are localized to different cell types in tomato, with the exodermis as a major regulatory hub. Identification of tomato NLP7a and NLP7b direct targets at cellular resolution supports their role in direct modulation of cell type N-transcriptional programs. This is evidenced by NLP7-controlled exodermal cell wall remodeling in low nitrate, highlighting functional conservation despite divergent cellular topographies.

## Introduction

Nitrogen (N) is a fundamental macronutrient that is essential for plant growth and development. Plants absorb N through their roots as inorganic nitrate (NO^-^_3_) and ammonium (NH^+^_4_). Previous studies have shown that fluctuations in the availability of nitrate triggers large-scale shifts in gene expression, which are orchestrated by complex networks of transcription factors (TFs) ^1–6^. Central to these responses is the coordination of NIN-LIKE PROTEIN 7 (NLP7). In Arabidopsis, AtNLP7 has been shown to directly sense nitrate, responding via rapid subcellular relocalization to initiate transcriptional reprogramming via numerous target genes ^7–12^. However, *AtNLP7* shows no nitrate-induced differential expression in whole-root studies ^7–12^. AtNLP7 interacts with its homolog, AtNLP6, to coordinate nitrate signaling by forming both homodimers and heterodimers, primarily mediated by their C-terminal PB1 (Phox and Bem1p) domains ^10,13,14^. AtNLP6 and AtNLP7 are partially functionally redundant. Arabidopsis *nlp6/nlp7* double mutants display a more severe loss of nitrate response phenotype than single mutants ^5,10^. A number of other transcription factors (TFs) have been associated with the coordination of N-responses in Arabidopsis ^15–18^. Notably, Arabidopsis *TGACG-BINDING FACTOR 1* and *4* (*AtTGA1* and *AtTGA4*) transcripts have been shown to significantly increase in response to nitrate ^4,6^. AtTGA1 and 4 are thought to be partly functionally redundant, coordinating the expression of a large number of downstream genes, many of which are also targets of AtNLP6 and AtNLP7, notably genes involved in primary metabolism and nitrate transport such as *AtNRT2*.*1* and *AtNRT2*.*2* ^4–6^.

In Arabidopsis, it is well-established that N availability affects several distinct programs such as root growth and flowering time ^19^. Some nitrate responses, for example lateral root initiation, have been clearly associated with a specific cell type. However, it remains unknown whether other responses, such as autophagy in low nitrate, are limited or occur across all cells ^20,21^. Early investigations of cell type nitrate responses profiled pools of protoplasts isolated from cell type- or tissue-specific GFP-expressing reporter lines enriched by fluorescence-activated cell sorting (FACS). This revealed coordinated yet functionally distinct cellular responses to nitrate ^22,23^. More recent studies used similar approaches to investigate the nitrate responsive regulatory networks of Arabidopsis endodermis, which supported a role for this cell layer as a regulatory hub of nitrate responses in which ABF2 and ABF3 play key roles ^24^. Other studies have mapped the downstream targets of AtNLP7 and AtTGA1 using methods such as transient assay reporting genome-wide effects of transcription factors (TARGET) ^1–5^. However, significant gaps in our understanding remain. First, it is unclear if or how such TFs coordinate the nitrate-responsive programs that occur in distinct cell types. Second, it remains unknown if cell type-specific nitrate responses are conserved in species with divergent root cellular architectures, such as *Solanum lycopersicum* (tomato). Recent comparative studies indicate that while some gene regulation is conserved between Arabidopsis and tomato, key differences exist; for instance, N regulation has been reported to be enriched in the suberized endodermis of Arabidopsis, whereas in tomato, this enrichment shift was predicted to occur in lignified and suberized exodermis ^22,25^.

The rapidly evolving field of single-cell transcriptomics (scRNA-seq) enables the investigation of cellular transcriptional programs. ScRNA-seq was first employed in Arabidopsis roots, generating transcriptional datasets for over tens of thousands of cells, identifying marker genes ^26–28^ and mapping gene expression changes during cell type differentiation ^29^). Several studies have used these methods to profile the responses of specific cell types to changes in the environment including phosphate ^30^, sucrose ^26^, heat ^31^, and compaction ^32^, as well as to developmental signals such as brassinosteroids ^33^. A scRNA-seq analysis of single nuclei from wheat roots grown in varying N concentration highlighted differences in the population of epidermal cells and their role in N metabolism ^34^, while a similar recent study in tomato highlighted N-responsive metabolic programs ^35^.

In this study, we compare scRNA-seq of protoplasts of Arabidopsis and tomato roots from plants grown in limiting and sufficient KNO_3_ conditions to address the cellular nature of nitrate responses in evolutionarily and anatomically divergent species, and elucidate the role of NLP7, its paralogs and orthologs in establishing these patterns. Our results support previous findings that different root cell types exhibit highly varied responses to N, but also show that cell type differentiation stage is an important component of N-response, specifically in sub-populations of mature cortex, endodermis, and pericycle cells. We show that AtNLP7, AtTGA1 and AtTGA4 modulate the expression of both broadly expressed and cell-type-specific target genes. However, AtNLP7 is not required for the cell-specific expression patterns of its target genes. Furthermore, we reveal that a striking spatial shift in key N-responsive programs in tomato compared to Arabidopsis, providing evidence for the exodermis as the regulatory hub of nitrate responses in the species. Finally, we explore the conservation and divergence of the NLP7 regulatory network across tomato and Arabidopsis, identifying direct targets of tomato NLP7a and NLP7b at cellular resolution. Exodermal cell wall remodeling is controlled by SlNLP7a in differing nitrate conditions. Collectively, these data provide a comparative evolutionary blueprint of root spatiotemporal N responses.

## Results

### Detection of nitrate responses in single cell transcriptomes

A key aim of this study was to investigate how limiting and sufficient nitrate inform transcriptional programs at spatiotemporal resolution and the role of key N-regulatory TFs in these programs. To obtain single cell transcriptomes of Arabidopsis root tissues, we grew plants on sterile media with either 1 mM (limiting) or 10 mM (sufficient) KNO_3_. Whole roots of 12 day old seedlings were harvested from ∼2,000 roots and protoplasts were extracted. Two independent replicate pools of protoplasts were used to prepare libraries for scRNAseq (Supplementary Figure S1). In total, we captured 123,489 Arabidopsis cells with an average of 69,665 reads and a median of 1,222 genes per cell (Supplementary Figure S2; Supplementary Dataset 1). Following alignment and filtering, cell types were annotated both by label transfer using an existing Arabidopsis root atlas as a reference and the index of cell identity (ICI) method ^29,36^. Developmental stages were predicted by label transfer from the Arabidopsis root atlas (Supplementary Figures S3-4). Annotations revealed ten major groups of cells: two types of epidermal cells (atrichoblast and trichoblast); the cortex, endodermis, pericycle, procambium, phloem, and xylem; a population of mixed identities comprised of quiescent center, inner columella and outer lateral root cap cells (hereafter referred to as “root cap”); and cells in proliferative stages of mixed identities (hereafter referred to as meristematic) (Figure 1a)(Supplementary Figures S4-5). All annotated cell types were validated using marker genes with known expression patterns (Supplementary Figure S6, Dataset S2).

**Figure 1.**
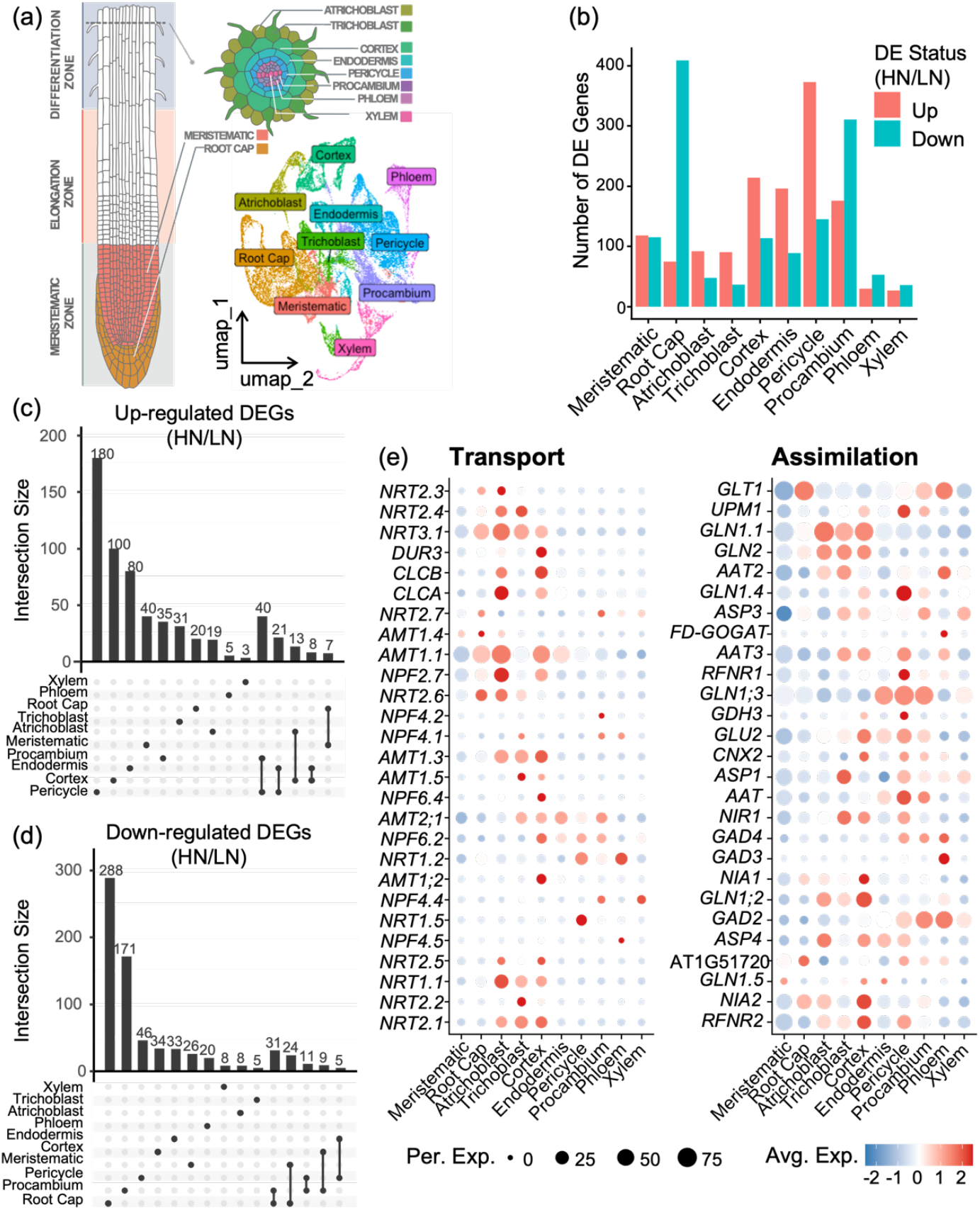
Arabidopsis root cell types show varied responses to nitrate availability. (a) Visualization of ten cell type clusters. Each dot denotes a single cell. Colors denote cell clusters. (b) Quantification of differentially expressed genes detected in each cell type in 10 mM KNO_3_ (high/sufficient nitrate; HN) vs 1 mM KNO_3_ (limiting nitrate; LN). (c, d) Overlaps of genes up- (c) and down-regulated (d) in HN vs LN. (e, f) Cell-type expression of genes involved in nitrogen transport (e) and assimilation (f).

The influence of protoplasting on cell type identity in these N conditions was assessed. First, RNA-seq was performed on whole roots grown in the same KNO_3_ conditions (bulk transcriptome) and N-dependent differential gene expression in the bulk transcriptome data was compared to that observed in the combined population of all single cells (pseudobulk transcriptome) (Supplementary Dataset S3). Principal component analysis (PCA) indicated that while the preparation of protoplasts expectedly affects gene expression, KNO_3_ responses detected in the pseudobulk transcriptome are similar to those observed in the bulk transcriptome (Supplementary Figure S7). Previous studies have defined a set of genes affected during protoplast isolation ^37^. Cell clustering and annotation analyses performed with and without the removal of these genes from the datasets further indicated that they did not impact the outcome (Supplementary Figure S8). The single cell transcriptome also identified N-induced DEGs that were not differentially expressed in the bulk transcriptome, highlighting the potential of this dataset for discovering previously unknown N-responsive processes (Supplementary Figure S7). Therefore we chose to proceed to spatiotemporal analysis without removal of protoplasting-induced genes.

### Arabidopsis root cell types show varied responses to nitrate

We analyzed and compared the responses of the ten Arabidopsis root cell types to KNO_3_. Differential expression analysis was performed using a Wilcoxon rank sum test to compare 1 mM (limiting) and 10 mM (sufficient) KNO_3_ for each cell type (Supplementary Dataset S4). The abundance of cells annotated as root cap was enriched in samples from roots grown in high KNO_3_. Consistent with previous studies, roots grown in high KNO_3_ were shorter than those grown in limiting KNO_3_^3,11^. As the whole root was sampled, these cells comprised a larger proportion of the total cell population than in samples from limiting KNO_3_ in which roots were longer (Supplementary Figures S1 and S9). Cell types differed in their transcription when grown in different KNO_3_ concentrations: the cortex, endodermis, pericycle, procambium and “root cap” tissue had the highest proportion of differentially expressed genes (DEGs), while meristematic, atrichoblast, phloem and xylem cells had the lowest proportion of DEGs (Figure 1b). The endodermis, pericycle, cortex had the largest numbers of genes upregulated in high KNO_3_. Some genes were only differentially expressed in one cell type while the expression of others responded to KNO_3_ in multiple cell-types (Figure 1c). In contrast, the procambium and root cap had the largest number of genes downregulated by high KNO_3_ (Figure 1d). We also performed Pearson correlations of the pseudobulked transcriptomes of each cell type in limiting and sufficient KNO_3_, which indicated that the procambium, “root cap” and pericycle had large numbers of DEGs while trichoblast had only a small number of DEGs that respond very strongly (Figure 1b; Supplementary Figure S10). Despite differences in the number of DEGs and KNO_3_ responsiveness, GO term analysis of DEGs in each cell type identified several terms associated with responses to environmental stimuli that are common to multiple cell types(Supplementary Figures S11-S12).

We examined the cellular expression profiles of 54 genes known to be involved in N transport and assimilation (Figure 1e). Of these, 41 were found to be differentially expressed in the bulk transcriptome (Supplementary Figure S13) and 35 were differentially expressed in at least one cell type (Supplementary Figure S14). We also performed proteomic analyses of whole roots grown in the same KNO_3_ conditions (bulk proteome) and investigated the abundance of the gene products (Supplementary Dataset S5). We were able to detect 31 of these proteins, of which 14 showed differential abundance in limiting/sufficient KNO_3_ (Supplementary Figure S15).

Finally, we benchmarked our single cell transcriptome data against previous studies that used FACS to collect populations of protoplasts enriched in target cell types from plants expressing cell-type specific fluorescent markers ^6,22,23^. Despite differences in the growth conditions and expression profiling methods, common N-regulated genes were identified in the pericycle and cortex in all studies (Supplementary Figure S16).

### Nitrate-responsive genes are enriched in sub-populations of mature cells

Next, we mapped the expression of genes previously identified using time course transcriptomics following NO_3_^-^ and NH_4_^+^ supply in order to investigate N-responsiveness at cell type level ^2^. These 2,324 genes included 357 genes that were also differentially expressed in our bulk transcriptome of roots grown in limiting and sufficient KNO_3_ (Supplementary Figure S17). Interestingly, while the expression of the N-responsive genes was expectedly enriched in several of the cell types observed to be the most KNO_3_ responsive (endodermis, cortex, pericycle, procambium), their expression was non-uniform across these cell types with some cells showing higher expression (Figure 2a; Supplementary Figure S17). To further investigate, we reclustured the most N-responsive cell types (endodermis, cortex, pericycle) at a higher resolution (Figure 2b-d). We next asked if cells are responsive to N at a specific developmental stage within each trajectory. To obtain a high resolution of developmental gradient within each cell type, developmental potential was calculated using CytoTRACE, which estimates developmental potential from transcriptional diversity ^38^. CytoTRACE scores correlate with the developmental predictions obtained via label transfer and expression of known cell cycle genes (Supplementary Figure S18). This analysis revealed that while each cell type contained more than one sub-population of similar maturity, only one of these was enriched in N-responsive genes (Figure 2e-g; Supplementary Figures S19-23). We confirmed that gene expression in response to KNO_3_ within the sub-populations enriched in N-responsive genes (hereafter referred to as responsive) is significantly different to the other sub-population(s) of mature cells of the same cell types (hereafter referred to as non-responsive) (Figure 2h; Supplementary Dataset S6). A GO-term enrichment analysis of DEGs enriched in responsive sub-populations of all three cell types indicated the presence of both shared and specific functions (Figure 2i, Supplementary Figure S24). Shared programs were enriched in stress responses (responses to hypoxia, wounding, water, heat, jasmonic acid and biotic stimulus). Cortex programs included cation transport and defense programs; the pericycle was enriched for cell division programs; and the endodermis was enriched in cell wall loosening and metabolic processes (aromatic amino acids and indole compounds) (Figure 2i, Supplementary Figure S24).

**Figure 2.**
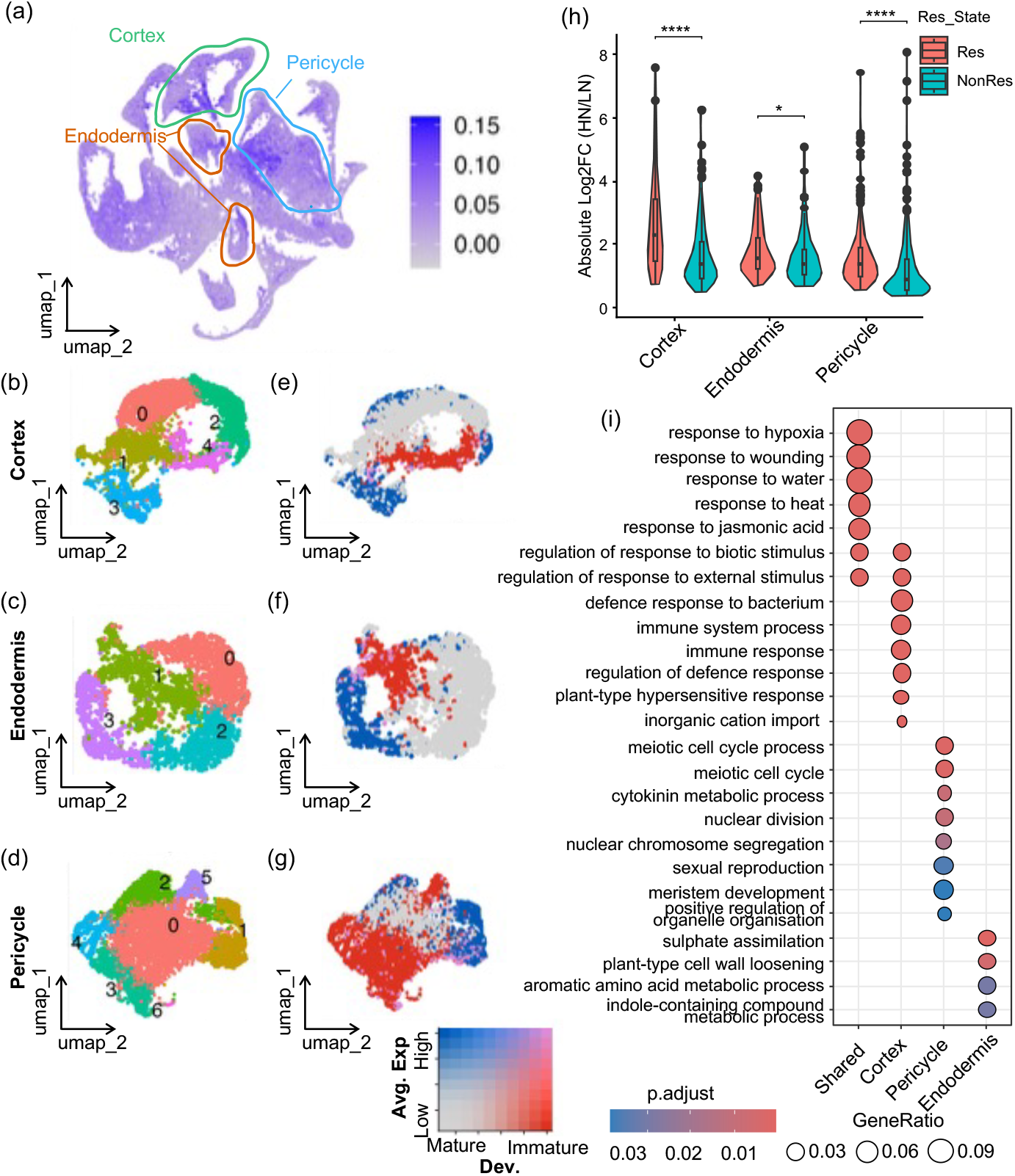
N-responsive genes are enriched in mature cortex, endodermis and pericycle subpopulations. (a) UMAP annotated with expression of known N-responsive genes (b-d) Sub-cluster UMAPs of cortex, endodermis and pericycle (e-g) Correlation of N-responsive genes with maturity in sub-clusters (h) Sub-clusters enriched in N-responsive gene expressions have higher KNO_3_ responses. Absolute log_2_ fold change (HN/LN) of N-responsive genes in responsive (Res) and non-responsive (NonRes) sub-populations of mature cortex, endodermis, and pericycle cells. Genes with adjusted p-value < 0.05 are included. Differences between sub-populations were assessed using the Wilcoxon rank-sum test with Benjamini-Hochberg correction. ns: p.adj > 0.05, *: p.adj < 0.05, ****: p.adj < 0.0001. Boxes show values between the first and third quartiles, and bold lines show median values. (i) GO enrichment of DEGs enriched in Res sub-clusters of each cell type, showing responsive sub-pops are specialized in different functions in each cell type.

### Nitrogen-regulated target genes show varied cell-type expression profiles

Next, we explored the role of AtNLP7, AtTGA1 and AtTGA4 and their direct targets in the N-responsive subpopulations of cortex, endodermis and pericycle. Direct targets were obtained from previously TARGET studies ^4,5^. Genes previously reported to be directly upregulated by AtNLP7, AtTGA1 or TGA4, increased expression in high KNO_3_, particularly in the responsive subpopulations of the cortex and pericycle in which the expression of *AtTGA1* was also observed to be upregulated in high KNO_3_ (Figure 3a). Direct targets of AtTGA1 and AtTGA4 previously reported to be down-regulated were lower in the responsive cortex subpopulation but this was less apparent in the endodermis and pericycle. Similarly, the expression of target gene reported to be directly down-regulated by AtNLP7 was not similar to the relative abundance of *AtNLP7* transcripts (Figure 3a). The pericycle contained the largest number of DEGs that are targets of both NLP7 and TGA1/4 (Supplementary Figure S25).

**Figure 3.**
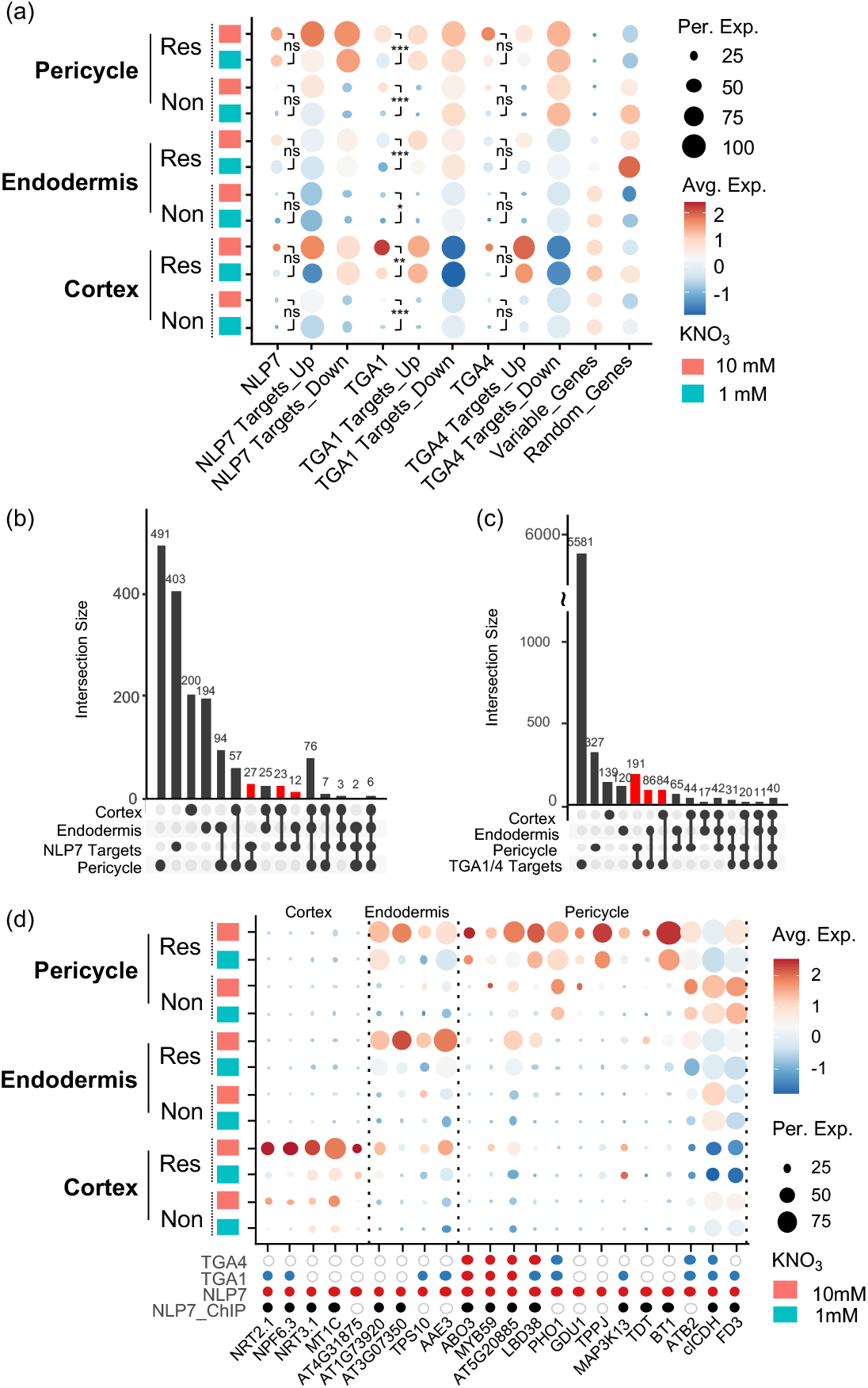
Transcriptional coordinators of nitrogen responses regulate targets with broad and cell type-specific expression patterns. **(a)** Expression of Arabidopsis N-responsive genes; selected transcription factors (*AtNLP7, AtTGA1* and *AtTGA4*); and their target genes in responsive and non-responsive subpopulations of pericycle, endodermis and cortex cells at limiting (1 mM) and sufficient (10 mM) KNO_3_. Variable genes = 2000 genes with high cell-to-cell variation used for clustering and integration. Random genes = 2652 randomly selected genes to confirm that subclusters are unbiased. ns: p.adj > 0.05, * p.adj < 0.05, ** p.adj < 0.01, *** p.adj < 0.001.. **(b-c)** Targets of AtNLP7 (b) and AtTGA1/4 (c) have broad and cell-type-specific (red bars) differential expression profiles. **(d)** Directly upregulated targets of AtNLP7 that are differentially expressed in responsive subpopulations of the pericycle, endodermis or cortex. Red and blue dots indicate direct activation or repression, respectively, in TARGET assays with corresponding TF; empty circles denote no known regulation; black dots denote AtNLP7 binding detected by ChIP.

Previously, AtABF2 and AtABF3 have been shown to be key regulators of N responses, particularly in the endodermis ^24^. We therefore compared the cell type-specific expression of AtABF2 and AtABF3 and their target genes. In agreement with previous studies, we found enrichment and N-induced expression in the endodermis, pericycle, procambium (Supplementary Figure S26).

We then investigated the expression profiles of specific genes that are direct targets of AtNLP7, AtTGA1 and AtTGA4 in the N-responsive sub-populations, finding that while some target genes have broad expression patterns (differentially expressed in multiple cell types), most are differentially expressed in specific cell types (Figure 3b and c). In Figure 3d, we illustrate genes that are directly activated by NLP7 that were significantly upregulated by high KNO_3_ in the cortex, endodermis or pericycle (or responsive sub‐population of these cells) (Figure 3d; Supplementary Figure S27). These include genes previously shown to be involved in N responses. For example, in the responsive cortex, genes encoding transporters *AtNPF6*.*3, AtNRT2*.*1, AtNRT3*.*1*. (Figure 3d; Supplementary Figure S27). The expression of these transporters was also enriched in atrichoblast cells, indicating that cortex and atrichoblast cells share some N-responsive programs (Supplementary Figure S27). Similarly, the pericycle and procambium were also observed to share some N-responsive programs (Supplementary Figure S27, Figure 1c, d). In the responsive subpopulation of endodermal cells *AtTPS10*, which encodes a putative trehalose phosphate synthase, and *AtAAE3*, which encoded oxalyl-CoAsynthetase, involved in oxalate degradation, were upregulated by KNO_3_ (Figure 3d; Supplementary Figure S27). These genes were also upregulated in the pericycle, but not significantly. The responsive pericycle had the largest number of N-responsive targets of AtNLP7. These included genes involved in lateral root development (*AtLBD38*), ABA responses (*AtABO3/AtWRKY63*), and a putative transcription factor involved in K+/NO3-transport and expression of the AtNPF7.3 transporter (*AtMYB59*). We also identified genes involved in nutrient movement (*AtPHO1*), glutamine secretion (*AtGDU1*), energy balancing (*AtTPPJ*), malate/fumarate transport (*AtTDT*) as well as *AtclCDH* (NADP+-dependent isocitrate dehydrogenase), and *AtFD3* (ferredoxin 3) (Figure 3d; Supplementary Figure S27).

### Spatial expression patterns of N-responsive genes are retained in Arabidopsis mutants with perturbed N-responses

Our findings indicate that AtNLP7 regulates the nitrate-responsive expression of target genes that are expressed in multiple cell types as well as those for which expression is enriched in one or more specific cell types. Thus, we hypothesized that AtNLP7 is unlikely to be responsible for the cell type-specific expression patterns of its target genes. To further investigate this, we conducted single-cell transcriptome sequencing of root protoplasts of the *nlp7-1* mutants, grown in the same limiting and sufficient KNO_3_ conditions as the original single cell profiling dataset described here. Arabidopsis *nlp7-1* mutant alleles have previously been shown to have reduced N responses ^3,7,8^. As further validation of the role of NLP7 in these responses, we performed scRNAseq of a mutant allele of the Arabidopsis NAC domain-containing protein 32 (AtANAC032) - a direct negative regulator of *AtNLP7*. Further, *anac032* mutant alleles show an increase in N-responsiveness ^11^.

Clustering and cell-type annotations of *nlp7-1* and *anac032* mutants were performed together with Col-0 in a single integrated Seurat analysis (Supplementary Figure S28). No bias in clusters or cell types by samples or KNO_3_ conditions was observed (Supplementary Figure S29). Sub-clustering analysis was performed independently for each genotype and revealed that N-responsive genes are also enriched in mature sub-populations of cortex, endodermis and pericycle cells within these mutants (Figure 4a; Supplementary Figures S30-S39). Comparison of the spatial expression of AtNLP7 targets showed that most genes retain their cell type expression patterns in the mutants (Figure 4b). Similarly, significant overlaps of DEGs in the cortex, endodermis and pericycle cell types between Col-0 and mutants were found (Supplementary Figure S40), and GO-term enrichment analyses of DEGs indicated that these cell types retained similar N response programs (Supplementary Figure S41). However, fewer genes were differentially expressed in KNO_3_ treatments in *nlp7-1* cortex, endodermis and pericycle, with the greater change being a reduction in the number of downregulated genes. In contrast, all *anac032* cell types had a greater number of DEGs compared to wild type (Supplementary Figure S42). Furthermore, direct targets of AtNLP7 were notably reduced among DEGs enriched in *nlp7-1* cortex, while more direct targets of AtNLP7 were differentially expressed in all *anac032* cell types (Supplementary Figure S42).

**Figure 4.**
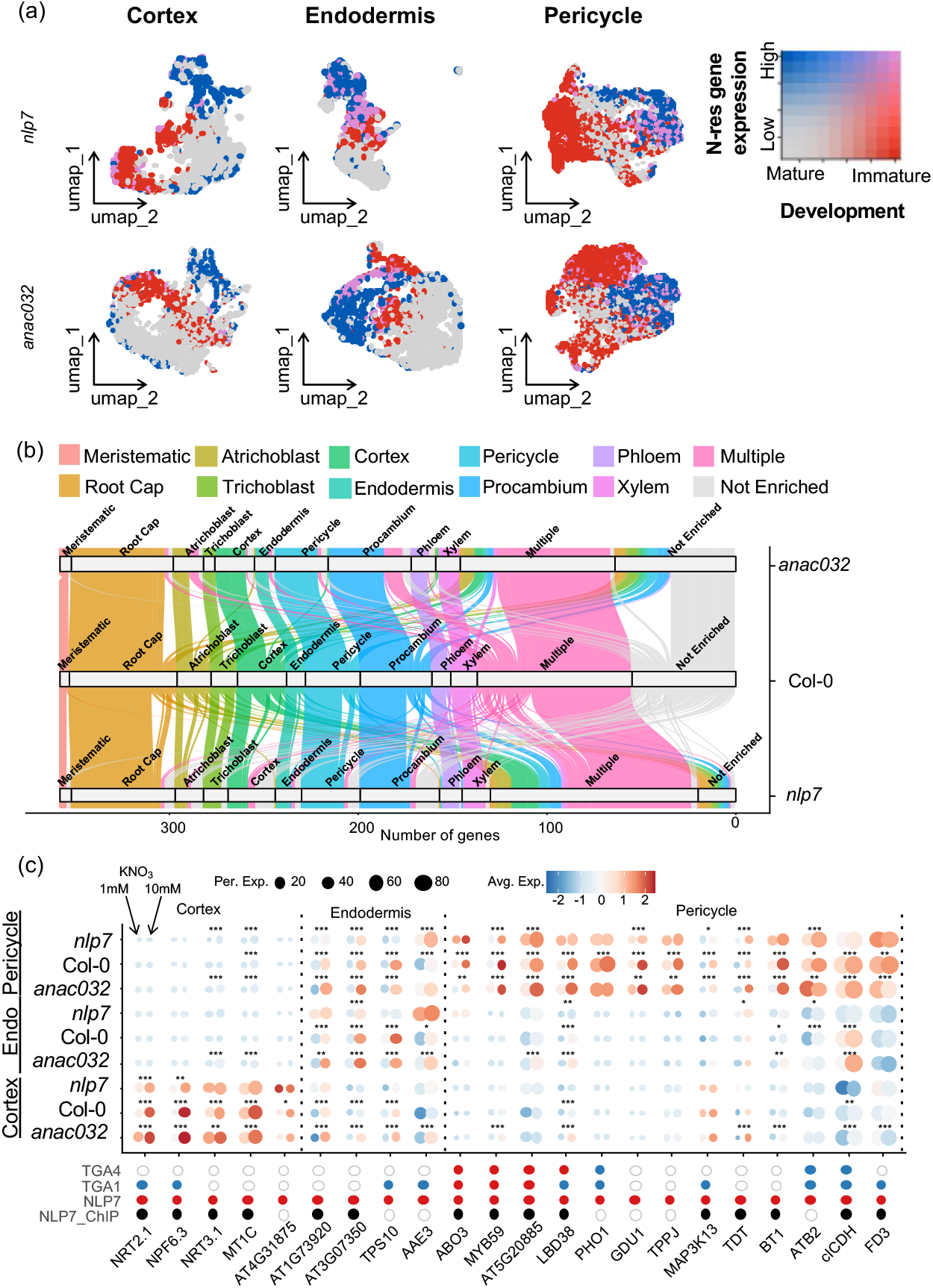
Targets of AtNLP7 retain their cell-type expression profiles in *nlp7* and *anac032* mutants. **(a)** Sub-populations of *nlp7* and *anac032* mature pericycle, endodermis and cortex cells are enriched for N-responsive genes. **(b)** Alluvial plot of the expression profiles of AtNLP7 target genes in Col-0, *nlp7* and *anac032*. Colors reflect expression profiles in Col-0. **(c)** Targets of *AtNLP7* that show differential expression in limiting/sufficient KNO_3_ and cell-type enrichment in Col-0 retain their expression profiles but show perturbed N-responses in *nlp7* and *anac032*. Red and blue dots indicate direct activation or repression, respectively, in TARGET assays with corresponding TF; empty circles denote no known regulation; black dots denote AtNLP7 binding detected by ChIP. Asterisks indicate significant gene expression at sufficient vs limiting KNO_3_ within each genotype. * p.adj < 0.05, ** p.adj < 0.01, *** p.adj < 0.001.

A small proportion of genes that are enriched in a specific cell type in Col-0, lose their cell type-specific enrichment in mutant lines and, conversely, a small proportion of genes that show no cell-type specific enrichment in Col-0 show cell type enrichment in the mutants (Figure 4b). N-responsive targets of NLP7 that showed cell-type specific enrichment (Figure 3d) retained their cell-type enrichment profiles in both mutants (Figure 4c). However, we observed differences in responsiveness to KNO_3_: many targets of AtNLP7 that are not co-regulated by AtTGA1/4 (*NRT3*.*1, MT1C, AT4G31875, AT1G73920, TPPJ* and *BT1*), were observed to have lost or reduced differential expression in *nlp7*, with several showing increased expression in *anac032*, similar to their respect N-responsive root phenotypes (Figure 4c; Supplementary Figure S43). Target genes that are also regulated by TGA1/4 retained significant N condition-differential expression in *nlp7* and *anac032* mutants as did *AT3G0735, GDU1* and *TDT*, perhaps suggesting co-regulation by unknown N-responsive TFs.

### Differences in cell type N-responsiveness in tomato

Arabidopsis and tomato have instances of both conserved and divergent regulatory architecture in the whole root ^11^. Anatomically, Arabidopsis and tomato differ - Arabidopsis has a single cortex layer and lacks an exodermal layer. Tomato has three layers of cortical cells, the outermost of which is the exodermis. Recent work in tomato found that genes associated with N metabolism were enriched in an inferred exodermal regulatory network ^25^. We intersected an RNAseq dataset from the whole tomato roots in differing N concentrations ^11^, and found it was enriched for a previously identified exodermis-enriched gene population (Supplementary Figure S44) ^25^.

To further investigate cell type N-dependent gene expression patterns, we performed single cell transcriptome sequencing of two pools (independent biological replicates) of tomato protoplasts isolated from plant roots grown in 1 mM and 10 mM KNO_3_. In total, we captured 23,929 tomato cells with an average of 47,928 reads and a median of 1,787 genes per cell (Supplementary Figure S45; Supplementary Dataset S7). Following alignment and filtering, cell types were annotated by label transfer using a previously annotated tomato root atlas as a reference ^39^. The expression profiles of cluster markers were verified by comparison to cell type-enriched transcriptomes obtained using FACS ^25^ (Figure 5a; Supplementary Figures S46-S47). Cluster annotations were validated using known marker genes (Supplementary Figure S48) and developmental scores were calculated using CytoTRACE ^38^.

**Figure 5.**
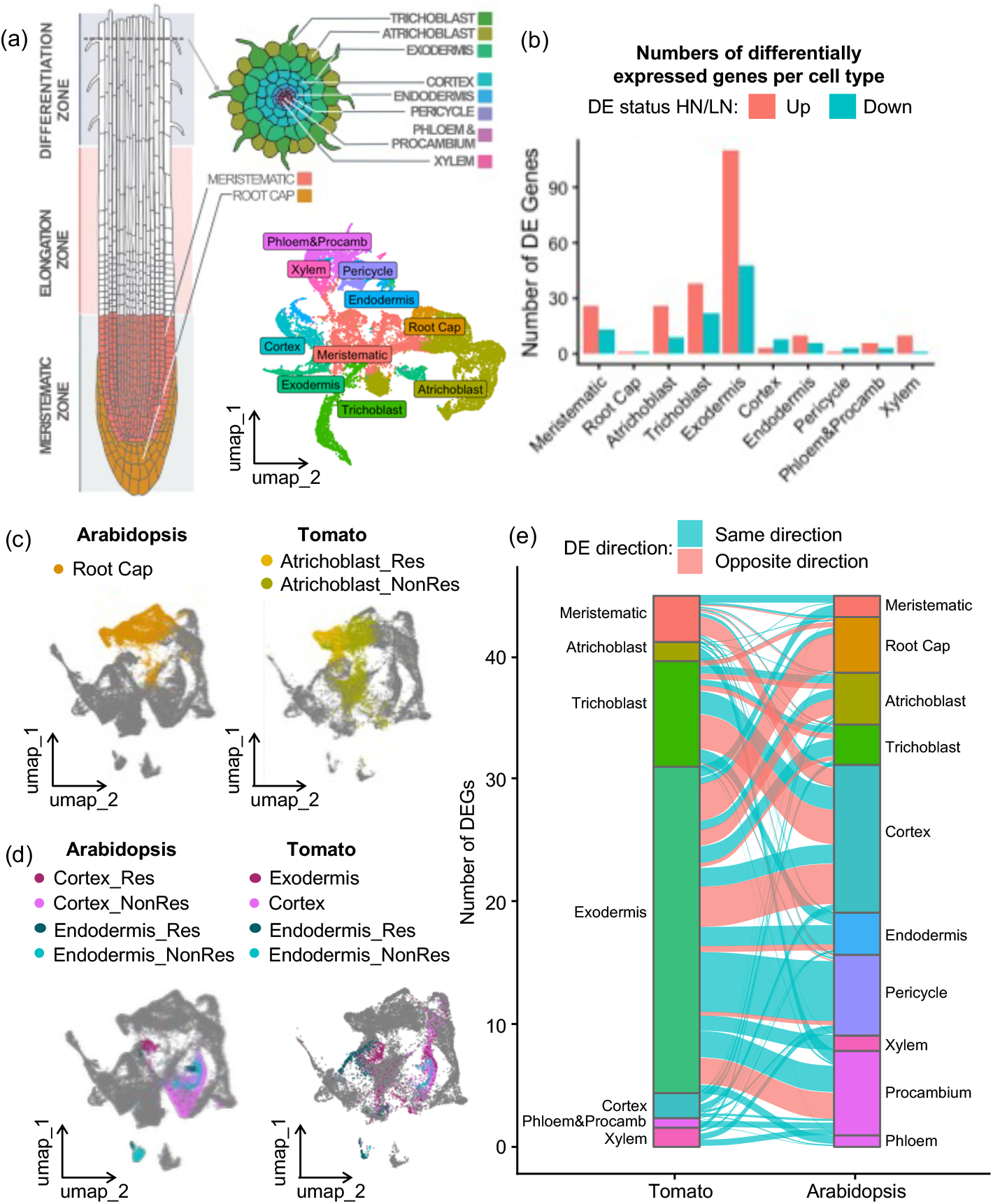
Transcriptional N-responses in tomato are enriched in the exodermis. (**a**) Tomato root cell clusters (UMAP) annotated by cell type. (**b**) Quantification of differentially expressed genes (DEGs) detected in each cell type in high/sufficient KNO_3_ (HN) vs limiting KNO_3_ (LN). (**c, d**) Integrated UMAPs of tomato and Arabidopsis using combined transcripts of all tomato expressologs of Arabidopsis genes, showing root cap and atrichoblast cells (c) and exodermis, endodermis and cortex cells (d). (**e**) Cell-type locations of tomato and Arabidopsis expressologs that are differentially expressed in HN vs LN in both species.

Differential expression analysis was performed as described for Arabidopsis (Supplementary Dataset S8. In tomato, we observed the exodermis to be the most highly N-responsive cell type, both in terms of up-regulated and down-regulated genes, consistent with the predicted exodermal network, and overlap of whole root N-responsive gene expression (Figure 5d; Supplementary Figure S49). Similar to Arabidopsis, we also performed RNA-seq on whole roots grown in the same KNO_3_ conditions. This identified 1199 genes that were significantly differentially expressed (Supplementary Dataset S9). These 1199 DEGs (termed N-DEGs) were most highly expressed in exodermis, endodermis and atrichoblast (Supplementary Figure S50). We then performed sub-clustering analysis of each tomato cell type in the same manner as described above for Arabidopsis, identifying similar sub-populations of mature cells in the endodermis and atrichoblast (Supplementary Figures S51-52; Supplementary Dataset S10). In contrast, exodermal cells were evenly enriched in the expression of N-DEGs. Sub-populations were not found in other cell types in which N-DEGs genes were not highly expressed.

In order to determine how similar the single cell transcriptional responses were between species, orthologous gene sets were used to integrate single cell transcriptomes of Arabidopsis and tomato with clustering visualized on UMAPs (Supplementary Figures S53-60). Several approaches were used to construct shared features for integration - aggregating transcripts across tomato and Arabidopsis expressologs (identified as described in ^25^) (Supplementary Figure S53), as well as aggregating transcripts across all genes within an EnsemblPlant homolog group (Supplementary Figure S54). Three methods of quantifying shared gene expression were employed - the average expression of expressologs or EnsemblPlant homolog groups (Supplementary Figures S55-56), the most abundant gene within expressologs or EnsemblPlant homolog groups (Supplementary Figures S57-58), or the expression of one-to-one expressologs or EnsemblPlant orthologs (Supplementary Figures S59-60). All approaches and datasets gave similar results with the exception of integration based on average expression, which did not recover tomato clusters. In these analyses, most cell types (trichoblast, pericycle, phloem, procambium and xylem) from each species co-clustered in broadly the same UMAP space (Supplementary Figures S53-60). In contrast, tomato atrichoblast cells co-clustered with Arabidopsis root cap cells, (Figure 5c; Supplementary Figures S53-60). Further, tomato exodermal cells, co-clustered with the responsive sub-populations of Arabidopsis cortex and endodermis (Figure 5c; Supplementary Figures S53-60). To further compare the expression profiles of N-responsive genes across cell types, we compared the expression patterns of tomato expressologs of Arabidopsis genes to their corresponding cell type expression profile in Arabidopsis (Figure 5d). Most N-responsive tomato genes maintain their expression directionality in response to N in Arabidopsis, but few show a conserved expression pattern across cell types. The expressologs of genes that are predominantly differentially expressed in tomato exodermis are differentially expressed in a range of different Arabidopsis cells, demonstrating vast rewiring of the N transcriptional cell type response.

### Differences in cell type N-transcriptional regulation are accompanied by divergent NLP7 targets

We then asked which biological processes are N-dependent in each cell type. Mapman ^40^ enrichment of differentially expressed genes indicated that the tomato exodermis shows significant enrichment of cell wall organization and N-related assimilation processes, suggesting structural remodeling is a key N response in this cell type (Supplementary Figure S61).

Given the differences in cell type-responsiveness between Arabidopsis and tomato, we next investigated whether the enrichment of N-responsive genes in the tomato exodermis could be explained by expression of *SlNLP7a* or *SlNLP7b*. In tomato, *SlNLP7a* and *SlNLP7b* are paralogs, but are not direct orthologs of Arabidopsis *AtNLP6* and *AtNLP7* ^11^. The cell type-expression patterns and N-responsiveness of *AtNLP6* and *AtNLP7* differ, as do *SlNLP7a* and *SlNLP7b* (Figure 6a; Supplementary Figure S62), and there is no clear similarity in their respective expression patterns between species. *AtNLP6* expression is enriched in the phloem, and *AtNLP7* in the pericycle, endodermis, cortex, trichoblast and root cap. *SlNLP7a* expression is enriched in pericycle and atrichoblast cells while *SlNLP7b* is enriched in the exodermis (Figure 6a). To determine the degree to which the N spatial transcriptional program is controlled by SlNLP7a and SlNLP7b, as well as its conservation and divergence from that of Arabidopsis, we sought to identify the direct targets of SlNLP7a and SlNLP7b at cellular resolution. SlNLP7a and SlNLP7b glucocorticoid receptor fusions were generated and introduced into tomato by hairy root transformation. A TARGET assay was performed in which hairy roots expressing either SlNLP7a or SlNLP7b fused to a C-terminal glucocorticoid receptor (GR) tag were exposed to four treatments - dexamethasone (DEX), controlling nuclear entry, cycloheximide (CHX) as a translational inhibitor, both DEX and CHX (CD), and a mock control (Supplementary Figure S63). Protoplasts were isolated from each treatment group from each respective hairy root line. In total, 96,304 cells were sequenced and data was processed using COPILOT and Seurat ^29,41^. In this experiment, protoplasting genes were removed and annotation was performed as described above (Supplementary Figures S64-S66; Supplementary Dataset S11). *SlNIR1* and *SlNIR2* were previously identified as targets of SlNLP7a/b ^11^ and were also identified as significantly upregulated in response to CD compared to CHX (Figure 6b; Supplementary Figure S67; Supplementary Datasets S12-S13). Data from all four treatments were integrated and direct upregulated and downregulated targets for SlNLP7a and SlNLP7b were identified for each cell type (Figure 6c; Supplementary Datasets S14-S15). The exodermis had the most downregulated targets for both SlNLP7a and SlNLP7b, while meristematic cells had the most upregulated targets (Figure 6b). Some SlNLP7a and SlNLP7b targets were also N-responsive in a cell type-specific manner (Figure 6c). Interestingly, *SlAREB1*, an ortholog of *AtABF2* (a known coordinator of endodermal N-responses), was found to be highly enriched in the tomato exodermis. While there is no single ortholog of *AtABF3* in tomato, the closest relatives were also found to be enriched in the exodermis ^42,43^ (Supplementary Figure S68).

**Figure 6.**
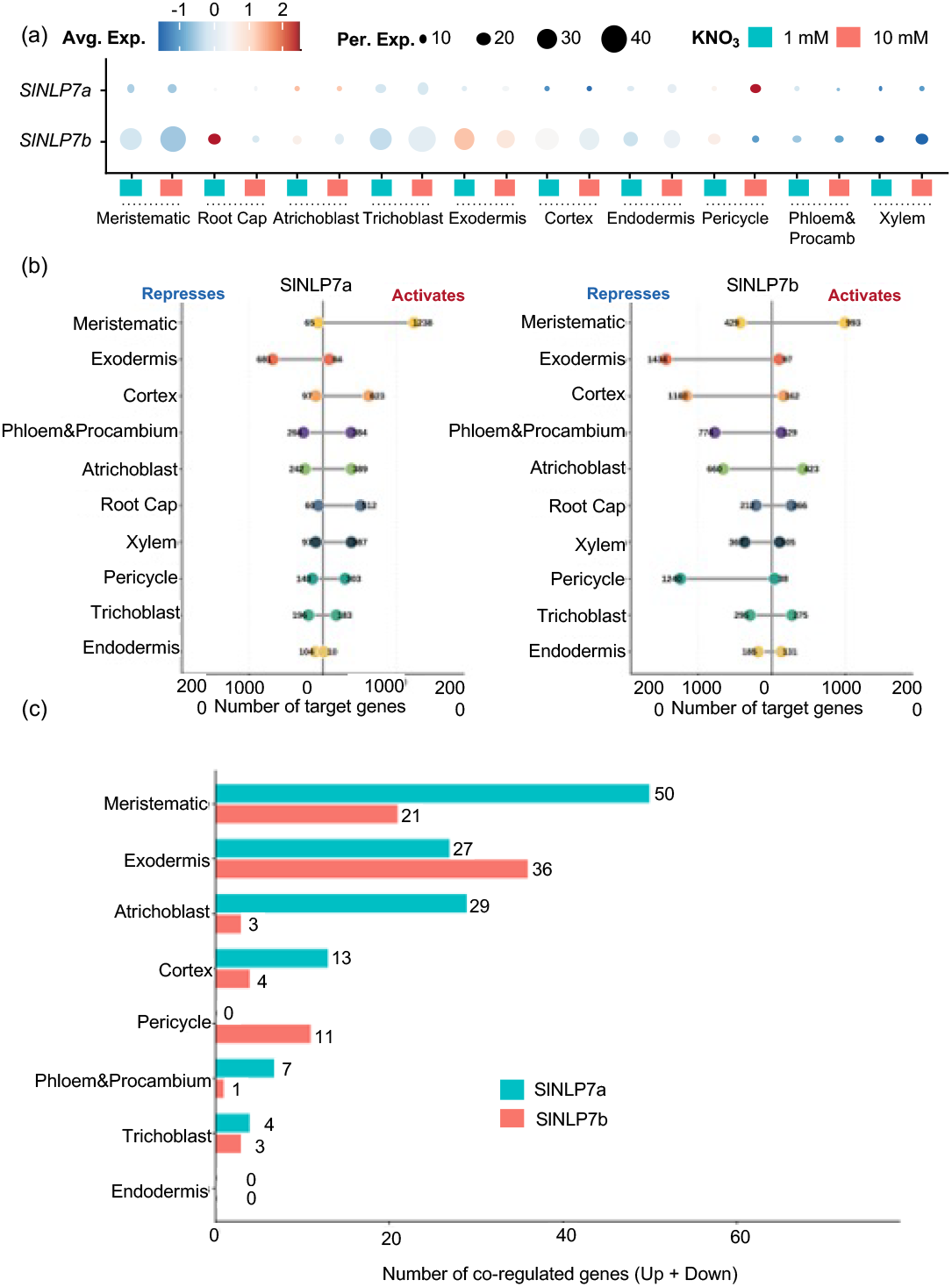
Direct targets of SlNLP7a and SlNLP7b are enriched in the exodermis. **(a)** Cell type expression profiles of SlNLP7a and SlNLP7b in limiting (1 mM) vs sufficient (10 mM) KNO_3_. **(b)** Lollipop plots showing numbers of direct targets of SINLP7a (left) and SINLP7b (right) in each cell type **(c)** Numbers of NLP7 target genes that are also nitrogen-responsive in each cell type.

### Exodermal cell wall remodeling in response to nitrogen is controlled by SlNLP7a and SlNLP7b

The tomato exodermis cell wall is remodeled throughout its development, with deposition of a polar lignin cap, followed by suberin. We further curated a gene list with specific categories of genes associated with these processes as well as N transport and signaling, and performed gene set enrichment analysis (Supplementary Dataset S16). The majority of these gene sets were significantly down-regulated by SlNLP7a and SlNLP7b (Figure 7a). We thus hypothesized that the exodermis undergoes N-responsive lignification and suberization of its cell wall. Tomato roots were exposed to 1 mM (limiting) and 10 mM (sufficient) KNO_3_ for 5 days, sectioned, cleared with ClearSee ^44^, and stained with basic fuchsin to visualize lignin, or fluorol yellow for suberin (Figure 7b, c). Lignin and suberin are both increased in limiting nitrate compared to sufficient nitrate. Given that lignin and suberin-related genes are down-regulated targets of SlNLP7a and SlNLP7b, we hypothesized that these factors repress these metabolic programs in sufficient nitrate. To investigate this further, we analyzed suberin and lignin levels in the *slnlp7a* mutant ^11^. Consistent with this hypothesis, suberin and lignin levels are high in the exodermis in both limiting and sufficient KNO_3_ (Figure 7d, e significantly).

**Figure 7.**
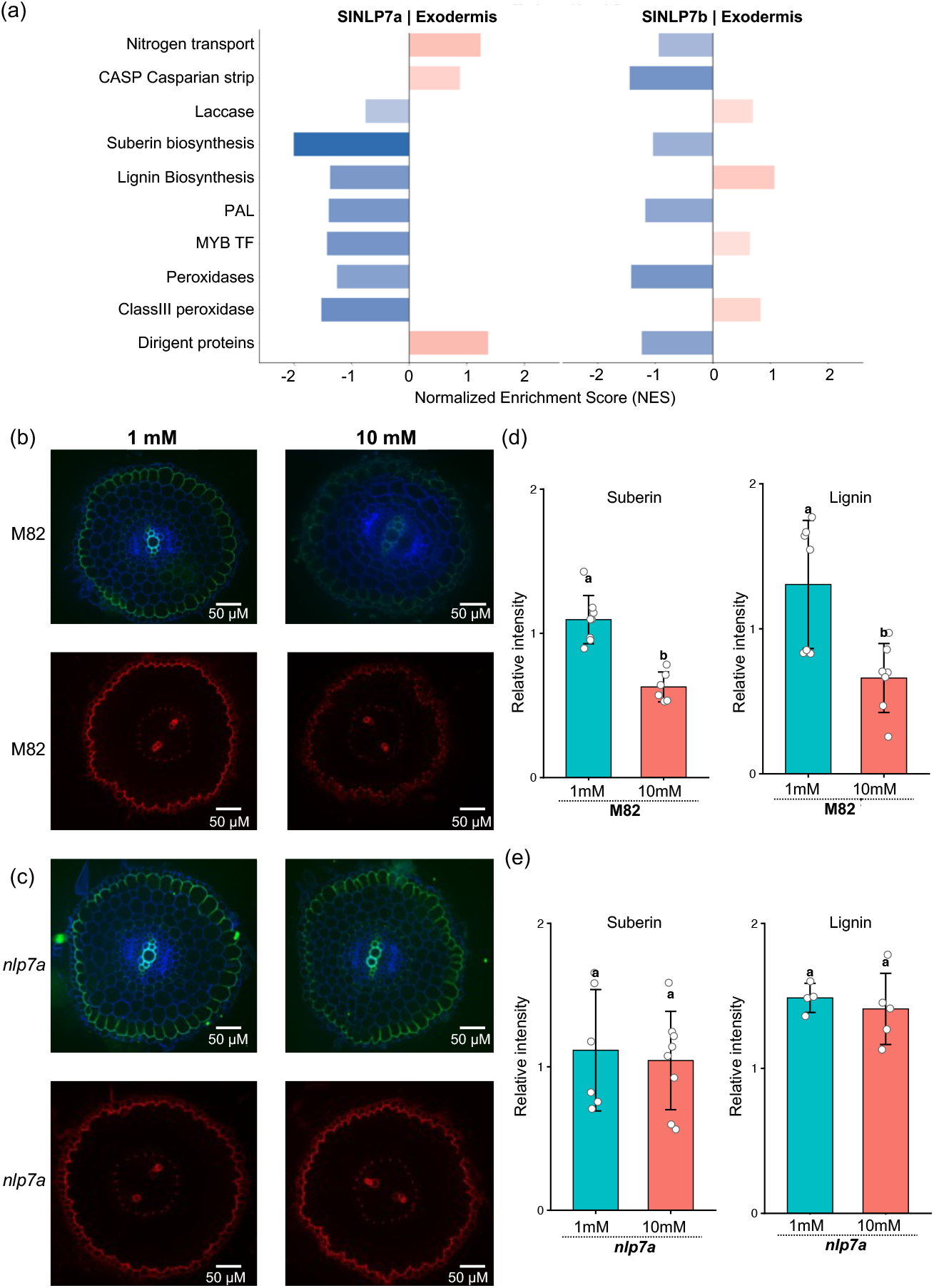
Lignin and suberin deposition in the tomato exodermis increases in limiting nitrate. **(a)** Gene set enrichment analysis of direct targets of SlNLP7a and SlNLP7b expressed in tomato exodermis **(b-c)** wild type (M82) **(b)** and *nlp7a* **(c)** tomato roots grown in 1 mM (left) and 10 mM (right) KNO_3_ stained with basic fuchsin to visualize lignin (top) and fluorol yellow to visualize suberin (below) **(d-e)** Quantification (relative intensity) of suberin and lignin in (d) wild type (M82) and (e) *nlp7a* tomato roots.

## Discussion

This study provides a high-resolution, cell-type and developmental stage-specific map of the transcriptional landscape in Arabidopsis and tomato roots grown in varying concentrations of nitrate. Using single-cell transcriptomics and cross-species integration, we have identified conserved and divergent aspects of N-responses and signaling, highlighting that while key genes coordinating nitrate response are conserved, their cellular deployment varies significantly across these lineages.

Our single-cell data provides a new level of resolution for the TF-dependent coordination of N-transcriptional responses. While some DEGs detected in the bulk transcriptome were not differentially expressed in any individual cell type, more DEGs were detected in the single cell data than in the bulk transcriptome, as expected, revealing the complexity of the N-transcriptional landscape (Supplementary Figure S7). Comparison with proteomic data collected from plants grown in the same conditions showed that changes in the abundance of some proteins correlated with changes at the transcript level (Supplementary Figure S15). Others remained unchanged, consistent with previous studies in which differences in functional proteins are not fully reflective of transcriptional changes ^17,45^. Nonetheless, the patterns observed in these datasets largely represent the true nature of N-dependent signaling.

Supporting previous literature, we found that the endodermis, pericycle and cortex were enriched in N-responsive genes and that many characterized genes involved in N-responses show cell-type enrichment (Figure 1) ^22–24^. The application of scRNAseq to whole roots identified that sub-populations of mature cortex, endodermis and pericycle cells are enriched for N-responsive genes. Interestingly, a recent single-cell study also identified sub-populations of cortex cells in several species of Brassicaceae, including Arabidopsis ^46^. Previous work with nitrate sensors showed variation in nitrate across root cell layers with endodermis cells accumulating the most nitrate when seedlings were grown under N-free conditions, the cortex showing the highest and fastest accumulation following nitrate exposure, suggesting higher rates of nitrate update and/or transport within the ground tissue ^47^. As AtNLP7 can directly sense N ^9^, activating expression of its target genes in response, the N-responsiveness of AtNLP7 target genes in the cortex is likely a product of both cell-specific enrichment of nitrate and its consequent perception and downstream transcriptional cascade

The expression pattern of *AtNLP7* in our single-cell data is consistent with previous *in situ* hybridization observations ^7^. Analysis of cell type transcriptomes in limiting and sufficient KNO_3_ revealed both shared and specific N-responsive programs (Figures 2 and 3). Notably, they reveal that NLP7 and TGA1 regulate the N-responsive expression of different sets of target genes in different cell types (Figure 3b-c). Integration of our Arabidopsis scRNAseq data with previously published TARGET gene data further revealed the pericycle to be enriched in N-responsive root development genes that are regulated by AtNLP7, consistent with known changes to lateral root number in response to nitrate ^48^. Consistent with observations that nitrate rapidly accumulates in the Arabidopsis cortex, these cells were enriched in transporters (Figure 2i and 3d) ^47^. For example, we found *AtNPF6*.*3* (*AtNRT1*.*1*) is enriched in the cortex and strongly upregulated in response to KNO_3_, which is consistent with previous studies ^49,50^ (Figure 3d). A similar expression pattern was observed for *AtNRT2*.*1* in agreement with previous studies, although that study reported repression in high nitrate ^51,52^.

Analysis of Arabidopsis *nlp7-1* and *anac032* mutant lines show that N-responsive targets of AtNLP7 retain their cell type enrichment in the absence of these TFs but their N-responses are perturbed (Figure 4). The extent of N-response perturbation appears to be dependent on whether they are co-regulated by other N-responsive transcription factors: in this study we observed that AtNLP7 target genes that are co-regulated by AtTGA1 tend to retain their N-responsiveness in *nlp7* mutants. It is likely that other target genes may be co-regulated by other N-responsive TFs, highlighting the immense complexity and robustness of this N-regulatory network. Our data provides evidence that AtNLP7 is not required for cell-type expression patterns of its target genes, which suggests that the regulatory sequences of these genes may encode separate modules for spatial expression and N-responsiveness. It remains to be observed if this is a property of transcriptional regulatory network associated with the metabolism of addition critical nutrients.

Recent work has highlighted similarities and differences in the genetic networks that underlie regulation of N-associated metabolism between Arabidopsis and tomato ^11^. A remaining gap in these networks was how these regulatory networks were deployed at spatiotemporal resolution. Differences in upstream regulation of NLP7 in tomato and Arabidopsis, coupled with enrichment of genes associated with N metabolism in an inferred tomato exodermal regulatory network suggested that spatiotemporal differences in these species’ transcriptional regulatory networks ^11,25^. In this study, we find that while many of the N-responsive genes identified in tomato show the same expression directionality in differing concentrations of N in Arabidopsis, there is vast rewiring of cell type-expression. This suggests a role for the tomato exodermis as central to N transcriptional regulation and metabolism in this species ^25,35,39^. Additional support for rewiring of the exodermis as a central hub for N metabolic regulation is the exodermal enrichment of predicted tomato orthologs of *AtABF2* and *AtABF*3, which are major hubs for nitrate responses in the Arabidopsis endodermis ^24,42,43^ (Supplementary Figure S68).

The differing expression patterns of *AtNLP6/7* and *SlNLP7a/b* (Figure 3 and Figure 6; Supplementary Figure S62) reveal the likely source for some of the events that underlie these large-scale spatial changes. The TARGET assay, which identifies target genes of TFs ^53^, was adapted to obtain cell-type data and applied to tomato to identify direct targets of SlNLP7a/b at cellular resolution (Figure 6). Both SlNLP7a/b activate expression of genes in meristematic cells, with SlNLP7a primarily a transcriptional activator in most cell types, and SlNLP7b a repressor, with the exception of both factors primarily repressing expression of their targets in the exodermis (Figure 6a). Many but not all of these targets are also N-responsive genes (Figure 6c). Biological process enrichment of these target genes revealed cell wall remodeling as a target process. Refined assessment of an annotated list of genes associated with exodermal lignification and suberization revealed that these are directly repressed targets of SlNLP7a/bSlNLP7a/b in sufficient nitrate. The deposition of suberin lamellae in the endodermis of other plant species plants is induced by deficient iron, manganese, zinc, potassium, and sulfur as well as by osmotic stress ^54,55^. In tomato, suberin deposition is increased in response to ABA and water deficit ^39^. This nutrient and abiotic-stress induction of suberin influences the intercellular transport of mineral nutrients and solutes from the endodermis to the vasculature. Our TARGET data and patterns of lignin and suberin accumulation suggest that SlNLP7a (and likely SlNLP7B) actively repress suberin deposition in sufficient nitrate. The high accumulation of suberin in limiting nitrate may preserve available N within the root.

Plant cells reinforced by a secondary cell wall can be difficult to capture with protoplasting-based methods (Pasternak et al. 2021; Tang et al. 2026). Further, removal of a cell wall can elicit responses related to cell damage and its perception. Our decision to utilize single protoplasts was predicated on the need for higher read depth per cell to capture regulatory TF hubs. Multiple approaches were also used to assess the impact of protoplasting on cell type gene expression, and these were determined to be limited, and nitrate-specific signals were robustly detectable. However, this does not preclude our ability to successfully capture cells that are highly suberized. Indeed, we only identified a small number of cells expressing genes controlling downstream steps within the suberin biosynthesis and transport pathway (Figure 7). Even with these limited cell numbers, we were able to validate N concentration-dependent changes in exodermal suberin deposition.

This work provides a comprehensive cellular resolution analysis of transcription in limiting and sufficient N in two plant species. The nitrate response is highly spatially dynamic, and enhanced in specific developmental stages. The massive changes in cell type transcriptional regulation between these two species represents the dramatic spatial rewiring of critical nutrient metabolism in between two species. Many plant species vary in cortex layer number and their functional specialization is generally unknown. These results demonstrate that different cell types can serve as nutrient regulatory hubs, and that in this case, the master transcriptional regulator controls only the N-dependent gene regulation and not its spatial context. The modes by which AtNLP6/7 and SlNLP7a/b are able to act as both activators or repressors are unknown, but are likely due to interactions with other factors. A remaining gap in our knowledge are the factors that coordinate the cell type specific transcription of N-associated metabolic and regulatory genes.

## Methods

### Plant material and growth conditions

Arabidopsis *nlp7* is a homozygous T-DNA insertion line (*nlp7-1*; SALK_26134) in a Col-0 background and was obtained from the Nottingham Arabidopsis Stock Centre ^7117^. Arabidopsis *anac032* (*anac032-cc-1*) has a homozygous, CRISPR-induced mutation and is in a Col-0 background (^11^). Arabidopsis seeds (Col-0, *nlp7* and *anac032)* were sieved to obtain seeds of 250-300 μm, surface sterilized with chlorine gas for ∼16 hours, stratified at 4°C for 3 days in saturated benomyl (methyl 1-(butylcarbamoyl)-2-benzimidazolecarbamate; Sigma-Aldrich 381586) and washed with sterile water three times. Approximately 500 surface-sterilized seeds were sown in 2 rows on 120 mm square petri dishes on top of a nylon mesh over growth media (1 mM or 10 mM KNO3; 4 mM MgSO_4_; 2 mM KH_2_PO_4_; 36 mg/L NaFe-EDTA; 1 mM CaCl_2_; 10 mM KCl; 146 mg/L MES, 5.72 mg/L H_3_BO_3_; 3.62 mg/L MnCl_2_·4H_2_O; 0.22 mg/L ZnCl_2_; 0.1 mg/L CuCl_2_·2H_2_O; 0.05 mg/L Na_2_MoO_4_·2H_2_O; 1% (w/v) sucrose; 0.75% phytagel (pH 5.7)) as previously described ^11^. *Solanum lycopersicum* (tomato) cultivar M82 seeds were surface sterilized with 70% (v/v) ethanol for 3 min, followed by three 5 min washes in sterile water, 20 minutes in 1:1 (v/v) sodium hypochlorite (Supelco, 6∼14% active chlorine) with 0.05% (v/v) Tween-20, and six 5 min washes in sterile water. Seeds were kept at 4°C for 3 days, and 10 seeds were sown as described above. Plates were placed vertically in a controlled environment chamber (22°C; 16-hour light/8-hour dark; 57 µmol/m^2^/s photosynthetic photon flux density (PPFD)).

### Sample preparation and data processing for bulk RNA-seq

Whole roots from 12-14 days old seedlings grown as described above were harvested and snap-frozen in liquid N2. Roots from all seedlings (∼ 500 Arabidopsis or 10 tomato seedlings) grown on a single plate were pooled to form one biological replicate. Two biological replicates were prepared for each species at each nitrate concentration. Total RNA was extracted from Arabidopsis roots using the Spectrum Plant Total RNA Kit (Sigma-Aldrich, STRN250) and from tomato roots using the Vazyme FastPure Universal Plant Total RNA isolation kit (Vazyme, RC411-01), with on-column gDNA digestion using the QIAGEN RNase-free DNase Set (QIAGEN, 79254). RNA-seq libraries were constructed using the NEBNext Ultra II RNA Library prep for Illumina kit (New England Biolabs, E7760), NEBNext Poly(A) mRNA Magnetic Isolation Module (NEB#7490) and NEBNext Multiplex Oligos for Illumina® (96 Unique Dual Index Primer Pairs) (E6440S) at a concentration of 10 µM. Libraries were sequenced on the NovaSeq X Plus 10B (tomato) or NovaSeq 6000 SP (Arabidopsis) flow cell with 150bp PE reads. Reads were quantified using Salmon (v1.10.0) ^56^ against the reference transcriptomes of *Arabidopsis thaliana* (TAIR10, Ensembl Plants release 52) and *Solanum lycopersicum* (ITAG4.0 with appended organellar sequences as described below). Salmon indices were built with a k-mer size of 31 and the corresponding genomic sequences as the decoys. Paired-end reads were quantified using the Salmon quant command with the -- validateMappings option enabled. Gene-level counts were summarized using the tximport (v1.32.0) ^57^ and further processed using DESeq2 ^58^.

### Sample preparation, LC-MS and data processing for bulk proteomics

Whole roots from 11-day old Col-0 seedlings grown as described above were harvested, snap-frozen in liquid N2, and freeze-dried for 4 days at -52°C, 90 mTorr using the VirTis Benchtop Freeze Dryer. 5∼30 mg (dry weight) of freeze-dried material was homogenized using 3 mm tungsten carbide beads with a TissueLyser II at 30 Hz for 3 min (Qiagen, Hilden, Germany). Roots from all seedlings grown on a single plate were pooled to form one biological replicate. Four biological replicates were prepared for seedlings grown under each nitrate concentration. Ground tissue was then extracted, digested and desalted as previously described, without deviation ^59^.

Peptides were then analyzed on a Fusion Lumos Orbitrap mass spectrometer (Thermo Fisher Scientific) using DIA mode, where a total of 2 μg of resuspended peptide was injected per replicate using Easy-nLC 1200 system (LC140; Thermo Fisher Scientific) and an Acclaim PepMap 100 C18 trap column (Cat# 164750; Thermo Fisher Scientific) followed by a 50 cm Easy-Spray PepMap C18 analytical column (ES903; Thermo Fisher Scientific) warmed to 50 °C. Peptides were eluted at 0.3 μl/min using a segmented solvent B gradient of 0.1% (v/v) formic acid in 80% (v/v) acetonitrile (A998, Thermo Fisher Scientific) from 4 to 41% solvent B (0–85 min). FAIMSpro gas flow was fixed to 3.5 L/min using CV settings -30, -50 and -70 ^59^. A positive ion spray voltage of 2.3 kV was used with an ion transfer tube temperature of 300 °C and an RF lens setting of 40%.

A BoxCar data independent acquisition (DIA) LC-MS acquisition approach was used to acquire all LC-MS data ^60^. MS1 spectra were acquired using two multiplexed targeted SIM scans of 10 BoxCar windows each. Full scan MS1 spectra (350–1400 m/z) were acquired with a resolution of 120,000 at 200 m/z and normalized AGC targets of 100% per BoxCar isolation window. Fragment spectra were acquired at resolution of 30,000 across 28 total 38.5 m/z windows overlapping by 1 m/z using a dynamic maximum injection time and an AGC target value of 2000%, with a minimum number of desired points across each peak set to 6, as previously described ^60^. Higher-energy collisional dissociation (HCD) fragmentation was performed using a fixed 27% fragmentation energy.

BoxCarDIA LC-MS data analysis was performed using Spectronaut ver. 18 (Biognosys AG) with default settings. All searches made against a custom-made decoy (reversed) version of the Arabidopsis protein database from Araport 11 (27,533 protein encoding genes; ver. 2022-09-14). Search parameters included the following: trypsin digest permitting two missed cleavages, fixed modifications (carbamidomethyl (C)), variable modifications (oxidation (M)) and a peptide spectrum match, peptide and protein false discovery threshold of 0.01. Differences in protein abundance between KNO_3_ conditions were assessed using the Wilcoxon rank-sum test on MS2 quantification values.

### Protoplast isolation and library preparation for scRNA-seq

Whole roots from pools of 10 day-old tomato and Arabidopsis seedlings grown as described above were harvested and protoplasts were isolated as previously described ^11,39^ with minor modifications. In brief, roots were cut into 1 cm pieces and covered with digestion buffer (0.6 M mannitol, 20 mM KCl, 20 mM MES pH 5.6, 10 mM CaCl_2_, 0.1% BSA supplemented with 1.5% cellulase R10 (Duchefa C8001) and 0.15% pectolyase (Sigma-Aldrich P3026) (Arabidopsis), or 1.25% cellulase R10, 1.25% cellulase RS (Duchefa C8003), 0.3% macerozyme R10 (Duchefa M8002) and 0.12% pectolyase (tomato)). After 1.5 h in the dark with gentle shaking, the suspension was filtered through stacked 70 μm and 40 μm cell strainers and cells were collected into 50 mL centrifuge tubes on ice. Cells were collected by centrifugation for 5 min (400 x *g*, 4°C) with centrifuge acceleration and deceleration set to minimum. Collected protoplasts were washed with 15 mL ice cold wash solution (WS) (digestion buffer without enzymes) centrifuged for 5 min (350 x *g*; at 4°C) and resuspended in 1 mL ice cold WS. Cells were filtered through a 70 μm Flowmi strainer (Bel-Art H13680-0070) and transferred to 2 mL microcentrifuge tubes. Viability and cell density was determined by mixing cells with an equal volume of 50 μg/mL fluorescein diacetate (FDA) and imaged using an EVOS fluorescence microscope (ThermoFisher) with excitation 470/22 nm and emission 525/50 nm. Protoplasts were centrifuged for 5 mins (350 x *g*; at 4°C) and resuspended to 1000-2000 cells/μL before a final filtration through a 70 μm Flowmi strainer. Cells were loaded onto a Chromium Next GEM Chip G (10X Genomics) at a density that aimed to recover 10,000-16,000 cells per sample, and single-cell libraries were generated using the Chromium Next GEM Single Cell 3’ Kit v3.1 (10x Genomics PN-1000268) according to the manufacturer’s instructions. Libraries were pooled and sequenced to a target depth of 50,000 reads per cell with a 300 cycle NovaSeq X Series 10B Reagent Kit (Illumina 20085594) on the NovaSeq X Plus instrument. The pooled library was diluted to 0.7 nM using EB (10mM Tris pH8.0) in a volume of 34 µl per lane before spiking in 1% Illumina PhiX Control v3 (Illumina FC-110-3001). This was denatured by adding 8.5 µl 0.2 N NaOH and incubating at room temperature for 5 mins, neutralized by adding 127.5 µl of Illumina preload buffer. 165 µl was loaded onto the sequencing cartridge flow cell. Libraries were sequenced on a NovaSeq X sequencer (Illumina, San Diego, CA) with paired-end 150 bp reads. The data was base masked to 28bp (read 1) and 90bp (read 2), demultiplexed, and converted to fastq using bcl2fastq2 version 2.20.

### Modified TARGET assay for scRNA-seq

A variation of the TARGET assay ^11,53^was performed using protoplasts of transgenic hairy roots followed by scRNA seq. Tomato hairy root transformation was performed following the protocol described by ^61^. Briefly, competent *Rhizobium rhizogenes* (ATCC15834) cells were transformed by electroporation with plasmids harboring SlNLP7a or SlNLP6b translationally fused to a C-terminal glucocorticoid receptor (GR) tag and expressed from the cauliflower mosaic virus 35S promoter ^11^. A single colony was used to inoculate liquid cultures, which were incubated overnight and used to transform tomato cotyledons. For each construct, 20 fully expanded cotyledons were excised and immersed in *R. rhizogenes* culture and cultivated in the dark for three days on MS agar without antibiotics. On the third day, infected cotyledons were transferred to fresh MS agar supplemented with 200 mg/L cefotaxime and 50 mg/L kanamycin for selection. At least 20 independent antibiotic-resistant roots were subcloned per construct for further analysis. To identify stably transformed lines, three newly emerged roots were harvested from each of the 20 subcloned roots and the presence and expression of the transgene was analyzed by qPCR. Lines exhibiting consistent and uniform transgene expression levels across the newly grown roots were identified, and the most stable line was selected for subsequent experiments.

Protoplasts were isolated from transformed hairy roots as described by ^62^ with modifications. Briefly, roots of two-week-old subcloned lines were finely chopped and incubated for 4 hours at 28°C in an enzyme solution containing 1.5% cellulase R10, 1.5% cellulase RS, 0.4% macerozyme, and 0.13% pectolyase. Following incubation, the protoplast suspension was sequentially filtered through 70 µm and 40 µm strainers to remove debris. Protoplasts were pelleted by centrifugation at 500 × g for 5 minutes at 4°C and washed twice with 12 ml of ice cold W5 solution (2 mM MES, 154 mM NaCl, 125 mM CaCl_2_, 5 mM KCl, pH 5.7) and and resuspended in WI solution (0.2 M MES pH 5.7, 0.8 M mannitol, 2 mM KCl) in a round-bottom microcentrifuge tube. Protoplasts were divided into four treatment groups as follows: (1) 10 µM dexamethasone (in 100% ethanol) for 3 hours; (2) 35 µM cycloheximide (in DMSO) for 20 minutes; (3) sequential treatment with 10 µM dexamethasone followed by 35 µM cycloheximide; or (4) solvent control (DMSO and ethanol). Following treatment, protoplasts were pelleted, resuspended in WI solution, and counted prior to library preparation and sequencing. For library preparation barcoded 3’ single cell libraries were prepared from single cell suspensions using the Chromium GEM-X Single Cell 3’ kit v4 (10X Genomics, Pleasanton, California) for sequencing according to the recommendations of the manufacturer. The cDNA and library fragment size distribution were verified on a Bioanalyzer 2100 (Agilent, Santa Clara, CA). The libraries were quantified by fluorometry on a Qubit instrument (LifeTechnologies, Carlsbad, CA) and by qPCR with a Kapa Library Quant kit (Kapa Biosystems-Roche) prior to sequencing. The libraries were sequenced as described above.

### Pre-processing of scRNA-seq data

Sequencing reads obtained from scRNAseq were mapped to the respective reference genomes using CellRanger (v8.0.1, 10x Genomics). For Arabidopsis, the TAIR10 reference genome and the corresponding gene annotation (Ensembl Plants release 52) were used to build the CellRanger reference using cellranger mkref. For tomato, the SL4.0 reference genome and the corresponding gene annotation ITAG4.0 with extended 3’ UTR annotations and appended organellar (mitochondrion and plastid) sequences were used to build CellRanger reference: Organellar (mitochondrial and plastid) genomes from SL3.1 assembly and ITAG3.1 annotation were appended to the SL4.0 genome and ITAG4.0 annotation, respectively, to generate SL4.0-ORG and ITAG4.0-ORG references. To improve the mapping of scRNA-seq reads from tomato KNO_3_ treatment samples, 3’UTR annotations in ITAG4.0-ORG were extended using peaks2utr (v1.4.1) based on the scRNA-seq data generated in this study ^63^. The resulting annotation file was used to build the final CellRanger reference.

For scRNAseq datasets from protoplasts isolated from roots grown in different KNO_3_ treatments, gene expression matrices were generated using cellranger count and downstream analysis was performed using Seurat (v5) in R (v4) ^64^. Cells with a minimum of 300 UMI counts (nCount_RNA), between 300 and 10,000 detected genes (nFeature_RNA), and less than 10% (Arabidopsis) or 30% (tomato) organellar reads were retained for downstream analysis. Mitochondrial and plastid genes were removed from the gene expression matrix. Genes affected by protoplasting (q_value < 0.05 & absolute Log2 fold change > 2) ^37,39^ were either retained or removed as described in the results. Doublets (emulsion droplets containing two or more cells) were predicted using the DoubletFinder package (v2.0.6) ^65^. The expected multiplet rate was estimated based on the number of input cells using a linear regression model derived from estimates provided by the user guide of Chromium Next GEM Single Cell 3′ Reagent Kits v3.1 (Dual Index) (10x Genomics, https://cdn.10xgenomics.com/image/upload/v1722285481/support-documents/CG000315_ChromiumNextGEMSingleCell3_GeneExpression_v3.1_DualIndex_RevF.pdf).

Samples were integrated using the IntegrateLayers function of Seurat (method = CCAIntegration, normalization.method = “SCT”). Following integration, clustering and UMAP non-linear dimensional reduction were carried out using the first 30 principal components at a resolution of 0.4 (Arabidopsis) or 1.0 (tomato).

For scRNAseq datasets obtained from TARGET assays, high ribosomal RNA (rRNA) content was observed in the scRNAseq libraries generated from the Chromium GEM-X Single Cell 3’ Kit v4 (10x Genomics). To address this, low quality cells were filtered before Seurat analysis. Raw counts from CellRanger were processed using COPILOT ^29,41^. Barcodes with fewer than 500 UMIs were removed as low quality. The remaining barcodes were separated into high and low quality groups using a data-driven UMI threshold, with a minimum of 800 UMIs required for a cell to be considered high quality. Cells were further classified as a signal or noise by comparing their log normalized expression profiles to population level reference profiles using Pearson correlation, where cells correlating strongly with the noise profile were excluded. COPILOT filtered count matrices were loaded into Seurat (v5) for downstream analysis. Genes affected by protoplasting (q_value < 0.05 & absolute Log2 fold change > 2) ^39^ were removed. Doublets were identified and removed using scDblFinder ^66^. Batch correction across samples was performed using Harmony ^67^, integrating over sample identity using PCA embeddings as input. The top 15 Harmony-corrected principal components were used for neighbourhood graph construction and UMAP dimensionality reduction (n.neighbors = 40, min.dist = 0.08). DMSO samples (DM_X and DM_1) were subset from the integrated object without re-clustering, retaining the original cluster identities for downstream analysis.

### Annotation of Arabidopsis scRNAseq dataset

Annotation of Arabidopsis scRNAseq clusters to the corresponding cell types and developmental stages was performed using a combination of techniques. First, a previously published Arabidopsis root atlas (^29^ was reannotated for developmental stages as described by ^33^ and used as a reference for label transfer using the TransferData function in Seurat. Anchors between the reference and query datasets were identified using the FindTransferAnchors function (normalization.method = “SCT”, reference.reduction = “pca”). Cell-type and developmental stage labels were then transferred from the reference to the query dataset using the TransferData function with default parameters. Second, index of cell identity (ICI) scores were used to estimate cell identities based on a matrix of cell-type specific gene expression data ^36,68^. Briefly, specificity score tables for cell identities and developmental stages were constructed using existing cell-type/stage specific expression datasets from Arabidopsis curated in ^29^. The SCT normalized expression data were then used to compute ICI scores of cell identities and developmental stages for each cell. Cells with significant ICI scores (p_adj < 0.1) were assigned the corresponding cell identity or developmental stage with the highest ICI score. For clusters in which the majority of cells shared the same label transfer and ICI prediction, this cell type was assigned. Columella, lateral root cap (LRC) and quiescent center (QC) cells were combined and classified as root cap (RC).

Finally, annotations were refined based on developmental stage predictions and cluster marker genes as follows: 1) Cells in cluster 16 were predicted to be of mixed cell types in proliferative cell stages, and were annotated as meristematic (Supplementary Figure S4); 2) Cells in cluster 19 were predicted to be proximal lateral root cap and cells of mixed cell identity in the proliferation domain, and were annotated as meristematic (Supplementary Figure S5); 3) Cells in cluster 28 were predicted to be atrichoblast cells in the proliferation domain and proximal lateral root cap, were annotated as meristematic cells (Supplementary Figure S5); 4) Cells in cluster 21 contained a mix of cells with labels of procambium, trichoblast and pericycle. Marker genes enriched in this cluster were calculated using the FindConservedMarkers function (grouping.var = “sample”, only.pos = FALSE, assay = “RNA”,slot = “data”). Among the highly enriched markers in the cluster, *AtNPF7*.*3* (*AT1G32450*) is a transmembrane nitrate transporter known to be expressed in root pericycle cells under the control of *AtMYB59* (also enriched in cluster 21) ^69,70^. Both genes showed specific expressions in cluster 21. Therefore, cluster 21 was annotated as pericycle (Supplementary Figure S5).

### Annotation of tomato scRNAseq datasets

Similar to Arabidopsis, cell clusters from tomato scRNAseq data obtained from roots grown in different KNO_3_ conditions were annotated both by label transfer using a previously annotated tomato root atlas as a reference ^39^. The expression profiles of cluster markers was checked against the cell-type-enriched transcriptomes generated by FACS sorting ^25^. For clusters in which the majority of cells shared the same label transfer or the cluster markers were enriched in the same cell type, this cell type was assigned. Phloem and procambium cells were not separated due to lack of reference markers and dataset. Only markers specific to exodermis or inner cortex, not generic cortex markers, were used to distinguish between exodermis and inner cortex (annotated as Cortex). Annotations were refined for the following clusters: 1) Cells in cluster 15 showed mostly cortex labels (mostly inner cortex) according to label transfer and MapQuery results. A large number of cluster markers was shown to be enriched in exodermis than inner cortex. Exodermis-specific genes that were known to be mostly expressed in mature exodermis were enriched in cluster 15 ^39^. So cluster 15 is annotated as exodermis (Supplementary Figure S47). 2) Cells in cluster 16 showed mostly phloem&procambium labels, whereas top cluster markers were mostly expressed in the exodermis. Examination of *CAPSP1-3* expression, known to be enriched in the endodermis ^71^, showed enrichment in the clearly identified endodermis cluster 24 as well as the cluster 16 (Supplementary Figure S47). 3) Cells in cluster 26 showed trichoblast and meristematic labels. Cluster markers were mostly expressed in epidermis. Therefore, cluster 26 was annotated as epidermis. 4) Cells in cluster 27 showed atrichoblast and meristematic labels. Only one marker in that cluster was found in the cell-type-enriched transcriptome dataset ^25^ and it was enriched in the quiescence center. Therefore, cluster 27 was annotated as meristematic.

Cell type annotation of scRNAseq data from TARGET assays was performed using label transfer and cluster marker gene expression as described above (Supplementary Figure S64a-d). Final cell type assignments based on consensus between both approaches yielded 10 cell types across 24 clusters. Annotation was validated by the consistency of label transfer predictions within each manually annotated cell type (Supplementary Figure S64e), by expression of four cross-validated marker genes per cell type (Supplementary Figure S65).

### Calculation of developmental potentials

The developmental potential of cells was estimated using CytoTRACE (v0.3.3) ^38^. CytoTRACE scores were calculated using the CytoTRACE function on normalized expression data (using the “data” layer of the “RNA” slot) under fast-mode. Cells expressing less than 200 genes and genes expressed in less than five cells were filtered out before analysis. Datasets from each genotype were processed separately and the resulting CytoTRACE scores were added to the Seurat object as metadata for further analysis and visualization.

### Differential expression analysis

Differential expression analysis between nitrate conditions was performed using two methods: 1) Single-cell level DE analysis using the ‘FindMarkers’ function in Seurat, and 2) Pseudobulk DE analysis where cells in each cell type were aggregated into pseudobulk objects for each sample, and DE analysis was performed on these pseudobulk objects using DESeq2 ^58^. The results from both methods were integrated and cross-referenced to obtain differentially expressed genes (DEGs) for each cell type. For the single-cell level DE analysis, the ‘FindMarkers’ function was used with cells from high and low nitrate conditions as the two groups for comparison, and the default Wilcoxon Rank Sum test was applied. For the pseudobulk DE analysis, cells from each cell type were aggregated into pseudobulk objects for each sample using the ‘AggregateExpression’ function in Seurat, and DE analysis was performed on these pseudobulk objects using DESeq2. Genes with an adjusted p-value < 0.05 and an absolute log2 fold change > 1 were considered as DEGs in both methods if not specified otherwise. DEGs obtained from the single-cell level DE analysis were used for further analysis by default, while the DEGs from the pseudobulk DE analysis were used to cross-reference and filter the DEGs from the single-cell level analysis.

### Integration of Arabidopsis and tomato scRNAseq datasets

Arabidopsis and tomato single-cell transcriptomes were integrated using homologous genes to define a shared cross-species feature space. Homology relationships were obtained from two sources: (1) orthologs from EnsemblPlants (release 62) and (2) tomato– Arabidopsis expressolog relationships described by Kajala et al. (2021). For the EnsemblPlants dataset, homolog groups were defined as connected components of an orthology graph, allowing orthologs with many-to-many relationships to be represented in a single group. For the expressolog dataset, each tomato gene and its associated Arabidopsis expressolog(s) were treated as a homolog group.

Four strategies were used to construct shared features from homolog groups. First, expression values (using the “counts” layer of the “RNA” assay) of all genes within a homolog group were aggregated to generate a single feature per group. Second, the average expression of all genes within a homolog group was used. Third, the expression of the most highly expressed gene within each homolog group was retained. Fourth, only one-to-one ortholog pairs were used, with each ortholog pair treated as a shared feature.

For each strategy, species-specific expression matrices were transformed into a homolog-group feature matrix, and those from two species were merged into a common matrix. The resulting matrices were processed using Seurat as described above, enabling direct comparison of Arabidopsis and tomato single-cell transcriptomes in a shared transcriptional space.

### Identification of NLP7 target genes by cell type

To identify NLP7 target genes in a cell-type-specific manner, differential expression was performed separately for *SlNLP7a* and *NLP7b* within each cell type using a Fisher exact test. Direct targets were identified by comparing CD (dexamethasone + cycloheximide) versus CHX (cycloheximide only) cells, and indirect targets by comparing DEX versus DM cells. Genes were considered differentially expressed at |log2FC| > 0.20, p_adj < 0.05, expressed in at least 2% of cells. NLP7 target genes were then cross-referenced against N-responsive gene lists from tomato bulk RNA-seq and scRNA-seq datasets (Supplementary Datasets S8-S9) and co-regulated genes (regulated in the same direction by both NLP7 and N) were counted per cell type and visualized as bar plots.

### Pathway enrichment analysis

Gene set enrichment analysis (GSEA) was performed in exodermis using the fgsea package, with genes ranked by log2FC (CD versus CHX). Curated gene sets included CASP/Casparian strip, suberin biosynthesis, lignin biosynthesis, phenylpropanoid pathway, peroxidases, laccases, dirigent proteins, MYB transcription factors and N-responsive genes. Results were visualized as NES bar plots.

### Root sectioning and staining

For suberin and lignin staining, 5-day-old wild type (M82) and *nlp7a* mutant tomato roots were sectioned using a vibratome (Leica VT1000 S). Root segments were embedded in 4% agarose and cut into 170 µm sections. For suberin, sections were taken 1 cm below the hypocotyl junction; for lignin, sections were taken 1 cm above the root tip. To visualize suberin, sections were incubated in fluorol yellow 088 (0.01% w/v in lactic acid) for 1 h at room temperature in the dark, rinsed three times with water and counterstained with aniline blue (0.5% w/v in water) for 1 h in the dark. To visualize lignin, sections were stained in ClearSee solution ^72^ containing basic fuchsin for 30 min, followed by two washes in ClearSee. Confocal imaging was performed on an Olympus IX83 inverted microscope equipped with a cicero spinning disk confocal unit (50 µm pinhole diameter, 250 µm disk pitch) and a hamamatsu ORCA flas LT3 sCMOS camera (6.5 µm pixel pitch; effective pixel size of 0.325 µm at 20X), using a ×20 objective (NA-0.80) with the following settings: 488 nm excitation and 500–550 nm emission for fluorol yellow; 550–561 nm excitation and 570–650 nm detection for basic fuchsin; and 405 nm excitation and 425–475 nm detection for calcofluor white.

## Data Availability Statement

Sequencing data is available at the European Nucleotide Archive under Project numbers PRJEB113797 and PRJEB114121. Raw proteomics data have been deposited at the ProteomeExchange Consortium (http://proteomecentral.proteomexchange.org) via the PRoteomics IDEntification Database (PRIDE; https://www.ebi.ac.uk/pride/) partner repository with the data set identifiers PXD-(to be updated).

Detailed documentation of analytical pipelines are available at Zenodo: doi:10.5281/zenodo.20342388

Code used for analysis is available at https://github.com/dizhengao/cell-type-specific-plant-nitrate-response

All other data is provided in the Supplementary Materials.

## Conflict of Interest Statement

The authors have no conflicts of interests to declare. The funders had no role in the study design, data collection and analysis, decision to publish, or preparation of the manuscript.

## Author Contributions

Conceptualization - NJP, SMB, Y-MC, ZD, IM, WH; Data curation - ZD, DeS, DaS, YL, RGU; Formal analysis - ZD, DeS, NJP, SMB, RGU; Funding acquisition - NJP, SMB, WH, IM, RGU; Investigation - ZD, DeS, NJP, SB, MT, RGU, SH, FF, TB; Methodology - ZD, DeS, Y-MC, AL, MT, RGU, ThB; Project administration - NJP, SB, WH, IM,; Resources - ZD, DS, CB; Supervision - NJP, SB, IM, WH, LC; Validation - ZD; Visualization - NJP, SB, ZD, DeS; Writing (original draft, review and editing) - NJP, SMB, ZD, DeS.

## Acknowledgements

We thank Tufan Oz for genotyping the Arabidopsis mutant lines, Dave Wright for advice on scRNAseq analysis, Chris Watkins and Karim Gharbi for administration and supervision of the sequencing project, Tonni Grube Andersen for insights and discussions on endodermis function and suberin biosynthesis, and the Plant Cell Atlas community, for discussions and advice on cell-type annotation and analytical methods. Root cell diagrams were adapted from templates shared by Frédéric Bouché under a CC-BY license (https://doi.org/10.6084/m9.figshare.468875).

## Funding

The authors acknowledge funding from the Biotechnology and Biological Sciences Research Council (BBSRC), part of UK Research and Innovation. This research was funded by the Earlham Institute Strategic Programme Grant on Cellular Genomics (BBX011070/1), and its constituent work package (BBS/E/ER/230001C), supported by Core Capability Grants BB/CCG1720/1 and BB/CCG2220/1. Additional funding was received from a joint USDA/NSF/BBSRC Breakthrough Technologies Award, Grant numbers: USDA: 2019-67013-29012, and BBSRC: BB/S020853/1, BB/Z517082/1, USDA 1032060, NSF 2516703, and the Howard Hughes Medical Institute. Part of this work was delivered by the BBSRC National Capability in Genomics and Single Cell Analysis (BBS/E/T/000PR9816) and by Transformative Genomics, the BBSRC funded National Bioscience Research Infrastructure (BBS/E/ER/23NB0006) at Earlham Institute by members of the Technical Genomics and Core Bioinformatics Groups.

## Supplementary Materials

### Supplementary Datasets

**Supplementary Dataset S1**. Cell Ranger reports of Arabidopsis scRNAseq samples generated in this study.

**Supplementary Dataset S2**. Curated lists of known Arabidopsis and tomato cell type and developmental markers.

**Supplementary Dataset S3**. Lists of differentially expressed genes in different nitrate conditions from the bulk and pseudobulk analysis of Arabidopsis root.

**Supplementary Dataset S4**. Lists of differentially expressed genes in different nitrate conditions in each Arabidopsis cell type.

**Supplementary Dataset S5**. Proteomics quantification of protein abundances in different nitrate conditions of Arabidopsis Col-0 roots.

**Supplementary Dataset S6**. Lists of differentially expressed genes between responsive and non-responsive sub-populations of Arabidopsis Col-0 cortex, endodermis, pericycle, procambium and phloem.

**Supplementary Dataset S7**. Cell Ranger reports of Tomato scRNAseq samples generated in this study.

**Supplementary Dataset S8:** Lists of differentially expressed genes in different nitrate conditions in each Tomato cell type.

**Supplementary Dataset S9**. Lists of differentially expressed genes in different nitrate conditions from the bulk RNAseq analysis of tomato roots.

**Supplementary Dataset S10**. Lists of differentially expressed genes between responsive and non-responsive sub-populations of tomato endodermis and atrichoblast.

**Supplementary Dataset S11**. Cell Ranger reports of tomato scRNAseq samples generated from the modified TARGET assay.

**Supplementary Dataset S12**. Direct SlNLP7a/b targets from pseudobulk analysis of tomato scRNAseq of the modified TARGET assay.

**Supplementary Dataset S13**. Indirect SlNLP7a/b targets from pseudobulk analysis of tomato scRNAseq of the modified TARGET assay.

**Supplementary Dataset S14**. Direct SlNLP7a/b targets in each tomato root cell type from tomato scRNAseq of the modified TARGET assay.

**Supplementary Dataset S15**. Indirect SlNLP7a/b targets in each tomato root cell type from tomato scRNAseq of the modified TARGET assay.

**Supplementary Dataset S16**. Gene set enrichment analysis of specific pathways in tomato.

### Supplementary Figures

**Supplementary Figure S1**. Experimental workflow for single-cell transcriptomics of Arabidopsis and tomato protoplasts.

**Supplementary Figure S2**. Cell and read distributions of Arabidopsis scRNAseq samples and filtering cutoffs.

**Supplementary Figure S3**. UMAP visualisation of Arabidopsis scRNAseq from different annotation methods.

**Supplementary Figure S4**. Cell type and cell stage predictions of Arabidopsis scRNAseq dataset from different annotation methods

**Supplementary Figure S5**. Refinement of annotation of Arabidopsis clusters 19, 21 and 28.

**Supplementary Figure S6**. Validation of Arabidopsis cell type and stage annotation using marker genes with known expression patterns.

**Supplementary Figure S7**. scRNAseq of Arabidopsis root protoplasts captures similar nitrate responses as RNAseq from RNA extracted from whole roots.

**Supplementary Figure S8**. Clustering and annotation of Arabidopsis scRNAseq samples with and without protoplasting-induced genes.

**Supplementary Figure S9**. Cell type abundance in the Arabidopsis Col-0 scRNAseq samples.

**Supplementary Figure S10**. Pearson correlations between high and low nitrate pseudobulk transcriptomes across Arabidopsis root cell types.

**Supplementary Figure S11**. GO terms enriched in the DEGs up-regulated under high nitrate in Arabidopsis root cell types.

**Supplementary Figure S12**. GO terms enriched in the DEGs down-regulated under high nitrate in Arabidopsis root cell types.

**Supplementary Figure S13**. Changes in the expression of nitrogen transport and assimilation related genes under different nitrate conditions in bulk RNAseq of Arabidopsis root.

**Supplementary Figure S14**. Changes in the expression of nitrogen transport and assimilation related genes under different nitrate conditions in scRNAseq of Arabidopsis root cell types.

**Supplementary Figure S15**. Protein abundance of nitrogen transport and assimilation related genes under different nitrate conditions in the Arabidopsis root.

**Supplementary Figure S16**. Comparison with previous Arabidopsis cell-type-specific nitrogen response studies.

**Supplementary Figure S17**. Comparison between N-responsive genes and DEGs from Arabidopsis bulk RNAseq.

**Supplementary Figure S18**. Developmental potential estimation of Arabidopsis root cells using CytoTRACE.

**Supplementary Figure S19**. Sub-population analysis of Arabidopsis cortex in the Col-0 WT.

**Supplementary Figure S20**. Sub-population analysis of Arabidopsis endodermis in the Col-0 WT.

**Supplementary Figure S21**. Sub-population analysis of Arabidopsis pericycle in the Col-0 WT.

**Supplementary Figure S22**. Sub-population analysis of Arabidopsis procambium in the Col-0 WT.

**Supplementary Figure S23**. Sub-population analysis of Arabidopsis phloem in the Col-0 WT.

**Supplementary Figure S24**. Overlaps of DEGs up-regulated in the responsive sub-populations of Arabidopsis cortex, endodermis and pericycle.

**Supplementary Figure S25**. Number of DEGs that are targets of NLP7/TGA1/TGA4 in each cell type of Arabidopsis Col0.

**Supplementary Figure S26**. Expression of AtABFs and their target genes in Arabidopsis root cell types.

**Supplementary Figure S27**. Expression of AtNLP7 targets across all Arabidopsis cell types.

**Supplementary Figure S28**. Cell type abundance in the Arabidopsis mutant scRNAseq samples.

**Supplementary Figure S29**. Distribution of samples in Arabidopsis scRNAseq clusters and annotated cell types.

**Supplementary Figure S30**. Sub-population analysis of Arabidopsis cortex in the *nlp7* mutant.

**Supplementary Figure S31**. Sub-population analysis of Arabidopsis endodermis in the *nlp7* mutant.

**Supplementary Figure S32**. Sub-population analysis of Arabidopsis pericycle in the *nlp7* mutant.

**Supplementary Figure S33**. Sub-population analysis of Arabidopsis procambium in the *nlp7* mutant.

**Supplementary Figure S34**. Sub-population analysis of Arabidopsis phloem in the *nlp7* mutant.

**Supplementary Figure S35**. Sub-population analysis of Arabidopsis cortex in the *anac032* mutant.

**Supplementary Figure S36**. Sub-population analysis of Arabidopsis endodermis in the *anac032* mutant.

**Supplementary Figure S37**. Sub-population analysis of Arabidopsis pericycle in the *anac032* mutant.

**Supplementary Figure S38**. Sub-population analysis of Arabidopsis procambium in the *anac032* mutant.

**Supplementary Figure S39**. Sub-population analysis of Arabidopsis phloem in the *anac032* mutant.

**Supplementary Figure S40**. Common DEGs were found in the cortex, endodermis and pericycle of Arabidopsis wild type and mutants.

**Supplementary Figure S41**. GO terms enriched in the DEGs in the cortex, endodermis and pericycle of Arabidopsis wild type and mutants.

**Supplementary Figure S42**. Summary of DEGs in Arabidopsis cell types.

**Supplementary Figure S43**. Nitrate responses of AtNLP7 targets across Arabidopsis genotypes.

**Supplementary Figure S44**. Over-representation of cell type-enriched ribosome-associated transcripts (Kajala et al., 2021) within N concentration-dependent differentially expressed genes in the tomato whole root (Bian et al., 2025).

**Supplementary Figure S45**. Cell and read distributions of tomato scRNAseq samples from different KNO_3_ conditions and filtering cutoffs.

**Supplementary Figure S46**. Cell type prediction of tomato scRNAseq dataset from different KNO_3_ conditions.

**Supplementary Figure S47**. Refinement of annotation of tomato scRNAseq clusters from different KNO_3_ conditions.

**Supplementary Figure S48**. Validation of cell type annotation of tomato scRNAseq from different KNO_3_ conditions using marker genes with known expression patterns.

**Supplementary Figure S49**. Overlaps of DEGs between different KNO_3_ conditions in tomato root cell types.

**Supplementary Figure S50**. Expression of N-responsive genes in tomato root cell types.

**Supplementary Figure S51**. Sub-population analysis of tomato endodermis.

**Supplementary Figure S52**. Sub-population analysis of tomato atrichoblast.

**Supplementary Figure S53**. Integration of Arabidopsis (wild-type) and tomato scRNAseq datasets by combining transcripts of all genes in a homology group based on Kajala et al. (2021) expressologs.

**Supplementary Figure S54**. Integration of Arabidopsis (wild-type) and tomato scRNAseq datasets by combining transcripts of all genes in a homology group based on EnsemblPlant homology database.

**Supplementary Figure S55**. Integration of Arabidopsis (wild-type) and tomato scRNAseq datasets by taking averages of all genes in a homology group based on Kajala et al. (2021) expressologs.

**Supplementary Figure S56**. Integration of Arabidopsis (wild-type) and tomato scRNAseq datasets by taking averages of all genes in a homology group based on EnsemblPlant homology database

**Supplementary Figure S57**. Integration of Arabidopsis (wild-type) and tomato scRNAseq datasets by using the most abundant gene in a homology group based on Kajala et al. (2021) expressologs.

**Supplementary Figure S58**. Integration of Arabidopsis (wild-type) and tomato scRNAseq datasets by using the most abundant gene in a homology group based on EnsemblPlant homology database.

**Supplementary Figure S59**. Integration of Arabidopsis (wild-type) and tomato scRNAseq datasets by one-to-one orthologs based on Kajala et al. (2021) expressologs.

**Supplementary Figure S60**. Integration of Arabidopsis (wild-type) and tomato scRNAseq datasets by one-to-one orthologs based on EnsemblPlant homology database.

**Supplementary Figure S61**. MapMan broad category enrichment of nitrogen-responsive genes across tomato root cell types.

**Supplementary Figure S62**. Cell type expression profiles of AtNLP6 and AtNLP7 in low (1 mM) vs high (10 mM) KNO_3_.

**Supplementary Figure S63**. Experimental workflow for scRNA-seq of the modified TARGET assay.

**Supplementary Figure S64**. Cell type prediction of tomato root scRNA-seq from the modified TARGET assay.

**Supplementary Figure S65**. Validation of cell type annotation of tomato scRNAseq dataset from the modified TARGET assay.

**Supplementary Figure S66**. UMAP visualisation of cell type annotations and Seurat clusters of tomato scRNAseq samples from the modified TARGET assay.

**Supplementary Figure S67**. *SlNIR1* and *SlNIR2* were up-regulated in response to CD compared to CHX treatment.

**Supplementary Figure S68**. Expression of putative tomato orthologs of *AtABFs* in tomato root cell types at different KNO_3_ conditions.

## Notes

### Competing Interest Statement

The authors have declared no competing interest.

## References

1. Gutiérrez, R.A., Lejay, L.V., Dean, A., Chiaromonte, F., Shasha, D.E., and Coruzzi, G.M. (2007). Qualitative network models and genome-wide expression data define carbon/nitrogen-responsive molecular machines in Arabidopsis. Genome Biology 8, R7. 10.1186/gb-2007-8-1-r7.

2. Varala, K., Marshall-Colón, A., Cirrone, J., Brooks, M.D., Pasquino, A.V., Léran, S., Mittal, S., Rock, T.M., Edwards, M.B., Kim, G.J., et al. (2018). Temporal transcriptional logic of dynamic regulatory networks underlying nitrogen signaling and use in plants. Proceedings of the National Academy of Sciences 115, 6494–6499. 10.1073/pnas.1721487115.

3. Gaudinier, A., Rodriguez-Medina, J., Zhang, L., Olson, A., Liseron-Monfils, C., Bågman, A.M., Foret, J., Abbitt, S., Tang, M., Li, B., et al. (2018). Transcriptional regulation of nitrogen-associated metabolism and growth. Nature 563, 259–264. 10.1038/s41586-018-0656-3.

4. Brooks, M.D., Cirrone, J., Pasquino, A.V., Alvarez, J.M., Swift, J., Mittal, S., Juang, C.-L., Varala, K., Gutiérrez, R.A., Krouk, G., et al. (2019). Network Walking charts transcriptional dynamics of nitrogen signaling by integrating validated and predicted genome-wide interactions. Nature Communications 10, 1569. 10.1038/s41467-019-09522-1.

5. Alvarez, J.M., Schinke, A.-L., Brooks, M.D., Pasquino, A., Leonelli, L., Varala, K., Safi, A., Krouk, G., Krapp, A., and Coruzzi, G.M. (2020). Transient genome-wide interactions of the master transcription factor NLP7 initiate a rapid nitrogen-response cascade. Nature Communications 11, 1157. 10.1038/s41467-020-14979-6.

6. Alvarez, J.M., Riveras, E., Vidal, E.A., Gras, D.E., Contreras-López, O., Tamayo, K.P., Aceituno, F., Gómez, I., Ruffel, S., Lejay, L., et al. (2014). Systems approach identifies TGA1 and TGA4 transcription factors as important regulatory components of the nitrate response of Arabidopsis thaliana roots. Plant J. 80, 1–13. 10.1111/tpj.12618.

7. Castaings, L., Camargo, A., Pocholle, D., Gaudon, V., Texier, Y., Boutet-Mercey, S., Taconnat, L., Renou, J.-P., Daniel-Vedele, F., Fernandez, E., et al. (2009). The nodule inception-like protein 7 modulates nitrate sensing and metabolism in Arabidopsis. Plant J 57, 426–435. 10.1111/j.1365-313X.2008.03695.x.

8. Marchive, C., Roudier, F., Castaings, L., Bréhaut, V., Blondet, E., Colot, V., Meyer, C., and Krapp, A. (2013). Nuclear retention of the transcription factor NLP7 orchestrates the early response to nitrate in plants. Nat Commun 4, 1713. 10.1038/ncomms2650.

9. Liu, K.-H., Liu, M., Lin, Z., Wang, Z.-F., Chen, B., Liu, C., Guo, A., Konishi, M., Yanagisawa, S., Wagner, G., et al. (2022). NIN-like protein 7 transcription factor is a plant nitrate sensor. Science. 10.1126/science.add1104.

10. Cheng, Y.-H., Durand, M., Brehaut, V., Hsu, F.-C., Kelemen, Z., Texier, Y., Krapp, A., and Tsay, Y.-F. (2023). Interplay between NIN-LIKE PROTEINs 6 and 7 in nitrate signaling. Plant Physiol 192, 3049–3068. 10.1093/plphys/kiad242.

11. Bian, C., Demirer, G.S., Oz, M.T., Cai, Y.-M., Witham, S., Mason, G.A., Di, Z., Deligne, F., Zhang, P., Shen, R., et al. (2025). Conservation and divergence of regulatory architecture in nitrate-responsive plant gene circuits. Plant Cell 37. 10.1093/plcell/koaf124.

12. Liu, K.H., Niu, Y., Konishi, M., Wu, Y., Du, H., Sun Chung, H., Li, L., Boudsocq, M., McCormack, M., Maekawa, S., et al. (2017). Discovery of nitrate-CPK-NLP signalling in central nutrient-growth networks. Nature 545, 311–316. 10.1038/nature22077.

13. Guan, P., Ripoll, J.-J., Wang, R., Vuong, L., Bailey-Steinitz, L.J., Ye, D., and Crawford, N.M. (2017). Interacting TCP and NLP transcription factors control plant responses to nitrate availability. Proc Natl Acad Sci U S A 114, 2419–2424. 10.1073/pnas.1615676114.

14. Konishi, M., and Yanagisawa, S. (2019). The role of protein-protein interactions mediated by the PB1 domain of NLP transcription factors in nitrate-inducible gene expression. BMC Plant Biol 19, 90. 10.1186/s12870-019-1692-3.

15. Vidal, E.A., Álvarez, J.M., Moyano, T.C., and Gutiérrez, R.A. (2015). Transcriptional networks in the nitrate response of Arabidopsis thaliana. Current Opinion in Plant Biology 27, 125–132. 10.1016/j.pbi.2015.06.010.

16. Lamig, L., Moreno, S., Álvarez, J.M., and Gutiérrez, R.A. (2022). Molecular mechanisms underlying nitrate responses in plants. Curr. Biol. 32, R433–R439. 10.1016/j.cub.2022.03.022.

17. Undurraga, S.F., Ibarra-Henríquez, C., Fredes, I., Álvarez, J.M., and Gutiérrez, R.A. (2017). Nitrate signaling and early responses in Arabidopsis roots. J Exp Bot 68, 2541–2551. 10.1093/jxb/erx041.

18. Canales, J., Contreras-López, O., Álvarez, J.M., and Gutiérrez, R.A. (2017). Nitrate induction of root hair density is mediated by TGA1/TGA4 and CPC transcription factors in Arabidopsis thaliana. Plant J 92, 305–316. 10.1111/tpj.13656.

19. Vidal, E.A., Moyano, T.C., Canales, J., and Gutiérrez, R.A. (2014). Nitrogen control of developmental phase transitions in Arabidopsis thaliana. J Exp Bot 65, 5611–5618. 10.1093/jxb/eru326.

20. Vermeer, J.E.M., and Geldner, N. (2015). Lateral root initiation in Arabidopsis thaliana: a force awakens. F1000Prime Rep 7, 32. 10.12703/P7-32.

21. Masclaux-Daubresse, C., Clément, G., Anne, P., Routaboul, J.-M., Guiboileau, A., Soulay, F., Shirasu, K., and Yoshimoto, K. (2014). Stitching together the Multiple Dimensions of Autophagy Using Metabolomics and Transcriptomics Reveals Impacts on Metabolism, Development, and Plant Responses to the Environment in Arabidopsis. Plant Cell 26, 1857–1877. 10.1105/tpc.114.124677.

22. Gifford, M.L., Dean, A., Gutierrez, R.A., Coruzzi, G.M., and Birnbaum, K.D. (2008). Cell-specific nitrogen responses mediate developmental plasticity. Proc Natl Acad Sci U S A 105, 803–808. 10.1073/pnas.0709559105.

23. Walker, L., Boddington, C., Jenkins, D., Wang, Y., Grønlund, J.T., Hulsmans, J., Kumar, S., Patel, D., Moore, J.D., Carter, A., et al. (2017). Changes in Gene Expression in Space and Time Orchestrate Environmentally Mediated Shaping of Root Architecture. Plant Cell 29, 2393–2412. 10.1105/tpc.16.00961.

24. Contreras-López, O., Vidal, E.A., Riveras, E., Alvarez, J.M., Moyano, T.C., Sparks, E.E., Medina, J., Pasquino, A., Benfey, P.N., Coruzzi, G.M., et al. (2022). Spatiotemporal analysis identifies ABF2 and ABF3 as key hubs of endodermal response to nitrate. Proc Natl Acad Sci U S A 119. 10.1073/pnas.2107879119.

25. Kajala, K., Gouran, M., Shaar-Moshe, L., Mason, G.A., Rodriguez-Medina, J., Kawa, D., Pauluzzi, G., Reynoso, M., Canto-Pastor, A., Manzano, C., et al. (2021). Innovation, conservation, and repurposing of gene function in root cell type development. Cell 184, 3333–3348.e19. 10.1016/j.cell.2021.04.024.

26. Shulse, C.N., Cole, B.J., Ciobanu, D., Lin, J., Yoshinaga, Y., Gouran, M., Turco, G.M., Zhu, Y., O’Malley, R.C., Brady, S.M., et al. (2019). High-Throughput Single-Cell Transcriptome Profiling of Plant Cell Types. Cell Rep. 27, 2241–2247.e4. 10.1016/j.celrep.2019.04.054.

27. Ryu, K.H., Huang, L., Kang, H.M., and Schiefelbein, J. (2019). Single-cell RNA sequencing resolves molecular relationships among individual plant cells. Plant Physiol. 179, 1444–1456. 10.1104/pp.18.01482.

28. Zhang, T.-Q., Xu, Z.-G., Shang, G.-D., and Wang, J.-W. (2019). A Single-Cell RNA Sequencing Profiles the Developmental Landscape of Arabidopsis Root. Mol Plant 12, 648–660. 10.1016/j.molp.2019.04.004.

29. Shahan, R., Hsu, C.-W., Nolan, T.M., Cole, B.J., Taylor, I.W., Greenstreet, L., Zhang, S., Afanassiev, A., Vlot, A.H.C., Schiebinger, G., et al. (2022). A single-cell Arabidopsis root atlas reveals developmental trajectories in wild-type and cell identity mutants. Dev Cell 57, 543–560.e9. 10.1016/j.devcel.2022.01.008.

30. Wendrich, J.R., Yang, B., Vandamme, N., Verstaen, K., Smet, W., Van de Velde, C., Minne, M., Wybouw, B., Mor, E., Arents, H.E., et al. (2020). Vascular transcription factors guide plant epidermal responses to limiting phosphate conditions. Science 370. 10.1126/science.aay4970.

31. Jean-Baptiste, K., McFaline-Figueroa, J.L., Alexandre, C.M., Dorrity, M.W., Saunders, L., Bubb, K.L., Trapnell, C., Fields, S., Queitsch, C., and Cuperus, J.T. (2019). Dynamics of Gene Expression in Single Root Cells of. Plant Cell 31, 993–1011. 10.1105/tpc.18.00785.

32. Zhu, M., Hsu, C.-W., Peralta Ogorek, L.L., Taylor, I.W., La Cavera, S., Oliveira, D.M., Verma, L., Mehra, P., Mijar, M., Sadanandom, A., et al. (2025). Single-cell transcriptomics reveal how root tissues adapt to soil stress. Nature 642, 721–729. 10.1038/s41586-025-08941-z.

33. Nolan, T.M., Vukašinović, N., Hsu, C.-W., Zhang, J., Vanhoutte, I., Shahan, R., Taylor, I.W., Greenstreet, L., Heitz, M., Afanassiev, A., et al. (2023). Brassinosteroid gene regulatory networks at cellular resolution in the root. Science 379, eadf4721. 10.1126/science.adf4721.

34. Hai, C., Li, Y., Peng, C., Hu, L., and Xu, W. (2025). Single-cell nuclear transcriptomics reveal root tip adaptations to nitrogen scarcity in wheat. The Crop Journal 13, 1156–1167. 10.1016/j.cj.2025.05.010.

35. Patrick, R.M., Ranjan, R., Sumanasinghe, S.R., SanMiguel, P.J., Varala, K., and Li, Y. (2026). Every Cell Counts: Tomato Root Responses to Nitrogen at Single-Cell Resolution. bioRxiv, 2026.02.06.704465. 10.64898/2026.02.06.704465.

36. Efroni, I., Ip, P.-L., Nawy, T., Mello, A., and Birnbaum, K.D. (2015). Quantification of cell identity from single-cell gene expression profiles. Genome Biol 16, 9. 10.1186/s13059-015-0580-x.

37. Denyer, T., Ma, X., Klesen, S., Scacchi, E., Nieselt, K., and Timmermans, M.C.P. (2019). Spatiotemporal Developmental Trajectories in the Arabidopsis Root Revealed Using High-Throughput Single-Cell RNA Sequencing. Dev Cell 48, 840–852.e5. 10.1016/j.devcel.2019.02.022.

38. Gulati, G.S., Sikandar, S.S., Wesche, D.J., Manjunath, A., Bharadwaj, A., Berger, M.J., Ilagan, F., Kuo, A.H., Hsieh, R.W., Cai, S., et al. (2020). Single-cell transcriptional diversity is a hallmark of developmental potential. Science 367, 405–411. 10.1126/science.aax0249.

39. Cantó-Pastor, A., Kajala, K., Shaar-Moshe, L., Manzano, C., Timilsena, P., De Bellis, D., Gray, S., Holbein, J., Yang, H., Mohammad, S., et al. (2024). A suberized exodermis is required for tomato drought tolerance. Nat Plants 10, 118–130. 10.1038/s41477-023-01567-x.

40. Thimm, O., Bläsing, O., Gibon, Y., Nagel, A., Meyer, S., Krüger, P., Selbig, J., Müller, L.A., Rhee, S.Y., and Stitt, M. (2004). MAPMAN: a user-driven tool to display genomics data sets onto diagrams of metabolic pathways and other biological processes. Plant J 37, 914–939. 10.1111/j.1365-313x.2004.02016.x.

41. Hsu, C.-W., Shahan, R., Nolan, T.M., Benfey, P.N., and Ohler, U. (2022). Protocol for fast scRNA-seq raw data processing using scKB and non-arbitrary quality control with COPILOT. STAR Protoc 3, 101729. 10.1016/j.xpro.2022.101729.

42. Pan, X., Wang, C., Liu, Z., Gao, R., Feng, L., Li, A., Yao, K., and Liao, W. (2023). Identification of ABF/AREB gene family in tomato (L.) and functional analysis of in response to ABA and abiotic stresses. PeerJ 11, e15310. 10.7717/peerj.15310.

43. Orellana, S., Yañez, M., Espinoza, A., Verdugo, I., González, E., Ruiz-Lara, S., and Casaretto, J.A. (2010). The transcription factor SlAREB1 confers drought, salt stress tolerance and regulates biotic and abiotic stress-related genes in tomato. Plant Cell Environ 33, 2191–2208. 10.1111/j.1365-3040.2010.02220.x.

44. Kurihara, D., Mizuta, Y., Sato, Y., and Higashiyama, T. (2015). ClearSee: a rapid optical clearing reagent for whole-plant fluorescence imaging. Development 142, 4168–4179. 10.1242/dev.127613.

45. Menz, J., Li, Z., Schulze, W.X., and Ludewig, U. (2016). Early nitrogen-deprivation responses in Arabidopsis roots reveal distinct differences on transcriptome and (phospho-) proteome levels between nitrate and ammonium nutrition. Plant J 88, 717–734. 10.1111/tpj.13272.

46. Wang, G., Ryu, K.H., Dinneny, A., Lee, J., Oh, D.-H., Ramachandran, P., Oliva, M., Lister, R., Dinneny, J.R., Schiefelbein, J., et al. (2026). Evolutionary diversity of cell-type-specific expression and stress response in Brassicaceae roots. Nature Communications, 1–21. 10.1038/s41467-026-73270-2.

47. Chen, Y.-N., Cartwright, H.N., and Ho, C.-H. (2022). In vivo visualization of nitrate dynamics using a genetically encoded fluorescent biosensor. Sci Adv 8, eabq4915. 10.1126/sciadv.abq4915.

48. Liu, B., Wu, J., Yang, S., Schiefelbein, J., and Gan, Y. (2020). Nitrate regulation of lateral root and root hair development in plants. J Exp Bot 71, 4405–4414. 10.1093/jxb/erz536.

49. Huang, N.C., Chiang, C.S., Crawford, N.M., and Tsay, Y.F. (1996). CHL1 encodes a component of the low-affinity nitrate uptake system in Arabidopsis and shows cell type-specific expression in roots. Plant Cell 8, 2183–2191. 10.1105/tpc.8.12.2183.

50. Remans, T., Nacry, P., Pervent, M., Filleur, S., Diatloff, E., Mounier, E., Tillard, P., Forde, B.G., and Gojon, A. (2006). The Arabidopsis NRT1.1 transporter participates in the signaling pathway triggering root colonization of nitrate-rich patches. Proc Natl Acad Sci U S A 103, 19206–19211. 10.1073/pnas.0605275103.

51. Nazoa, P., Vidmar, J.J., Tranbarger, T.J., Mouline, K., Damiani, I., Tillard, P., Zhuo, D., Glass, A.D.M., and Touraine, B. (2003). Regulation of the nitrate transporter gene AtNRT2.1 in Arabidopsis thaliana: responses to nitrate, amino acids and developmental stage. Plant Mol Biol 52, 689–703. 10.1023/a:1024899808018.

52. Wirth, J., Chopin, F., Santoni, V., Viennois, G., Tillard, P., Krapp, A., Lejay, L., Daniel-Vedele, F., and Gojon, A. (2007). Regulation of root nitrate uptake at the NRT2.1 protein level in Arabidopsis thaliana. J Biol Chem 282, 23541–23552. 10.1074/jbc.M700901200.

53. Bargmann, B.O.R., Marshall-Colon, A., Efroni, I., Ruffel, S., Birnbaum, K.D., Coruzzi, G.M., and Krouk, G. (2013). TARGET: a transient transformation system for genome-wide transcription factor target discovery. Mol. Plant 6, 978–980. 10.1093/mp/sst010.

54. Barberon, M., Vermeer, J.E.M., De Bellis, D., Wang, P., Naseer, S., Andersen, T.G., Humbel, B.M., Nawrath, C., Takano, J., Salt, D.E., et al. (2016). Adaptation of Root Function by Nutrient-Induced Plasticity of Endodermal Differentiation. Cell 164, 447–459. 10.1016/j.cell.2015.12.021.

55. Kreszies, T., Shellakkutti, N., Osthoff, A., Yu, P., Baldauf, J.A., Zeisler-Diehl, V.V., Ranathunge, K., Hochholdinger, F., and Schreiber, L. (2019). Osmotic stress enhances suberization of apoplastic barriers in barley seminal roots: analysis of chemical, transcriptomic and physiological responses. New Phytol 221, 180–194. 10.1111/nph.15351.

56. Patro, R., Duggal, G., Love, M.I., Irizarry, R.A., and Kingsford, C. (2017). Salmon provides fast and bias-aware quantification of transcript expression. Nat Methods 14, 417–419. 10.1038/nmeth.4197.

57. Soneson, C., Love, M.I., and Robinson, M.D. (2015). Differential analyses for RNA-seq: transcript-level estimates improve gene-level inferences. F1000Res 4, 1521. 10.12688/f1000research.7563.2.

58. Love, M.I., Huber, W., and Anders, S. (2014). Moderated estimation of fold change and dispersion for RNA-seq data with DESeq2. Genome Biol 15, 550. 10.1186/s13059-014-0550-8.

59. Rodriguez Gallo, M.C., Li, Q., Talasila, M., and Uhrig, R.G. (2023). Quantitative Time-Course Analysis of Osmotic and Salt Stress in Arabidopsis thaliana Using Short Gradient Multi-CV FAIMSpro BoxCar DIA. Mol Cell Proteomics 22, 100638. 10.1016/j.mcpro.2023.100638.

60. Mehta, D., Scandola, S., and Uhrig, R.G. (2022). BoxCar and Library-Free Data-Independent Acquisition Substantially Improve the Depth, Range, and Completeness of Label-Free Quantitative Proteomics. Anal Chem 94, 793–802. 10.1021/acs.analchem.1c03338.

61. Ron, M., Kajala, K., Pauluzzi, G., Wang, D., Reynoso, M. a., Zumstein, K., Garcha, J., Winte Masson, H., Inagaki, S., et al. (2014). Hairy root transformation using Agrobacterium rhizogenes as a tool for exploring cell type-specific gene expression and function using tomato as a model. Plant Physiol. 166, 455–469. 10.1104/pp.114.239392.

62. Yoo, S.D., Cho, Y.H., and Sheen, J. (2007). Arabidopsis mesophyll protoplasts: A versatile cell system for transient gene expression analysis. Nat. Protoc. 2, 1565–1572. 10.1038/nprot.2007.199.

63. Haese-Hill, W., Crouch, K., and Otto, T.D. (2023). peaks2utr: a robust Python tool for the annotation of 3’ UTRs. Bioinformatics 39. 10.1093/bioinformatics/btad112.

64. Hao, Y., Stuart, T., Kowalski, M.H., Choudhary, S., Hoffman, P., Hartman, A., Srivastava, A., Molla, G., Madad, S., Fernandez-Granda, C., et al. (2024). Dictionary learning for integrative, multimodal and scalable single-cell analysis. Nat Biotechnol 42, 293–304. 10.1038/s41587-023-01767-y.

65. McGinnis, C.S., Murrow, L.M., and Gartner, Z.J. (2019). DoubletFinder: Doublet Detection in Single-Cell RNA Sequencing Data Using Artificial Nearest Neighbors. Cell Syst 8, 329–337.e4. 10.1016/j.cels.2019.03.003.

66. Germain, P.-L., Lun, A., Garcia Meixide, C., Macnair, W., and Robinson, M.D. (2021). Doublet identification in single-cell sequencing data using. F1000Res 10, 979. 10.12688/f1000research.73600.2.

67. Korsunsky, I., Millard, N., Fan, J., Slowikowski, K., Zhang, F., Wei, K., Baglaenko, Y., Brenner, M., Loh, P.-R., and Raychaudhuri, S. (2019). Fast, sensitive and accurate integration of single-cell data with Harmony. Nat Methods 16, 1289–1296. 10.1038/s41592-019-0619-0.

68. Birnbaum, K.D., and Kussell, E. (2011). Measuring cell identity in noisy biological systems. Nucleic Acids Res 39, 9093–9107. 10.1093/nar/gkr591.

69. Watanabe, S., Takahashi, N., Kanno, Y., Suzuki, H., Aoi, Y., Takeda-Kamiya, N., Toyooka, K., Kasahara, H., Hayashi, K.-I., Umeda, M., et al. (2020). The NRT1/PTR FAMILY protein NPF7.3/NRT1.5 is an indole-3-butyric acid transporter involved in root gravitropism. Proc Natl Acad Sci U S A 117, 31500–31509. 10.1073/pnas.2013305117.

70. Lin, S.-H., Kuo, H.-F., Canivenc, G., Lin, C.-S., Lepetit, M., Hsu, P.-K., Tillard, P., Lin, H.-L., Wang, Y.-Y., Tsai, C.-B., et al. (2008). Mutation of the Arabidopsis NRT1.5 nitrate transporter causes defective root-to-shoot nitrate transport. Plant Cell 20, 2514–2528. 10.1105/tpc.108.060244.

71. Manzano, C., Morimoto, K.W., Shaar-Moshe, L., Mason, G.A., Cantó-Pastor, A., Gouran, M., De Bellis, D., Ursache, R., Kajala, K., Sinha, N., et al. (2025). Regulation and function of a polarly localized lignin barrier in the exodermis. Nat Plants 11, 118–130. 10.1038/s41477-024-01864-z.

72. Ursache, R., Andersen, T.G., Marhavý, P., and Geldner, N. (2018). A protocol for combining fluorescent proteins with histological stains for diverse cell wall components. Plant J 93, 399–412. 10.1111/tpj.13784.

