## Supplementary Figure 1-68 for "Evolutionary Rewiring of Root Cell-Type Nitrogen Responses"

(a)

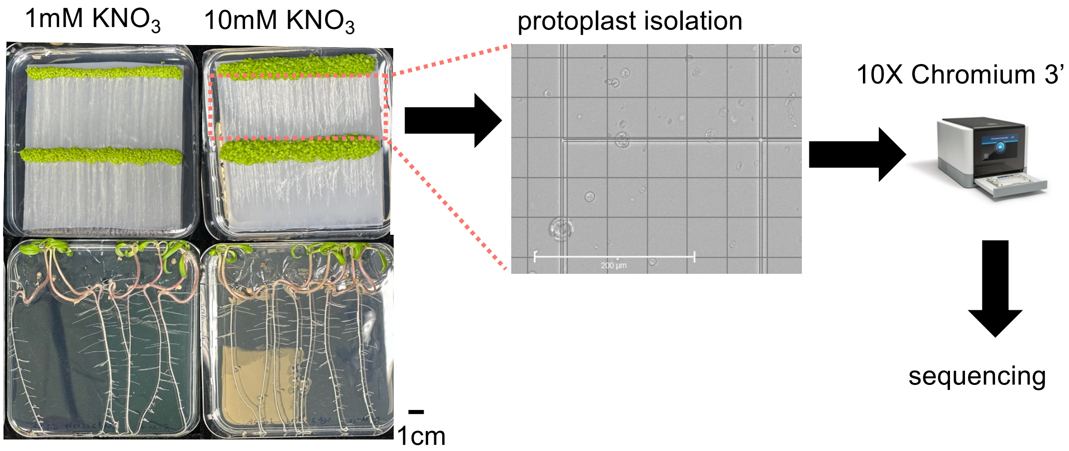


(b)

| Species | Genotype | 1 mM $KNO_3$ | 10 mM $KNO_3$ |
|---|---|---|---|
| *Arabidopsis thaliana* | Col-0 (WT) | 2 | 2 |
| *Arabidopsis thaliana* | *anac032* | 2 | 2 |
| *Arabidopsis thaliana* | *nlp7* | 1 | 2 |
| *Solanum lycopersicum* | M82 (WT) | 2 | 2 |

**Supplementary Figure S1. Experimental workflow for single-cell transcriptomics of Arabidopsis and tomato protoplasts. (a)** Arabidopsis and tomato seedlings were grown on MS media containing low (1 mM) or replete (10 mM) $KNO_3$ for ~10 days after germination. Protoplasts were harvested from whole roots using enzymatic cell wall digestion and encapsulated into Gel Beads-in-Emulsion (GEMs) using the 10X Chromium Next GEM Single Cell 3' system for single-cell library preparation. Libraries were sequenced on NovaSeq X Plus 10B flow cells. **(b)** Sample number for each plant genotype and $KNO_3$ treatment.

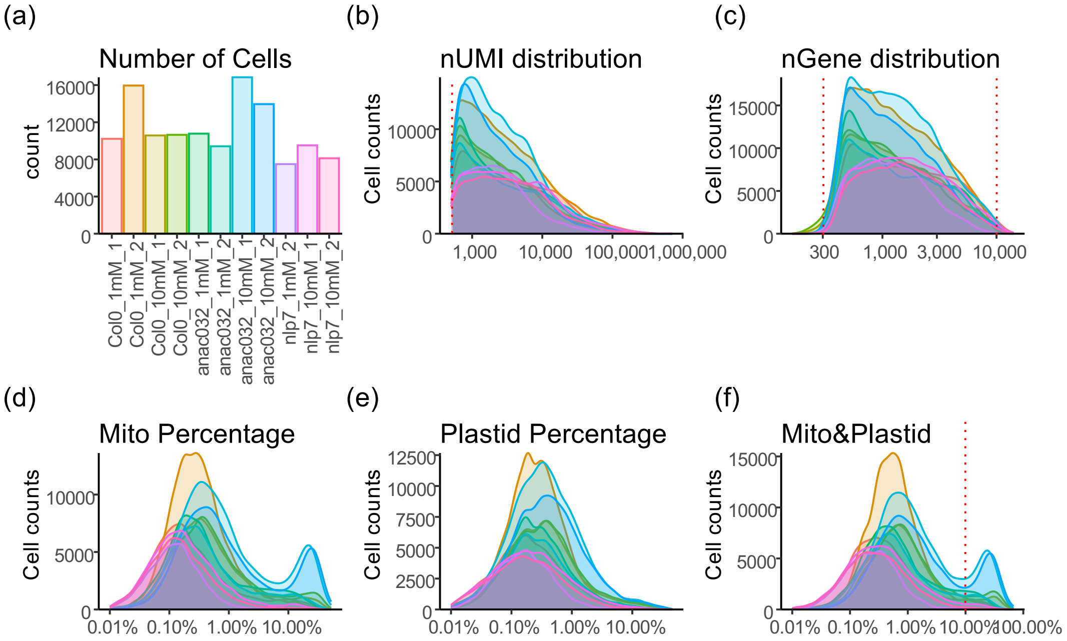


**Supplementary Figure S2. Cell and read distributions of Arabidopsis scRNAseq samples and filtering cutoffs. (a)** Number of cells in each sample before filtering. **(b-f)** Distribution of the number of nUMI (unique molecular identifier) (**b**), number of genes (**c**), percentage of mitochondrial transcripts (**d**), percentage of chloroplast transcripts (**e**), and percentage of the sum of mitochondrial and chloroplast transcripts (**f**) in each sample. Filtering thresholds are indicated by red dash lines (UMIs > 300, detected genes between 300 to 10000, and organellar (mitochondrial and chloroplast) transcripts <10%).

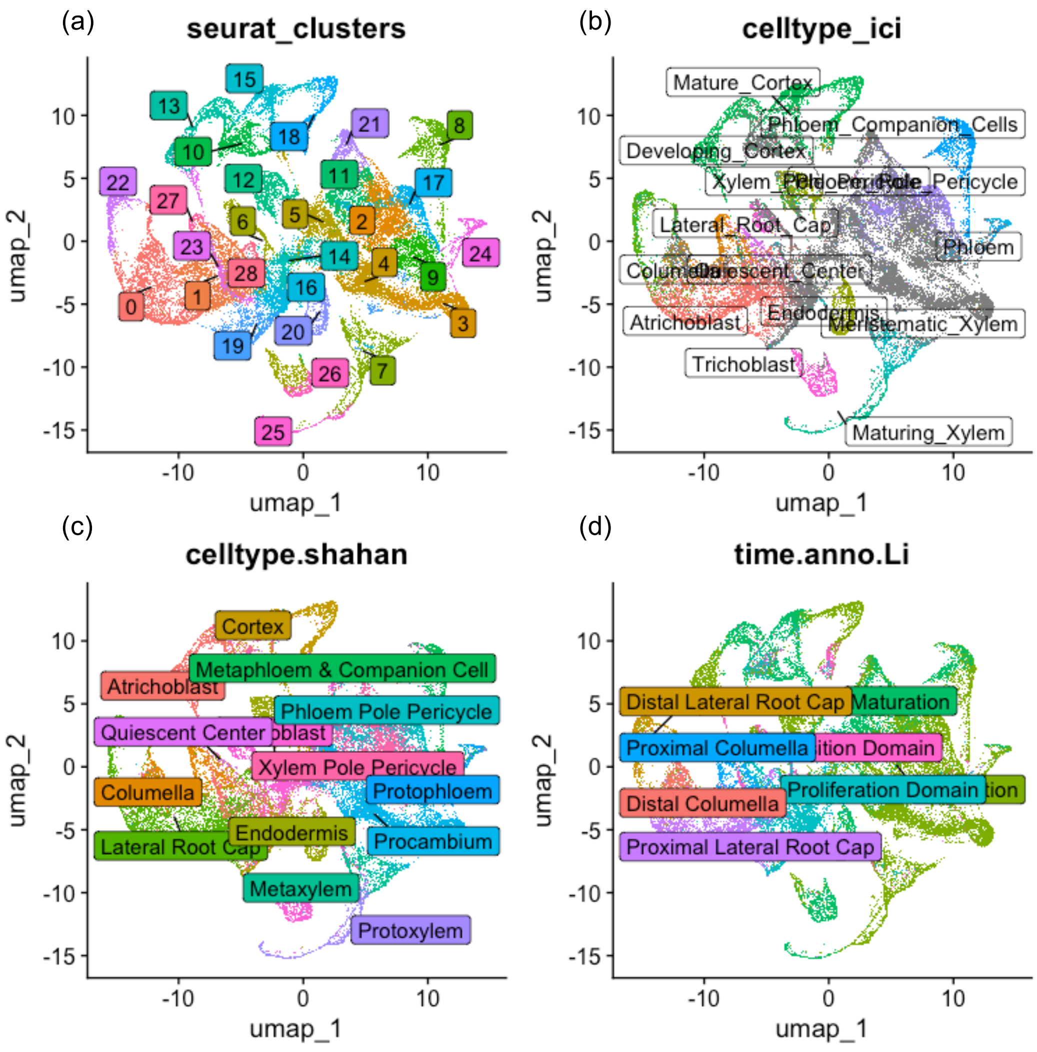


**Supplementary Figure S3. UMAP visualisation of Arabidopsis scRNAseq from different annotation methods. (a)** UMAP of Seurat clusters. **(b)** Cell type predictions by the Index of Cell Identity (ICI) method. **(c)** Cell type predictions by the label transfer method using the Arabidopsis root atlas (Shahan et al. 2022) as the reference. **(d)** Developmental stage predictions by label transfer using Shahan et al. (2022) Arabidopsis root atlas as the reference.

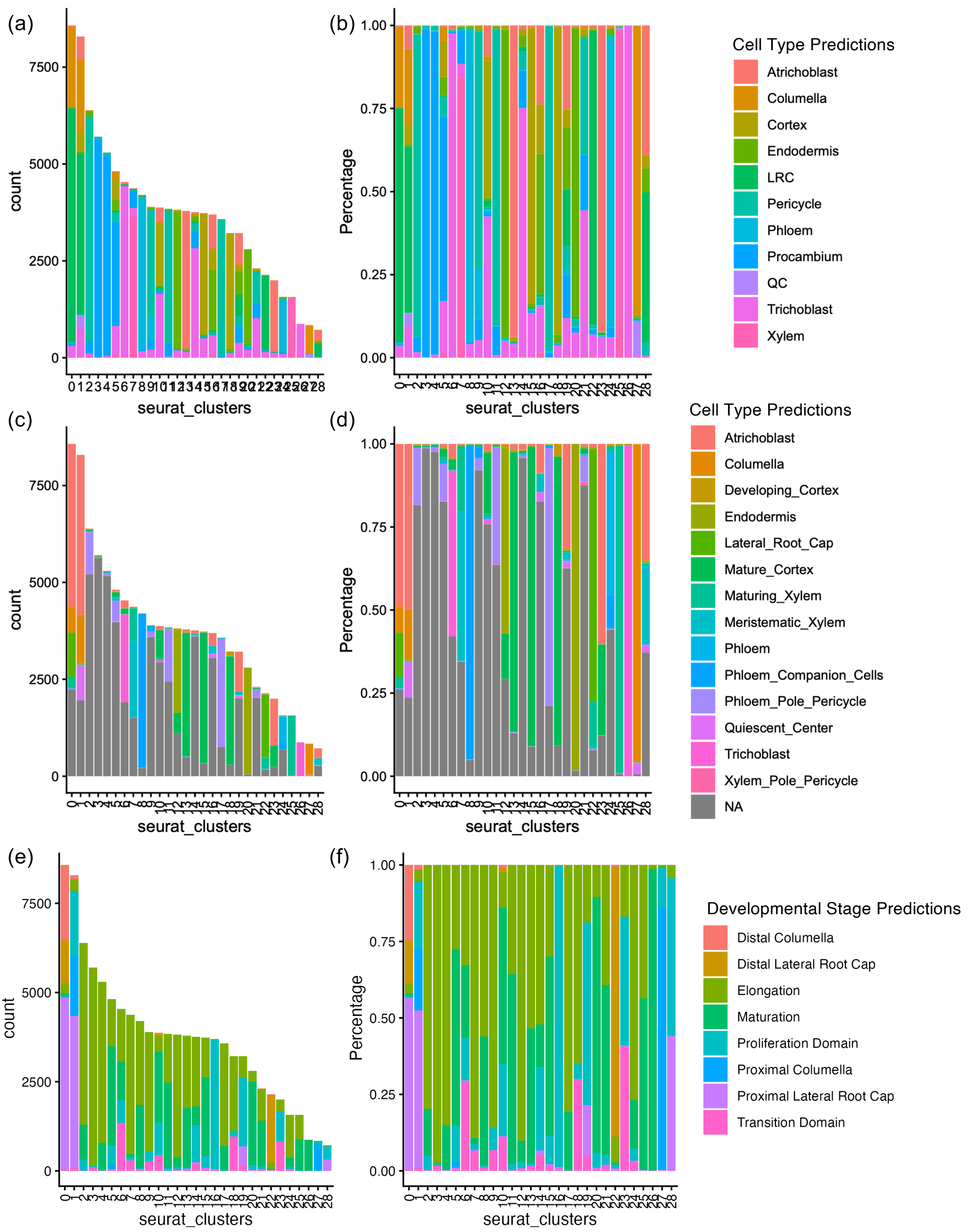


**Supplementary Figure S4. Cell type and cell stage predictions of Arabidopsis scRNAseq dataset from different annotation methods. (a)** The number and **(b)** percentage of predicted cell type labels in each cluster using the label transfer method. **(c)** The number and **(d)** percentage of predicted predicted cell type labels in each cluster using the ICI method. **(e)** The number and **(f)** percentage of predicted developmental stage labels in each cluster using the label transfer method.

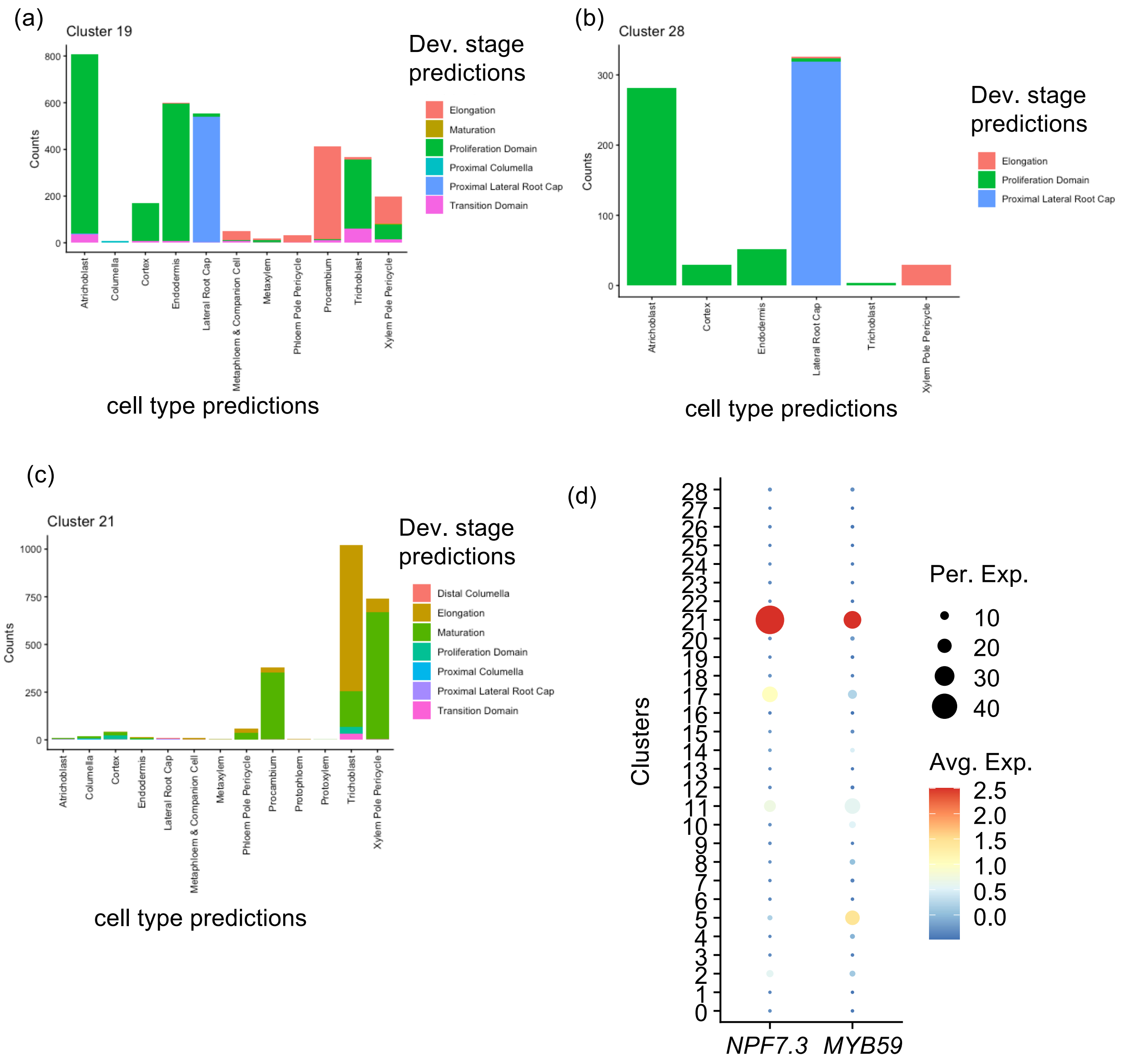


**Supplementary Figure S5. Refinement of annotation of Arabidopsis clusters 19, 21 and 28.** **(a)** Correlation of cell type predictions (via label transfer) and developmental (Dev.) stage predictions for cluster 19. **(b)** Correlation of cell type predictions (via label transfer) and developmental stage predictions for cluster 28. **(c)** Correlation of cell type predictions (via label transfer) and developmental stage predictions for cluster 21. **(d)** Expression of two cluster 21 markers which were known to be expressed in pericycle. Per. Exp., percentage of cells expressing the gene. Avg. Exp., scaled average expression level.

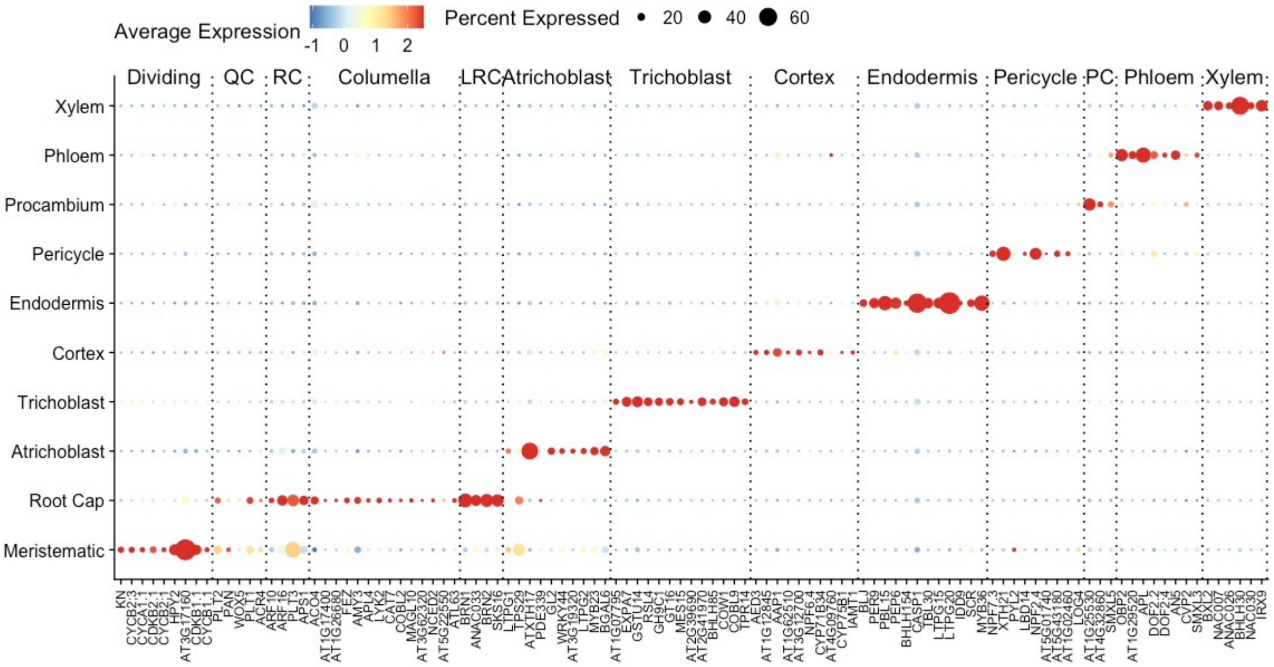


**Supplementary Figure S6. Validation of Arabidopsis cell type and stage annotation using marker genes with known expression patterns.**

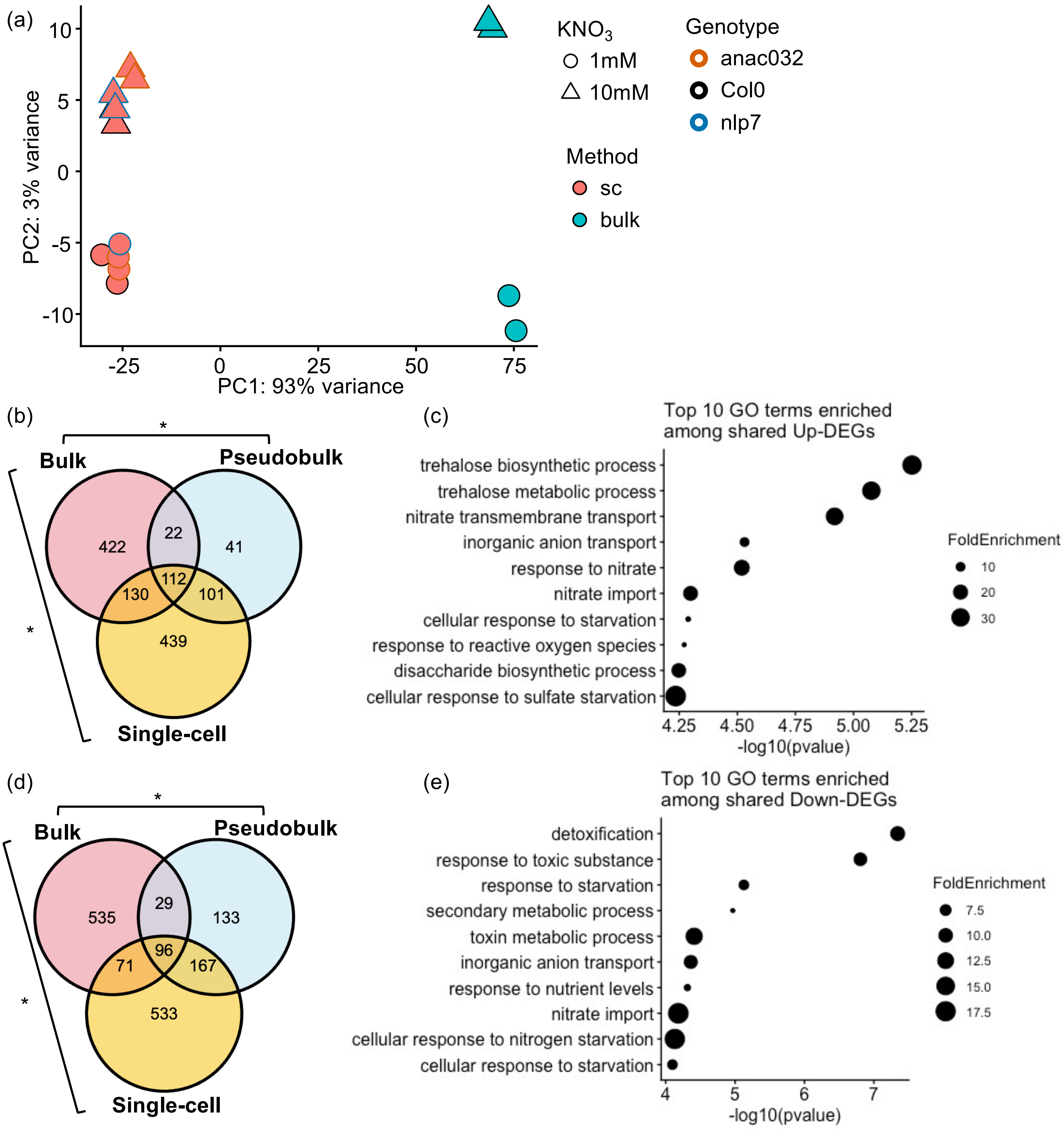


**Supplementary Figure S7. scRNAseq of Arabidopsis root protoplasts captures similar nitrate responses as RNAseq from RNA extracted from whole roots. (a)** Principal component analysis (PCA) of bulk transcriptomes of Arabidopsis roots without protoplasting and pseudobulked single-cell transcriptomes of Arabidopsis root protoplasts. **(b)** Venn plot showing the overlap of differentially expressed genes (DEGs) up-regulated at high $KNO_3$ in the whole-tissue transcriptome (bulk), pseudobulked single-cell transcriptome (Pseudobulk) and DEGs up-regulated in any cell types in the single-cell transcriptome (Single-cell) of wild-type Col-0 samples. **(c)** Top 10 GO terms enriched in the up-DEGs found in both bulk and pseudobulked samples. FoldEnrichment denotes the observed proportion divided by the background proportion for each GO term. (**d**) Venn plot **(e)** and GO enrichment for down-DEGs. Significance of overlaps between groups was evaluated using Fisher's Exact test, * indicates $p < 2.2*10^{-16}$.

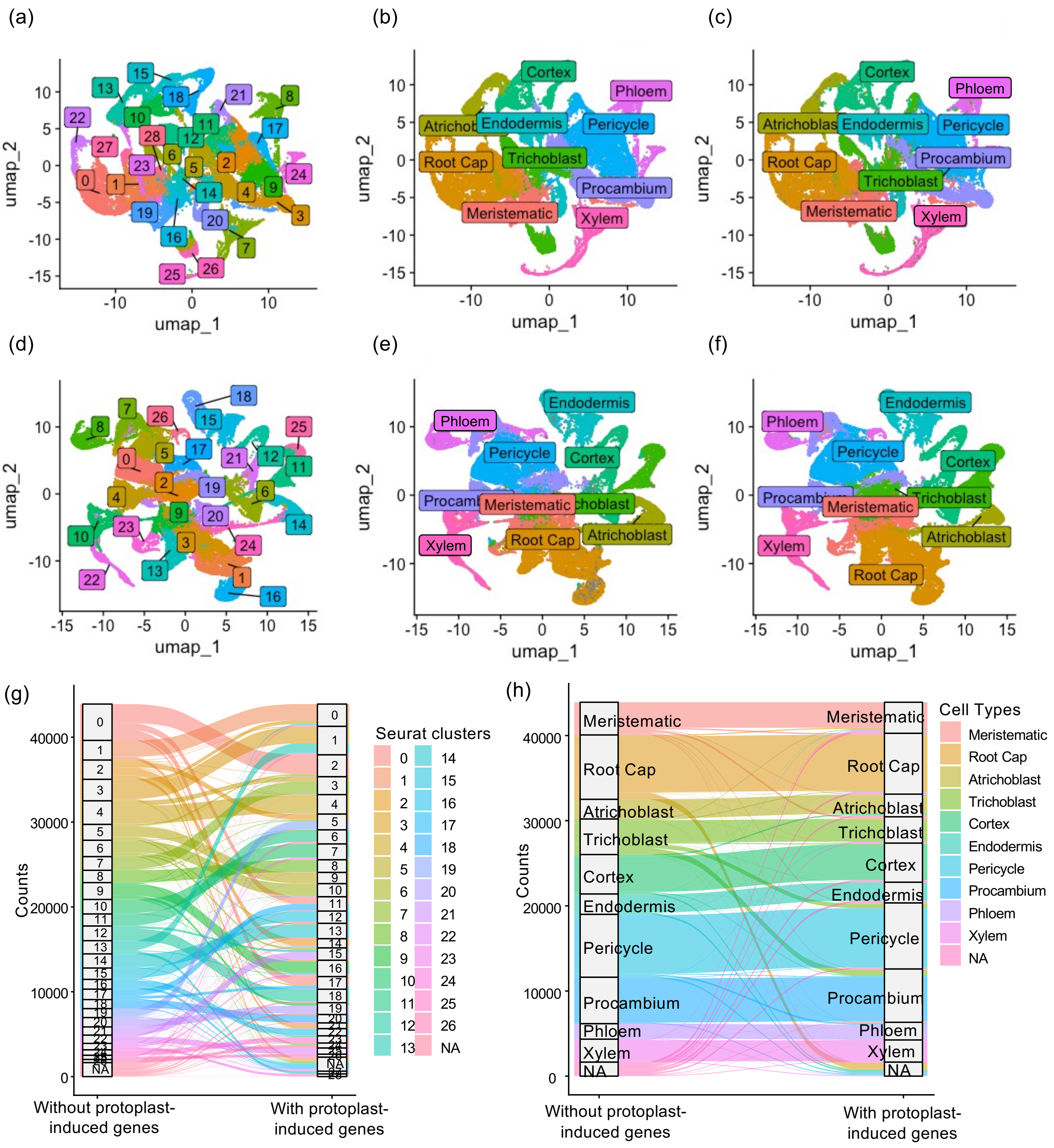


**Supplementary Figure S8. Clustering and annotation of Arabidopsis scRNAseq samples with and without protoplasting-induced genes.** (**a-c**) Col-0 samples processed and annotated with protoplasting-induced genes. (**a**) UMAP visualisation of Seurat clusters. (**b**) Cell type annotations. (**c**) Cell type annotation transferred from Col-0 samples processed without protoplasting-induced genes. (**d-f**) Col-0 samples processed and annotated without protoplasting-induced genes. (**d**) UMAP visualisation of Seurat clusters. (**e**) Cell type annotations. (**f**) Cell type annotation transferred from Col-0 samples processed with protoplasting-induced genes. (**g**) Correspondence of Seurat clusters between Col-0 samples processed with and without protoplasting-induced genes. (**h**) Correspondence of cell type annotations between Col-0 samples processed with and without protoplasting-induced genes.

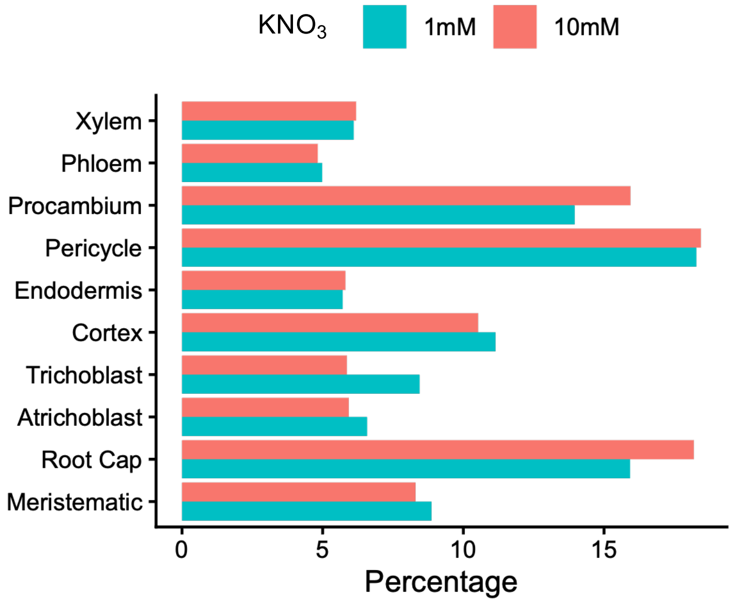


**Supplementary Figure S9. Cell type abundance in the Arabidopsis Col-0 scRNAseq samples.**

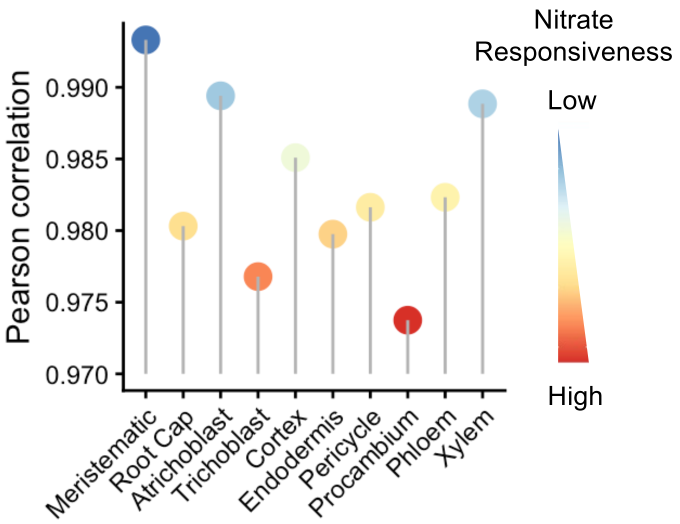


**Supplementary Figure S10. Pearson correlations between high and low nitrate pseudobulk transcriptomes across Arabidopsis root cell types.**

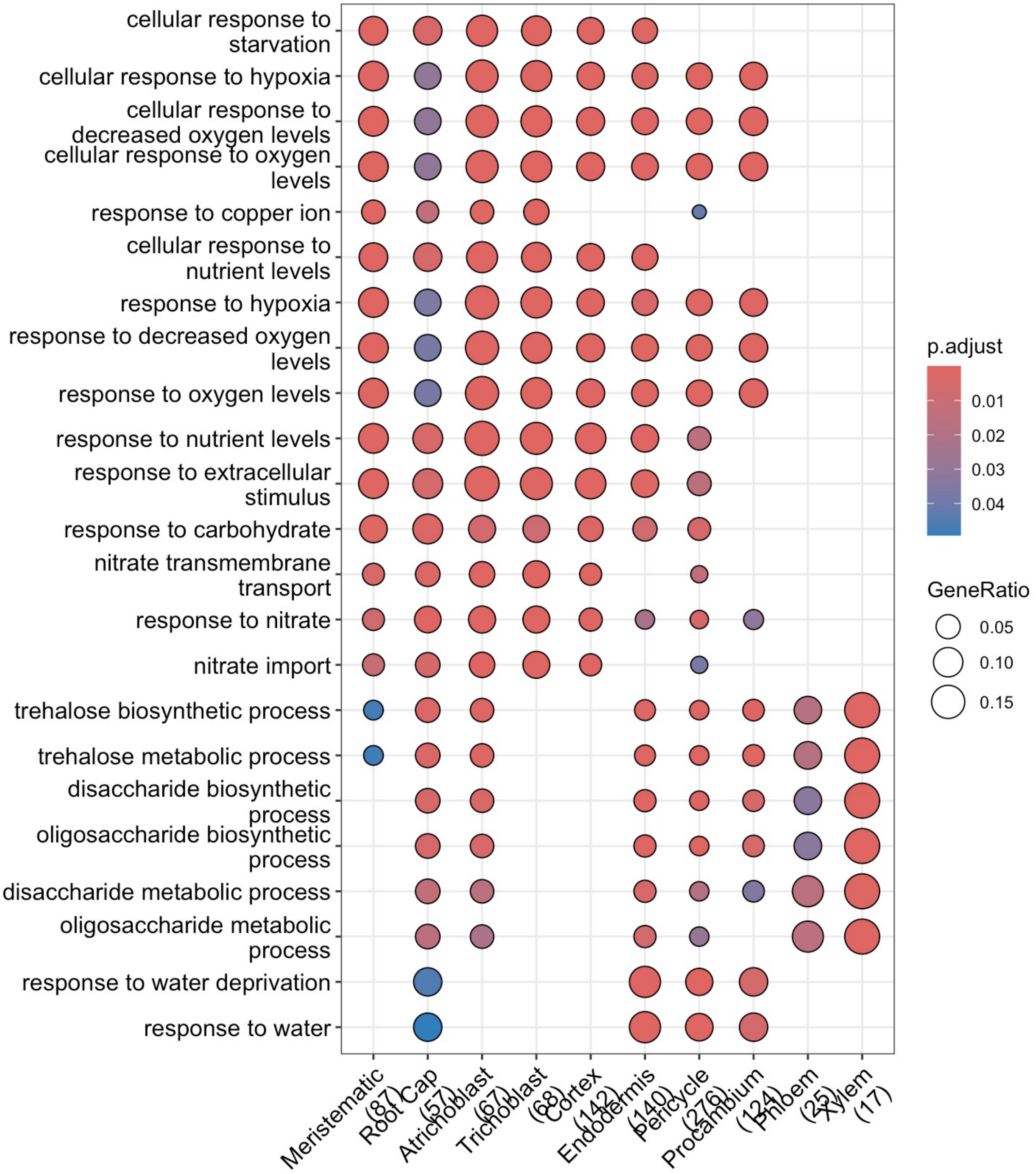


**Supplementary Figure S11. GO terms enriched in the DEGs up-regulated under high nitrate in Arabidopsis root cell types.** The top 5 GO terms for each cell type are shown.

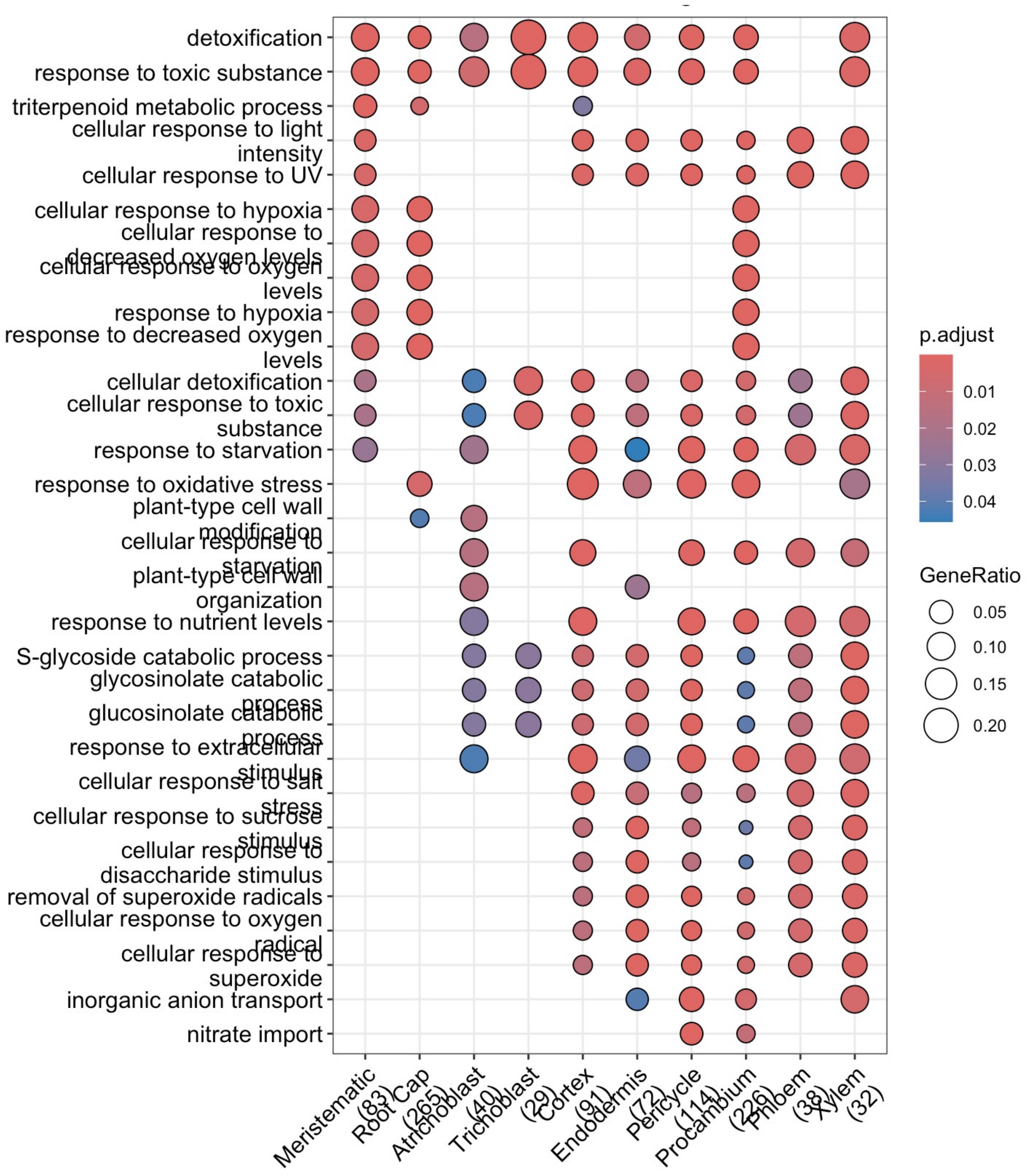


**Supplementary Figure S12. GO terms enriched in the DEGs down-regulated under high nitrate in Arabidopsis root cell types.** The top 5 GO terms from all cell types are shown.

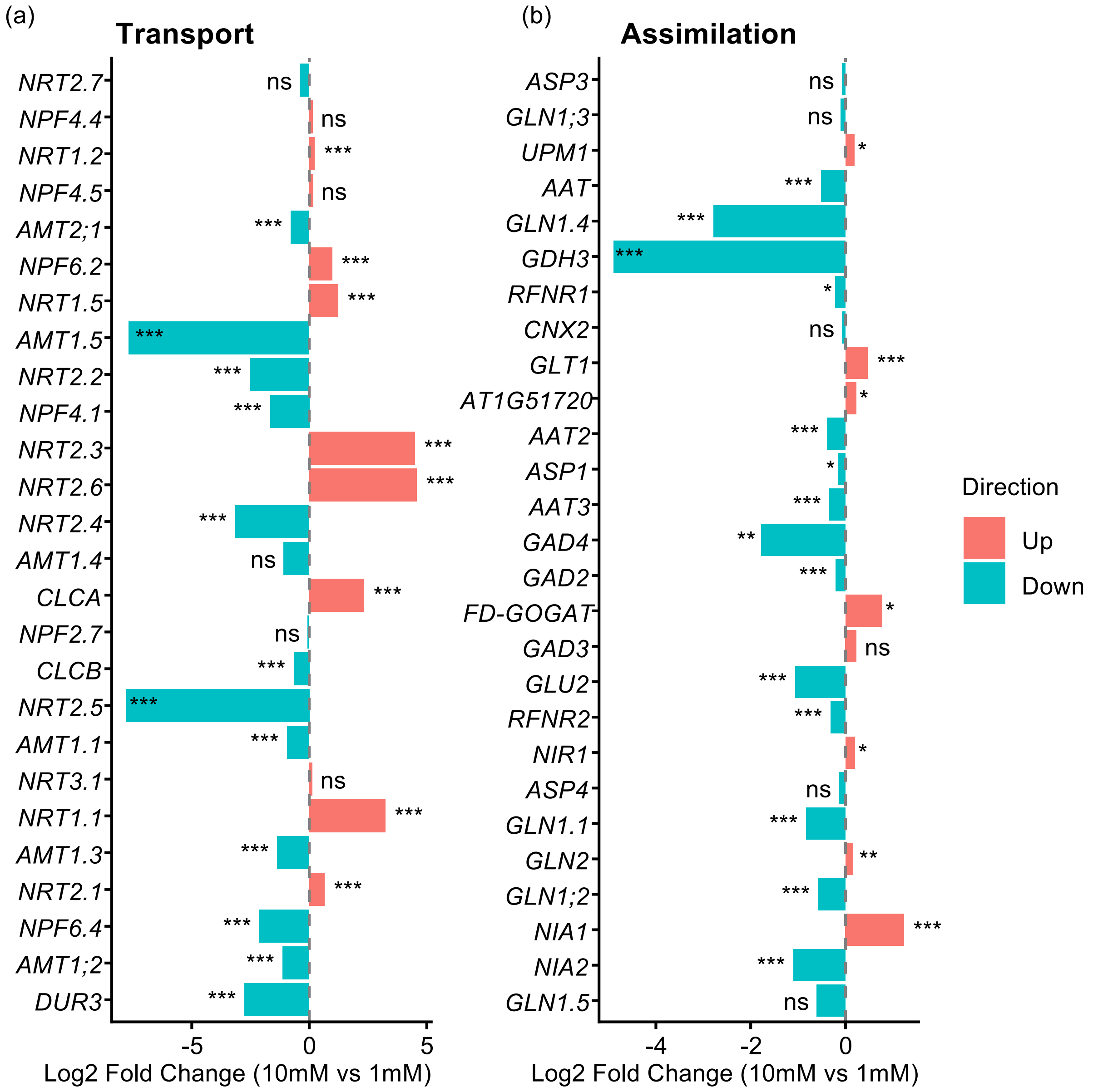


**Supplementary Figure S13. Changes in the expression of nitrogen transport and assimilation related genes under different nitrate conditions in bulk RNAseq of Arabidopsis root.** Log2 fold changes of gene expression in high vs low $KNO_3$ conditions were calculated by DESeq2. ns: p.adj > 0.05, * p.adj < 0.05, ** p.adj < 0.01, *** p.adj < 0.001.

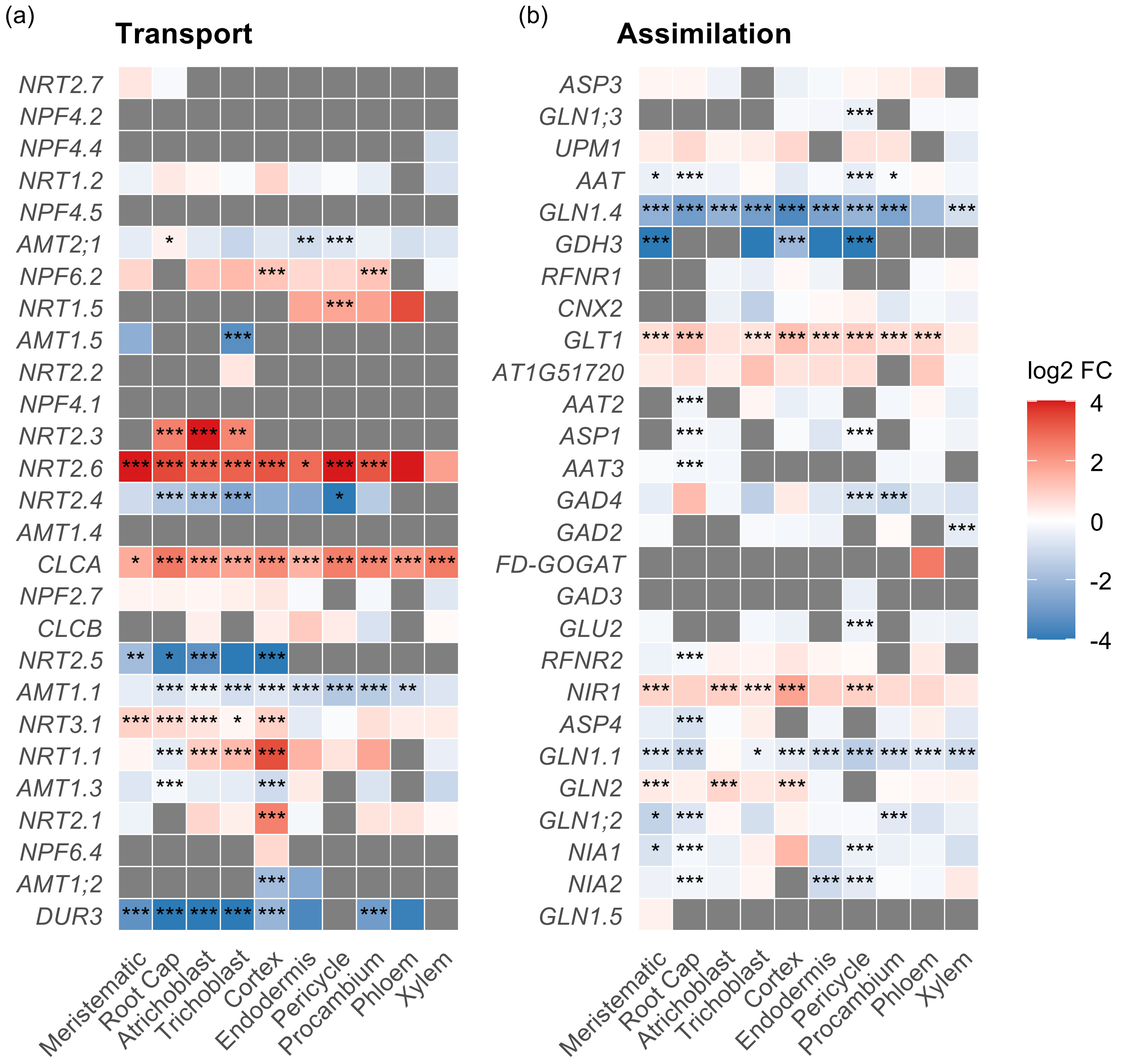


**Supplementary Figure S14. Changes in the expression of nitrogen transport and assimilation related genes under different nitrate conditions in scRNAseq of Arabidopsis root cell types.** Log 2 fold changes of genes involved in nitrogen transport **(a)** and nitrogen assimilation **(b)** at high vs low $KNO_3$ conditions in Arabidopsis Col-0 root cell types. Color scale capped at [-4, 4], extreme values shown at boundary colors. Differential expression analysis was performed using two $KNO_3$ conditions as contrasting groups in the Seurat FindMarkers function. Statistical significance was evaluated according to adjusted p values. * p.adj < 0.05, ** p.adj < 0.01, *** p.adj < 0.001.

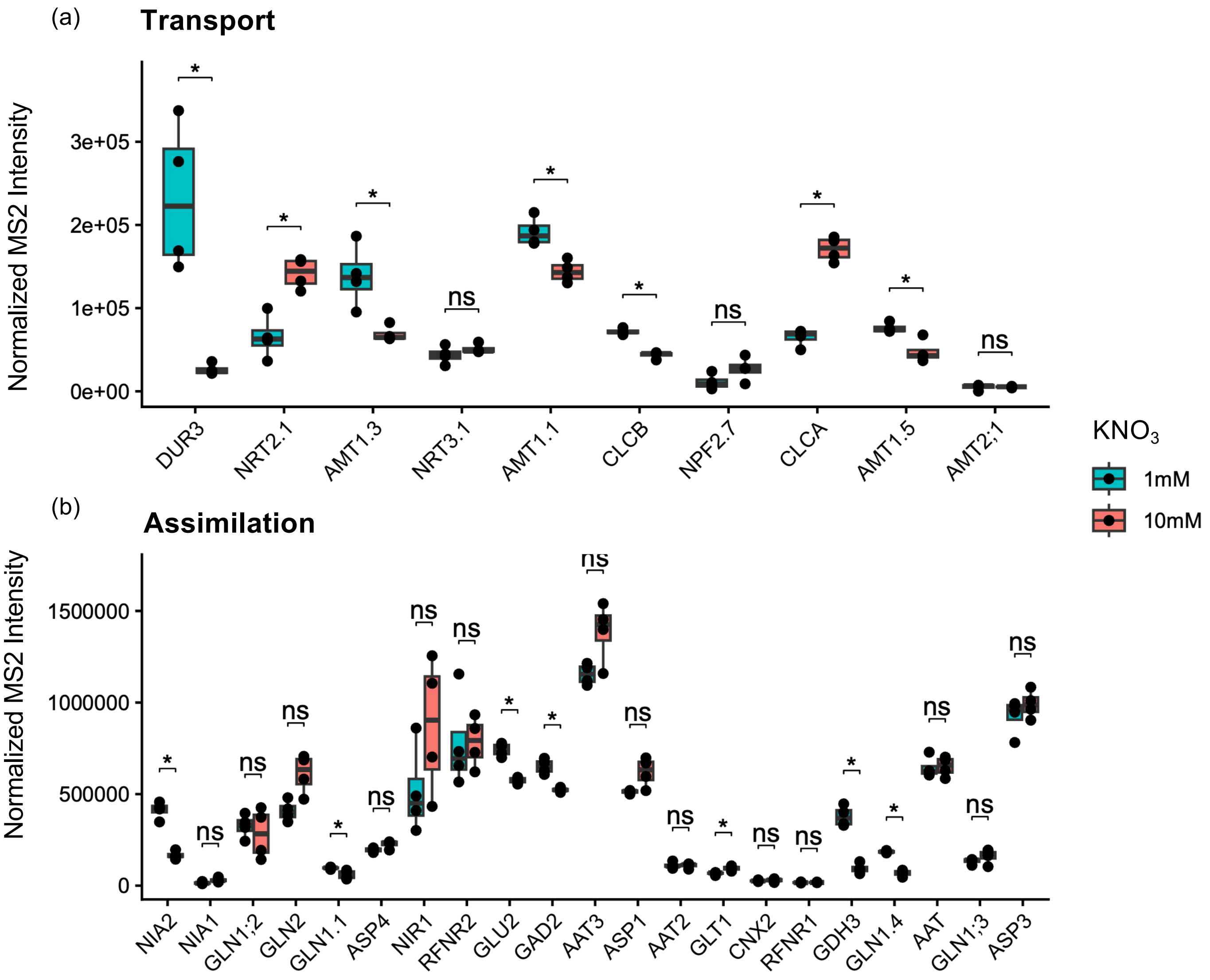


**Supplementary Figure S15. Protein abundance of nitrogen transport and assimilation related genes under different nitrate conditions in the Arabidopsis root. (a)** Detectable N transporters. **(b)** Detectable N-assimilation enzymes. Differences between $KNO_3$ conditions were assessed using the Wilcoxon rank-sum test without multi-comparison correction. ns: $p > 0.05$, * $p < 0.05$. Four biological replicates for each $KNO_3$ condition.

(a)

| Study | Growth condition | Nitrogen form | Treatment | Isolation method | Cell Types |
|---|---|---|---|---|---|
| This study | Solid media plate. 1 x modified MS with 1% (w/v) sucrose, and 1 mM or 10 mM $KNO_3$ as the only nitrogen soure. 10 days. | potassium nitrate | Roots were harvested ~10 days post germination growing on 1 mM or 10 mM nitrate plates. | Protoplast isolation followed by single-cell RNAseq | All root cell types. |
| Gifford et al. 2008 | Hydroponics. 1× MS, 3 mM sucrose and 0.5 mM ammonium succinate. 12 days. | potassium nitrate | Roots were then treated with 5 mM potassium nitrate or mock (KCl) for a total of 3.5 h and then isolated by FACS after a rapid enzymatic dissociation of cells. | Protoplast isolation followed by FACS sorting | lateral root cap, epidermis/cortex, endodermis/pericycle, stele, and xylem pole pericycle |
| Walker et al. 2017 | Solid media plate. 1× MS, 30 mM sucrose, 1.5 mM $CaCl_2$, 0.75 mM $MgSO_4$, 0.625 mM $KH_2PO_4$ , and 0.3 mM $NH_4NO_3$, pH 5.7. 9 days. | ammonium nitrate | Plants were moved to fresh 0.3 mM $NH_4NO_3$ plates or 5 mM $NH_4NO_3$ plates. Whole roots were harvested for protoplast generation and FACS over a 48-h growth period starting at dawn. | Protoplast isolation followed by FACS sorting | cortex and xylem pole pericycle |
| Contreras-López et al. 2022 | Hydroponics in media containing 0.5 mM ammonium succinate for 2 weeks. | potassium nitrate | Roots were treated with 5 mM potassium nitrate or mock (KCl), and harvested for protoplast generation and FACS after 12, 20, 60, and 120 min. | Protoplast isolation followed by FACS sorting | Epidermis, cortex, endodermis (incl. QC), xylem-pole pericycle, and stele |

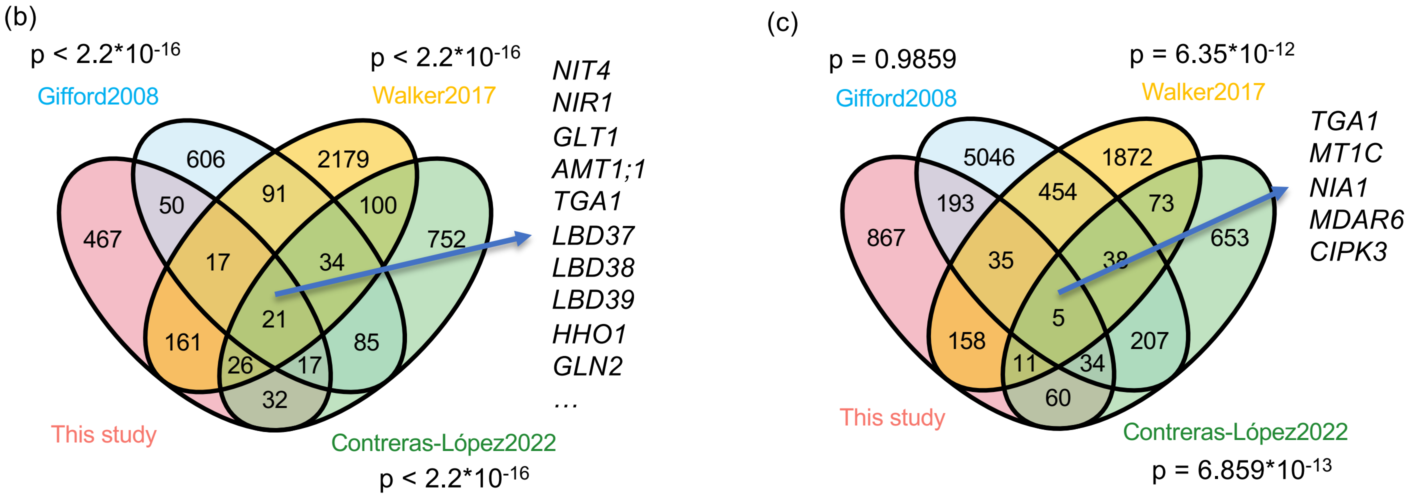


**Supplementary Figure S16. Comparison with previous Arabidopsis cell-type-specific nitrogen response studies. (a)** Comparison of experimental conditions. **(b)** Venn plot comparing differentially expressed genes (DEGs) found in the cortex in this study, Walker et al. 2017 study, Contreras-López et al. 2022 study, and the epidermis/cortex group in the Gifford et al. 2008 study. **(c)** Venn plot comparing differentially expressed genes (DEGs) found in the xylem-pole-pericycle of four studies. DEG threshold used for the sc study was padj <0.05. Significance of overlaps between this study and previous studies as calculated using Fisher's Exact Test and shown next to the studies. Examples of genes found in all studies are listed on the side.

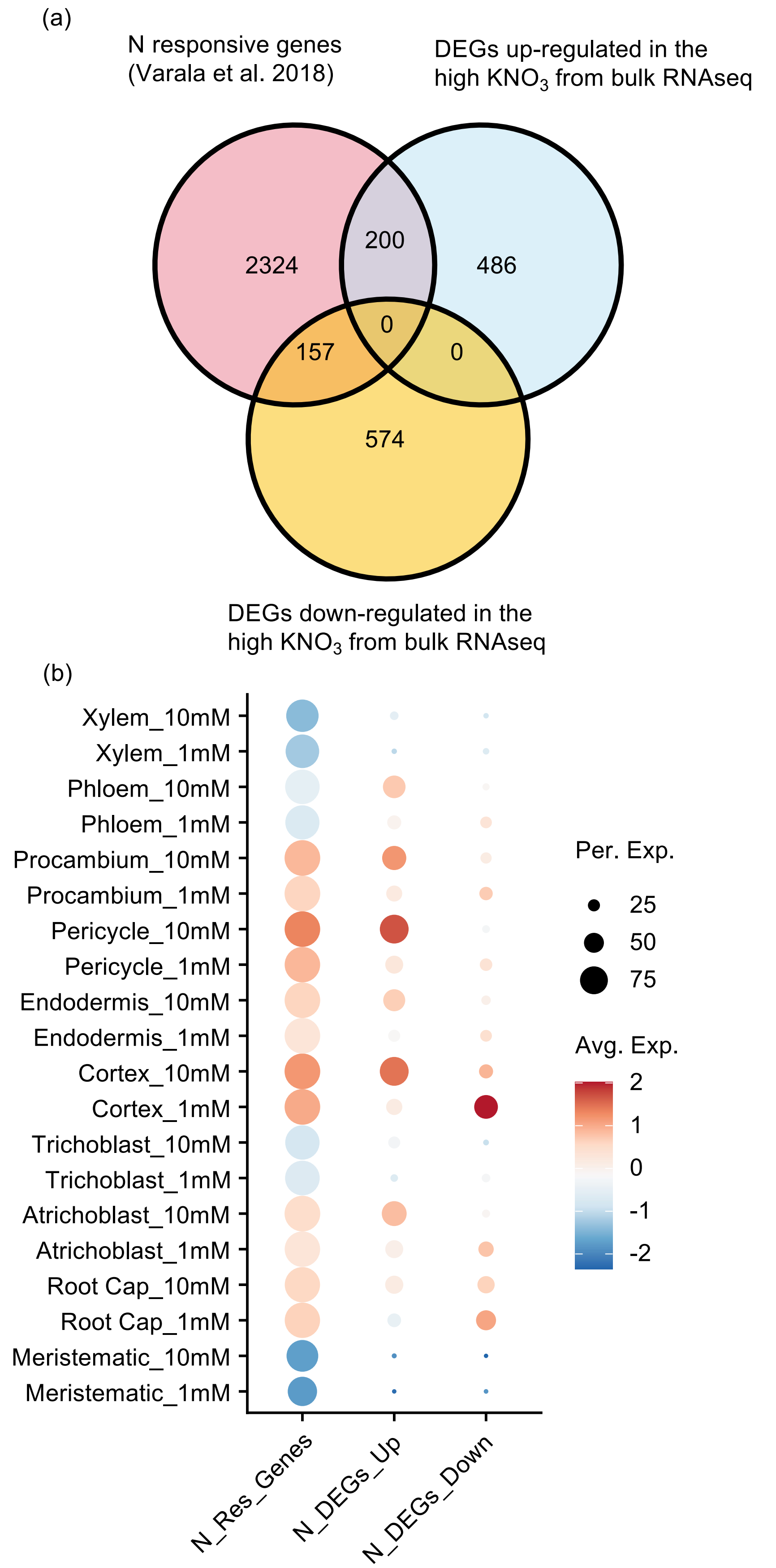


**Supplementary Figure S17. Comparison between N-responsive genes and DEGs from Arabidopsis bulk RNAseq. (a)** Overlaps of the nitrogen(N)-responsive genes discovered from time-course transcriptome responses after $NO_3^-$ and $NH_4^+$ supply (Varala et al. 2018) and differentially expressed genes (DEGs) between high and low $KNO_3$ found in the bulk RNAseq from this study. **(b)** Average expression of N responsive genes from Varala et al. 2018 (N_Res_Genes), and DEGs up- (N_DEGs_Up) and down- (N_DEGs_Down) regulated at high/low $KNO_3$ found in the bulk RNAseq.

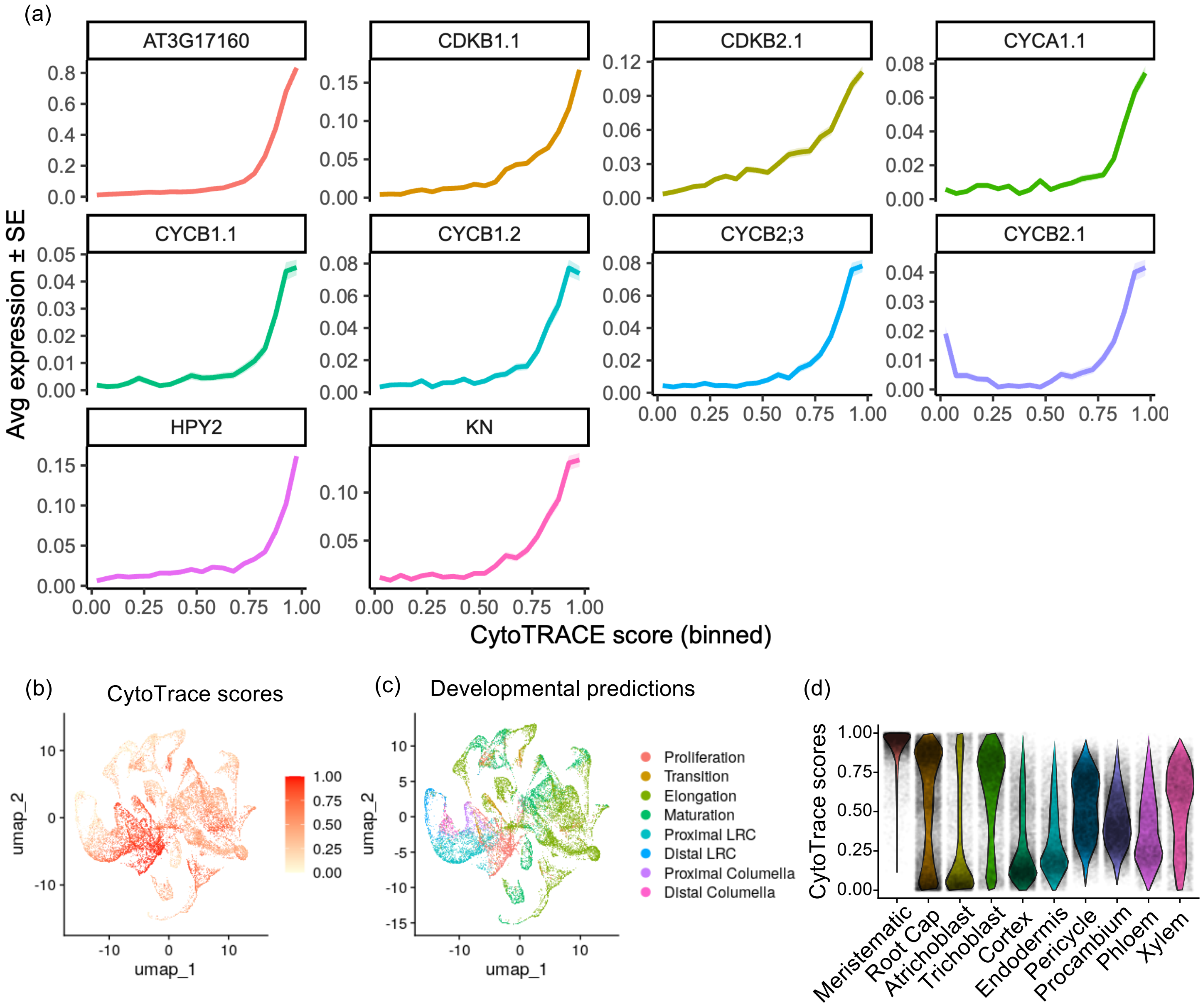


**Supplementary Figure S18. Developmental potential estimation of Arabidopsis root cells using CytoTRACE. (a)** Correlation between expression of Arabidopsis cell cycle genes and CytoTRACE scores (all cells grouped into 20 equal-sized bins). For each bin and gene, mean CytoTRACE score and mean expression (± standard error) were calculated and plotted. **(b, c)** UMAP visualisation of CytoTRACE scores (**b**) and developmental labels transferred from Shahan et al. (2022) Arabidopsis root atlas (**c**).**(d)** Distribution of CytoTRACE scores in each cell type of the Arabidopsis scRNAseq dataset.

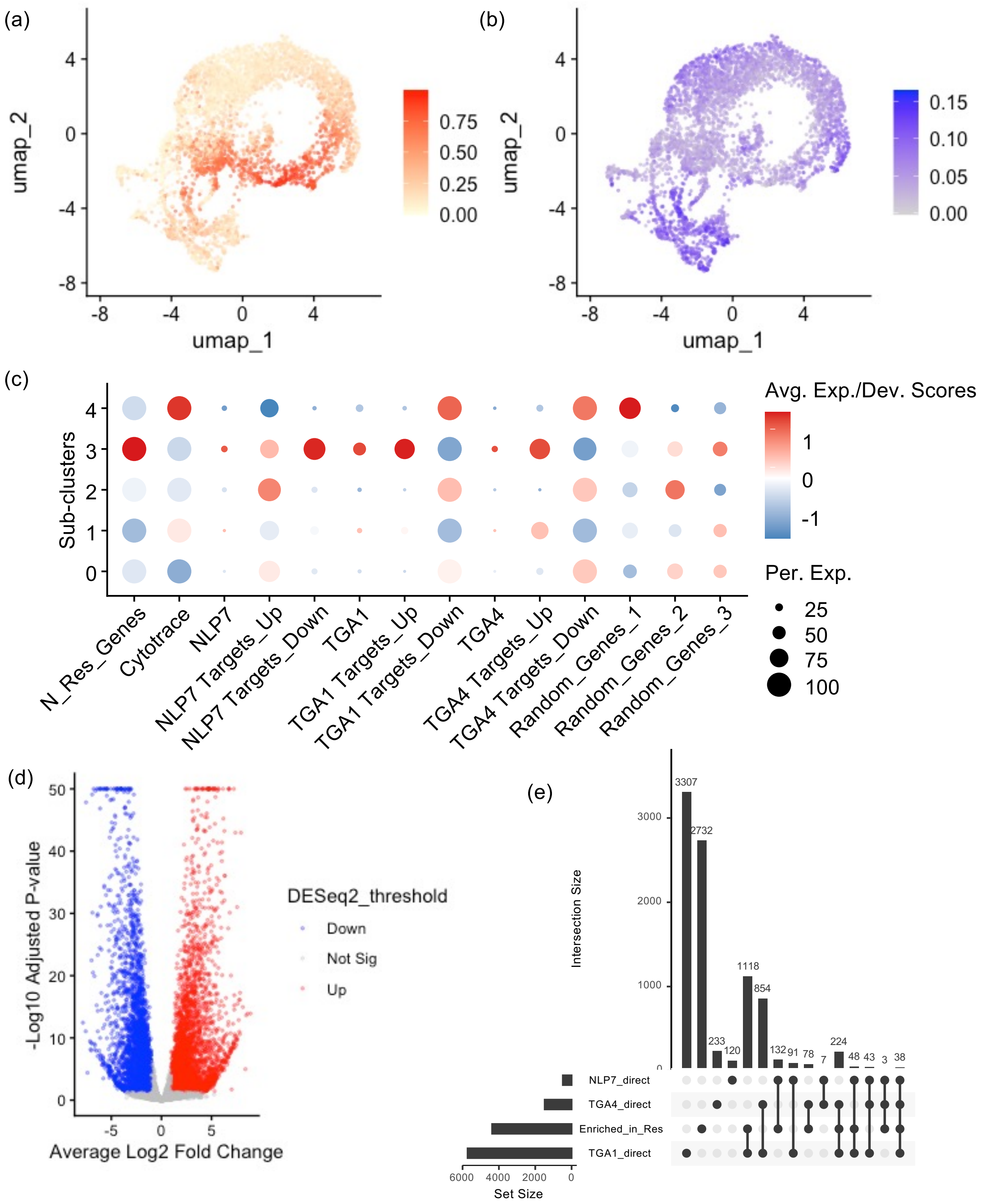


**Supplementary Figure S19. Sub-population analysis of Arabidopsis cortex in the Col-0 WT. (a)** UMAP visualization of CytoTRACE scores in cortex cells. **(b)** UMAP visualization of the average expression of the N-responsive genes (Varala et al. 2018). **(c)** Scaled CytoTRACE scores and average expression of corresponding genes in subclusters. N_Res_Genes, N-responsive genes (Varala et al. 2018). NLP7 Targets_Up/Down, direct targets of AtNLP7 (Alvarez et al. 2020). TGA1/TGA4 Targets_Up/Down, direct targets of AtTGA1/TGA4 (Brooks et al. 2019). Random_Genes_1/2/3, three independent sets of 2652 randomly sampled genes. **(d)** Volcano plots of the differently expressed gene between responsive (sub-cluster 3) and non-responsive, mature (sub-clusters 0 and 2) sub-clusters. Red dots indicate genes that are significantly up-regulated in the responsive sub-population (adjusted p_value < 0.05, log2 fold changes > 1) using DESeq2 on pseudobulked sub-populations. Blue dots for down-regulated genes. **(e)** Overlaps across NLP7/TGA1/TGA4 direct targets and genes that are up-regulated in the responsive sub-population.

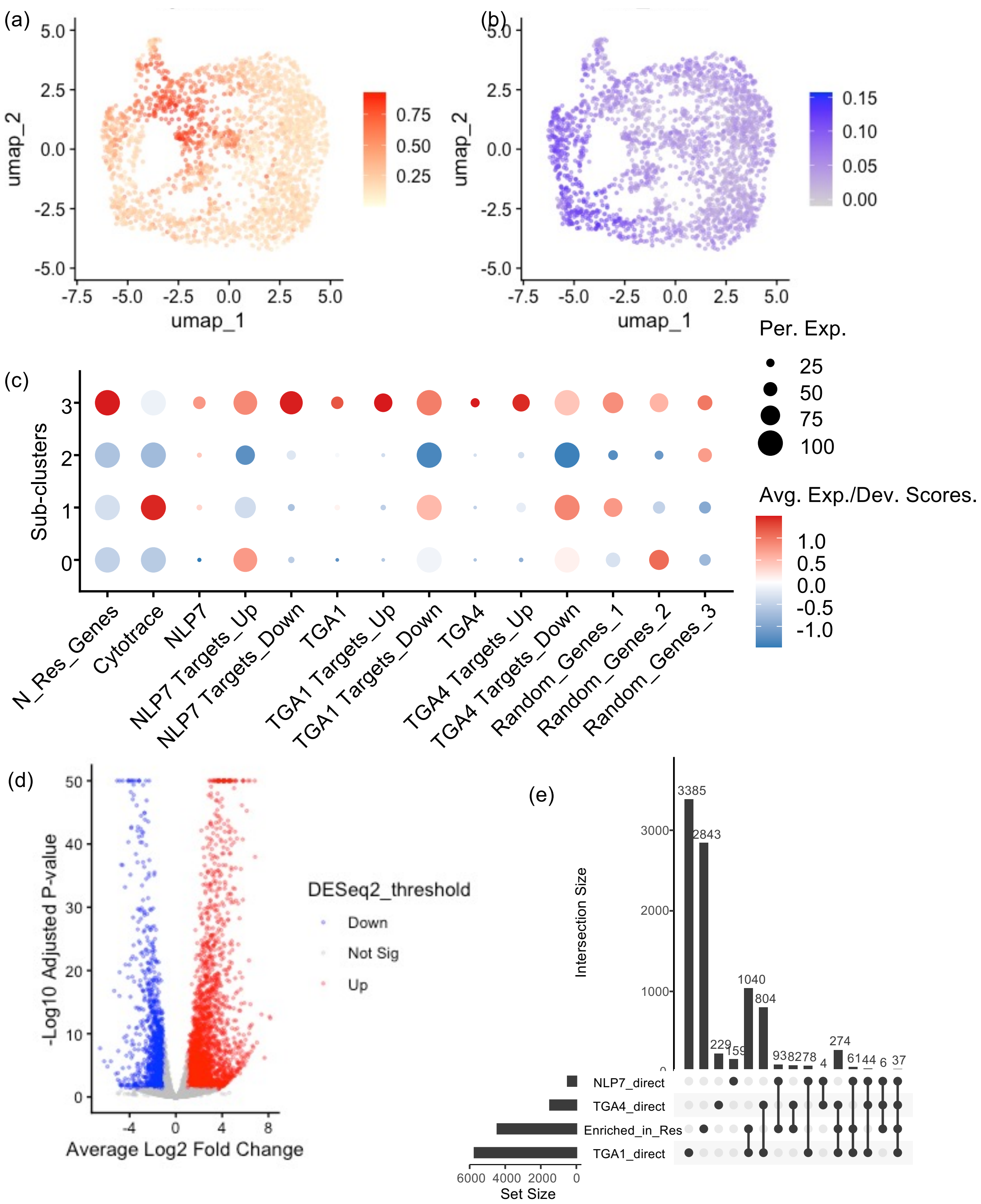


**Supplementary Figure S20. Sub-population analysis of Arabidopsis endodermis in the Col-0 WT. (a)** UMAP visualization of CytoTRACE scores in endodermis cells. **(b)** UMAP visualization of the average expression of the N-responsive genes (Varala et al. 2018). **(c)** Scaled CytoTRACE scores and average expression of corresponding genes in subclusters. N_Res_Genes, N-responsive genes (Varala et al. 2018). NLP7 Targets_Up/Down, direct targets of AtNLP7 (Alvarez et al. 2020). TGA1/TGA4 Targets_Up/Down, direct targets of AtTGA1/TGA4 (Brooks et al. 2019). Random_Genes_1/2/3, three independent sets of 2652 randomly sampled genes. **(d)** Volcano plots of the differently expressed gene between responsive (sub-cluster 3) and non-responsive, mature (sub-clusters 0 and 2) sub-clusters. Red dots indicate genes that are significantly up-regulated in the responsive sub-population (adjusted p_value < 0.05, log2 fold changes > 1) using DESeq2 on pseudobulked sub-populations. Blue dots for down-regulated genes. **(e)** Overlaps across NLP7/TGA1/TGA4 direct targets and genes that are up-regulated in the responsive sub-population.

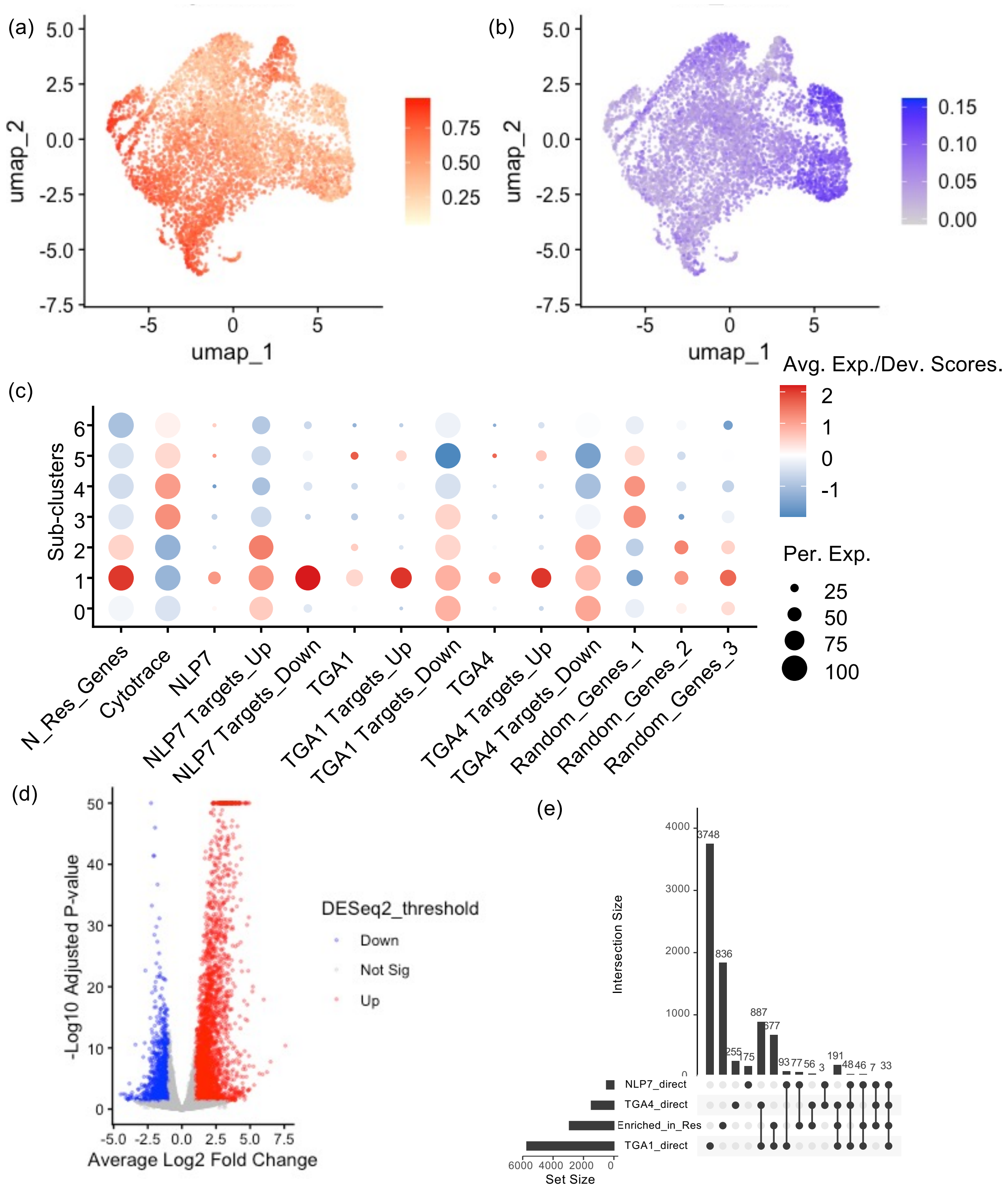


**Supplementary Figure S21. Sub-population analysis of Arabidopsis pericycle in the Col-0 WT.** **(a)** UMAP visualisation of CytoTRACE scores in pericycle cells. **(b)** UMAP visualisation of the average expression of the N-responsive genes (Varala et al. 2018). **(c)** Scaled CytoTRACE scores and average expression of corresponding genes in subclusters. N_Res_Genes, N-responsive genes (Varala et al. 2018). NLP7 Targets_Up/Down, direct targets of AtNLP7 (Alvarez et al. 2020). TGA1/TGA4 Targets_Up/Down, direct targets of AtTGA1/TGA4 (Brooks et al. 2019). Random_Genes_1/2/3, three independent sets of 2652 randomly sampled genes. **(d)** Volcano plots of the differently expressed gene between responsive (sub-cluster 1) and non-responsive, mature (sub-clusters 0 and 2) sub-clusters. Red dots indicate genes that are significantly up-regulated in the responsive sub-population (adjusted p_value < 0.05, log2 fold changes > 1) using DESeq2 on pseudobulked sub-populations. Blue dots for down-regulated genes. **(e)** Overlaps across NLP7/TGA1/TGA4 direct targets and genes that are up-regulated in the responsive sub-population.

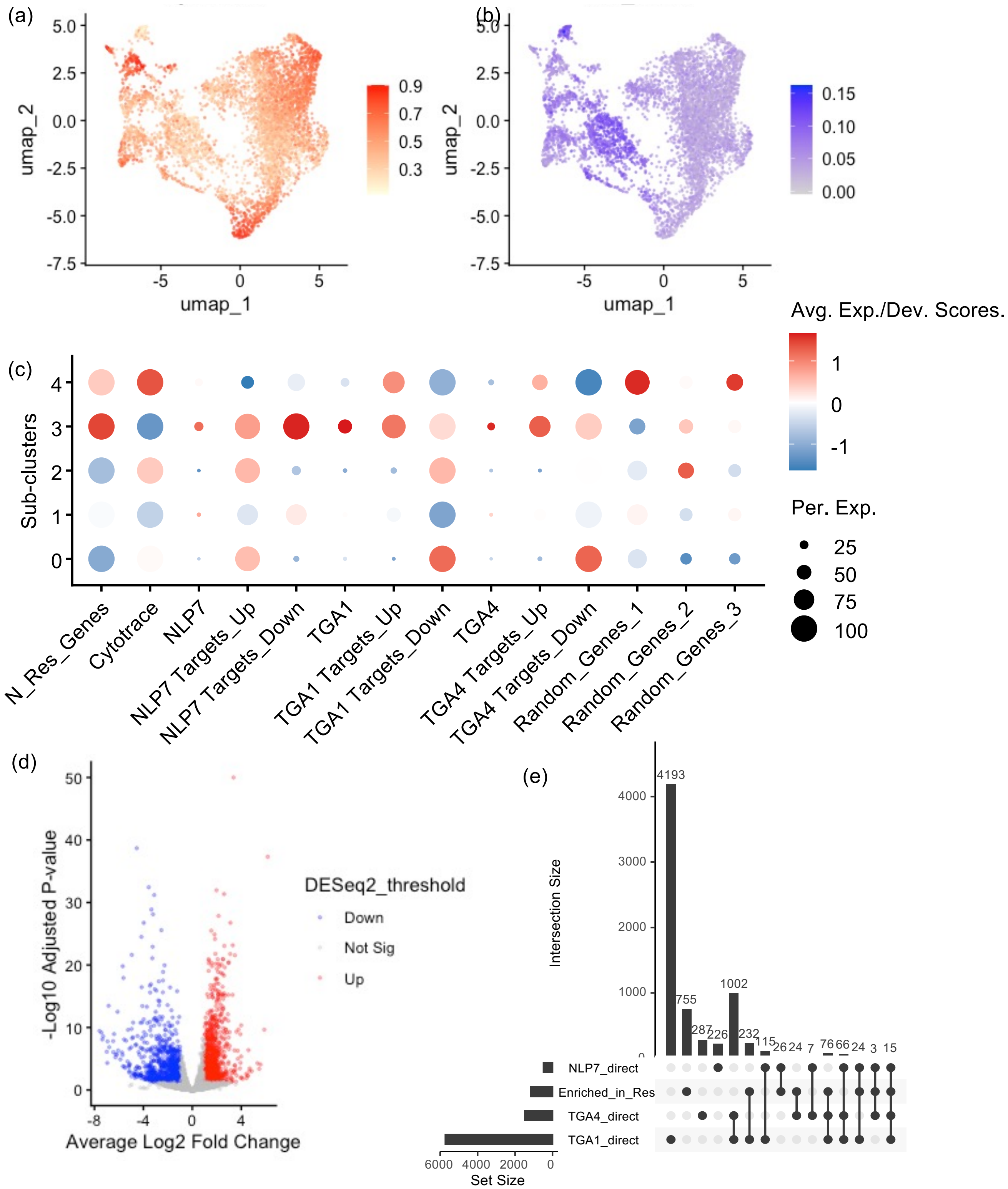


**Supplementary Figure S22. Sub-population analysis of Arabidopsis procambium in the Col-0 WT.** (a) UMAP visualization of CytoTRACE scores in procambium cells. **(b)** UMAP visualization of the average expression of the N-responsive genes (Varala et al. 2018). **(c)** Scaled CytoTRACE scores and average expression of corresponding genes in subclusters. N_Res_Genes, N-responsive genes (Varala et al. 2018). NLP7 Targets_Up/Down, direct targets of AtNLP7 (Alvarez et al. 2020). TGA1/TGA4 Targets_Up/Down, direct targets of AtTGA1/TGA4 (Brooks et al. 2019). Random_Genes_1/2/3, three independent sets of 2652 randomly sampled genes. **(d)** Volcano plots of the differently expressed gene between responsive (sub-cluster 3) and non-responsive, mature (sub-cluster 1) sub-clusters. Red dots indicate genes that are significantly up-regulated in the responsive sub-population (adjusted p_value < 0.05, log2 fold changes > 1) using DESeq2 on pseudobulked sub-populations. Blue dots for down-regulated genes. **(e)** Overlaps across NLP7/TGA1/TGA4 direct targets and genes that are up-regulated in the responsive sub-population.

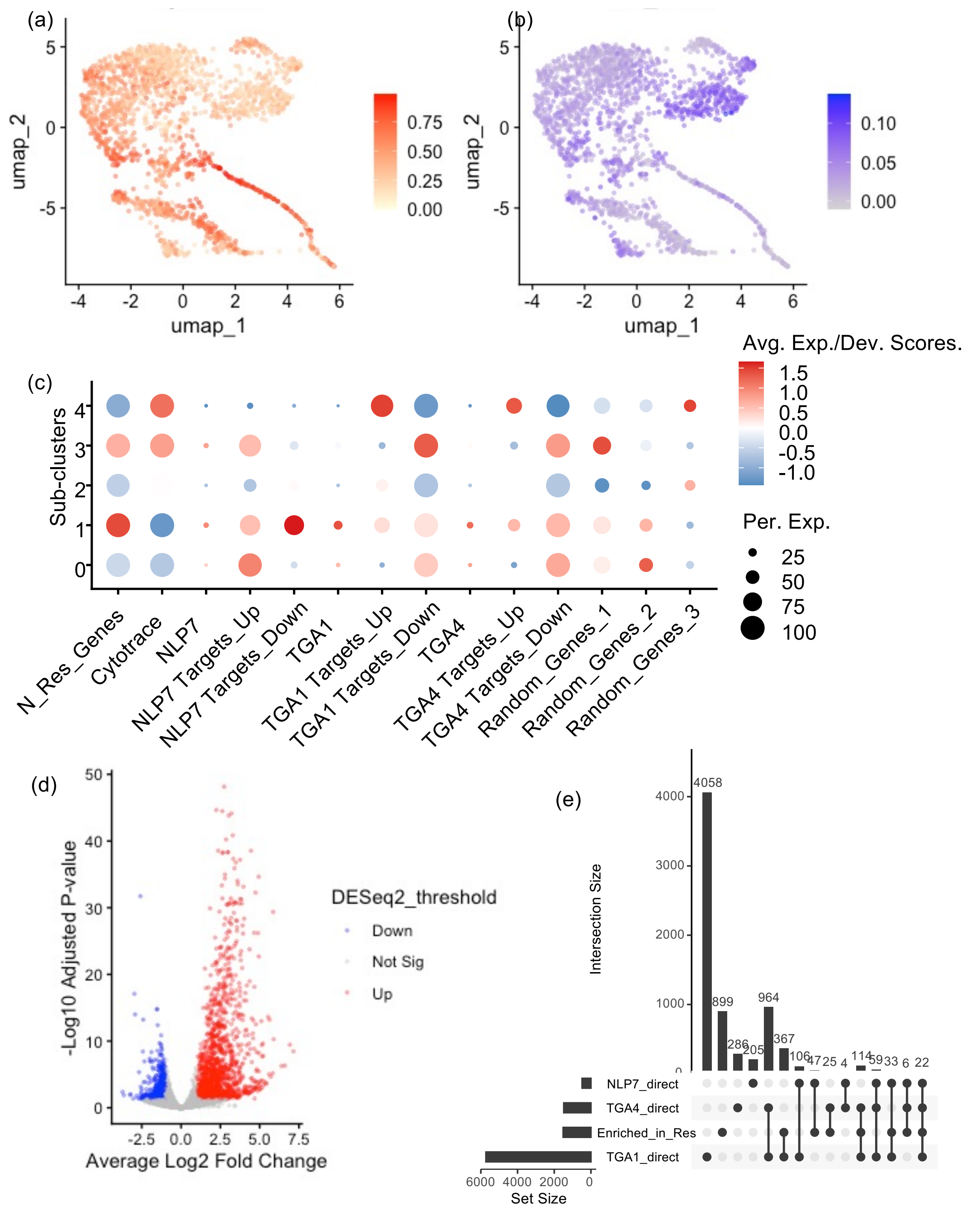


**Supplementary Figure S23. Sub-population analysis of Arabidopsis phloem in the Col-0 WT. (a)** UMAP visualization of CytoTRACE scores in phloem cells. **(b)** UMAP visualization of the average expression of the N-responsive genes (Varala et al. 2018). **(c)** Scaled CytoTRACE scores and average expression of corresponding genes in subclusters. N_Res_Genes, N-responsive genes (Varala et al. 2018). NLP7 Targets_Up/Down, direct targets of AtNLP7 (Alvarez et al. 2020). TGA1/TGA4 Targets_Up/Down, direct targets of AtTGA1/TGA4 (Brooks et al. 2019). Random_Genes_1/2/3, three independent sets of 2652 randomly sampled genes. **(d)** Volcano plots of the differently expressed gene between responsive (sub-cluster 1) and non-responsive, mature (sub-cluster 0) sub-clusters. Red dots indicate genes that are significantly up-regulated in the responsive sub-population (adjusted p_value < 0.05, log2 fold changes > 1) using DESeq2 on pseudobulked sub-populations. Blue dots for down-regulated genes. **(e)** Overlaps across NLP7/TGA1/TGA4 direct targets and genes that are up-regulated in the responsive sub-population.

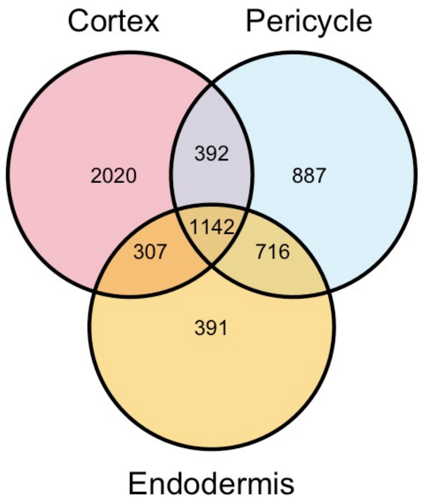


**Supplementary Figure S24. Overlaps of DEGs up-regulated in the responsive sub-populations of Arabidopsis cortex, endodermis and pericycle.**

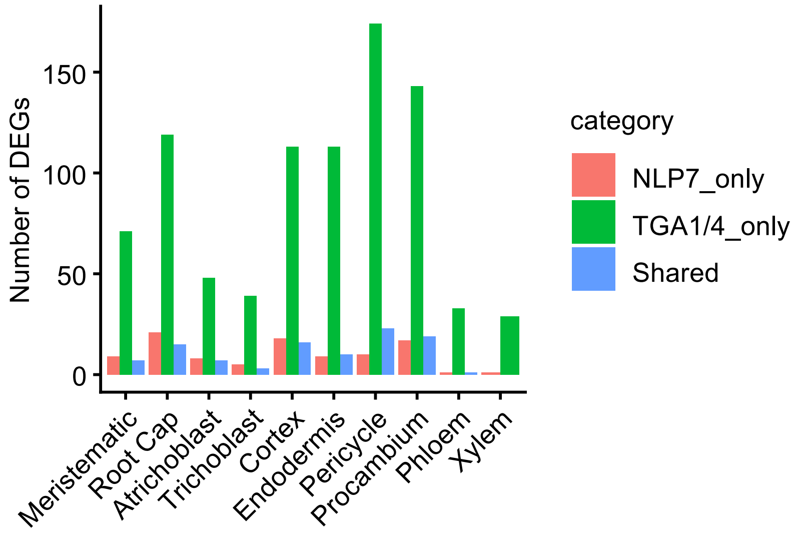


**Supplementary Figure S25. Number of DEGs that are targets of NLP7/TGA1/TGA4 in each cell type of Arabidopsis Col0.** NLP7_only, direct targets of only NLP7. TGA1/4_only, direct targets of only TGA1 or TGA4. Shared, direct targets of all of NLP7, TGA1 and TGA4.

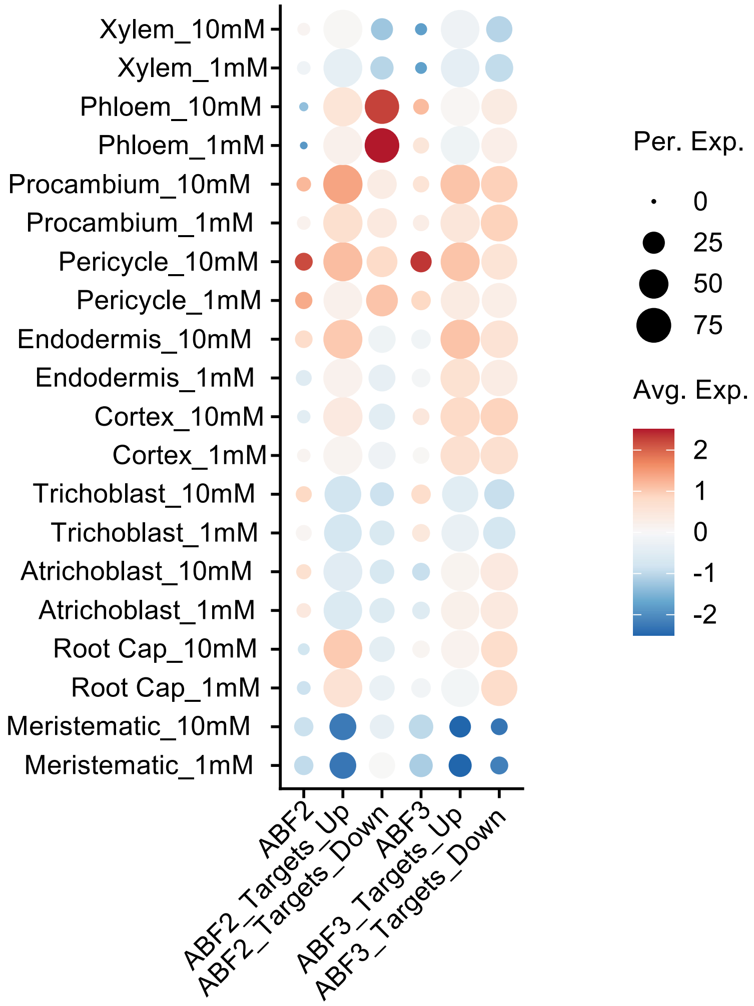


**Supplementary Figure S26. Expression of AtABFs and their target genes in Arabidopsis root cell types.**

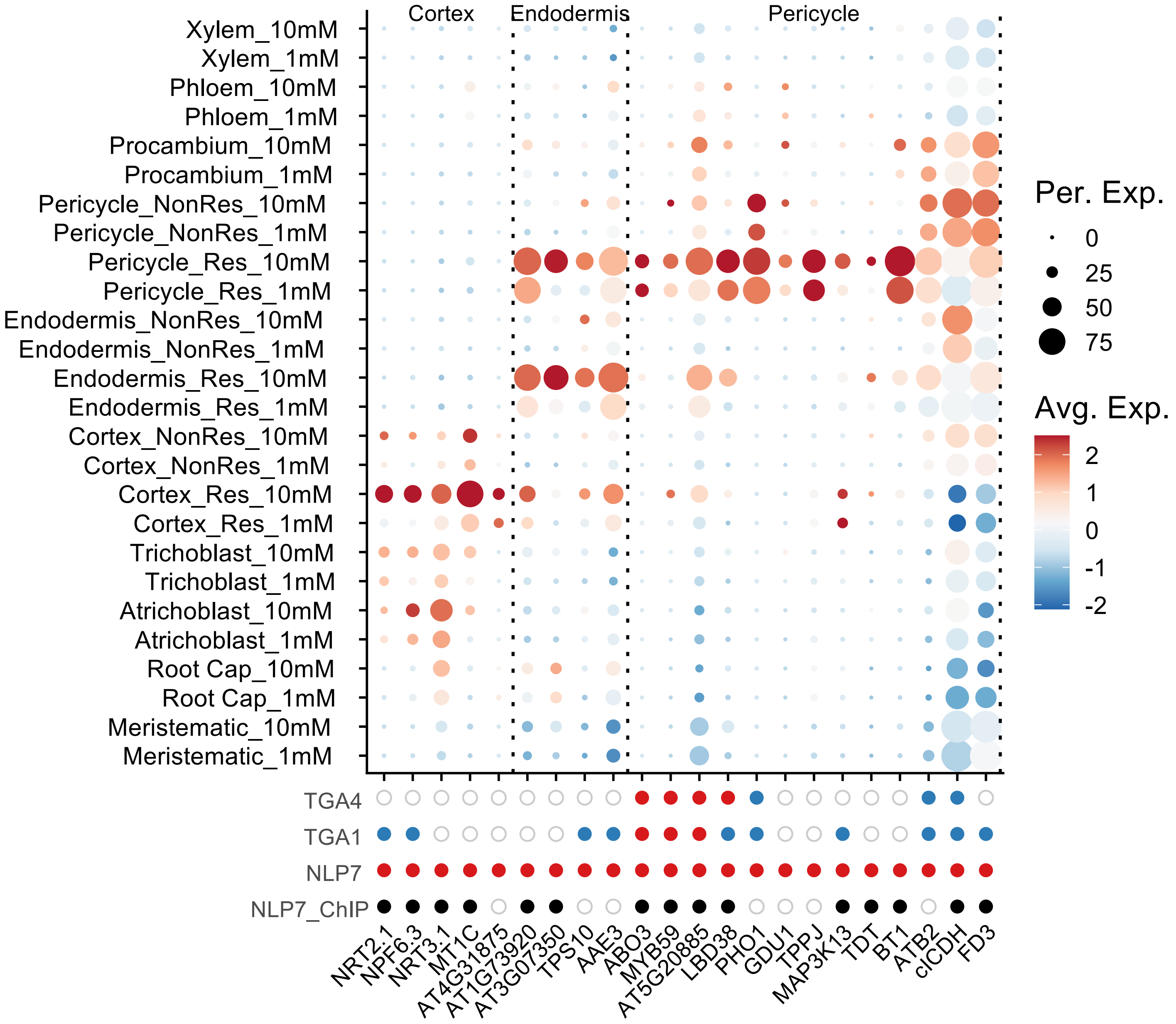


**Supplementary Figure S27. Expression of AtNLP7 targets across all Arabidopsis cell types.** AtNLP7 directly induced targets that show differential expression and cell-type enrichment in cortex, endodermis or pericycle are shown. The same genes are shown in Figure 3d. AtNLP7, AtTGA1 and AtTGA4 targets are indicated at the bottom (red dots indicate genes that are activated by the corresponding TF, blue dots indicate genes that are repressed, empty circles indicate genes that are not targeted). AtNLP7 binding from ChIP assay (Marchive et al. 2013) is indicated by solid black dots.

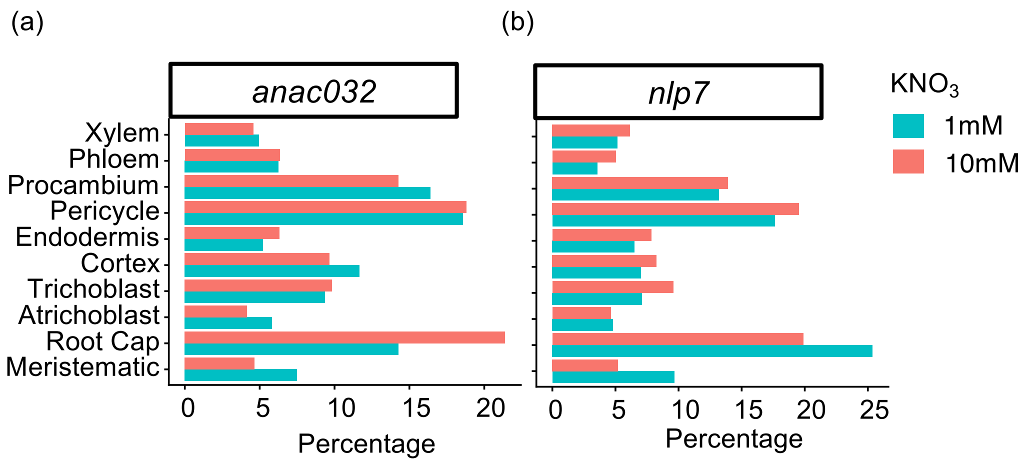


**Supplementary Figure S28. Cell type abundance in the Arabidopsis mutant scRNAseq samples.**
**(a, b)** Percentage of cell types in *anac032* **(a)** and *nlp7* **(b)**.

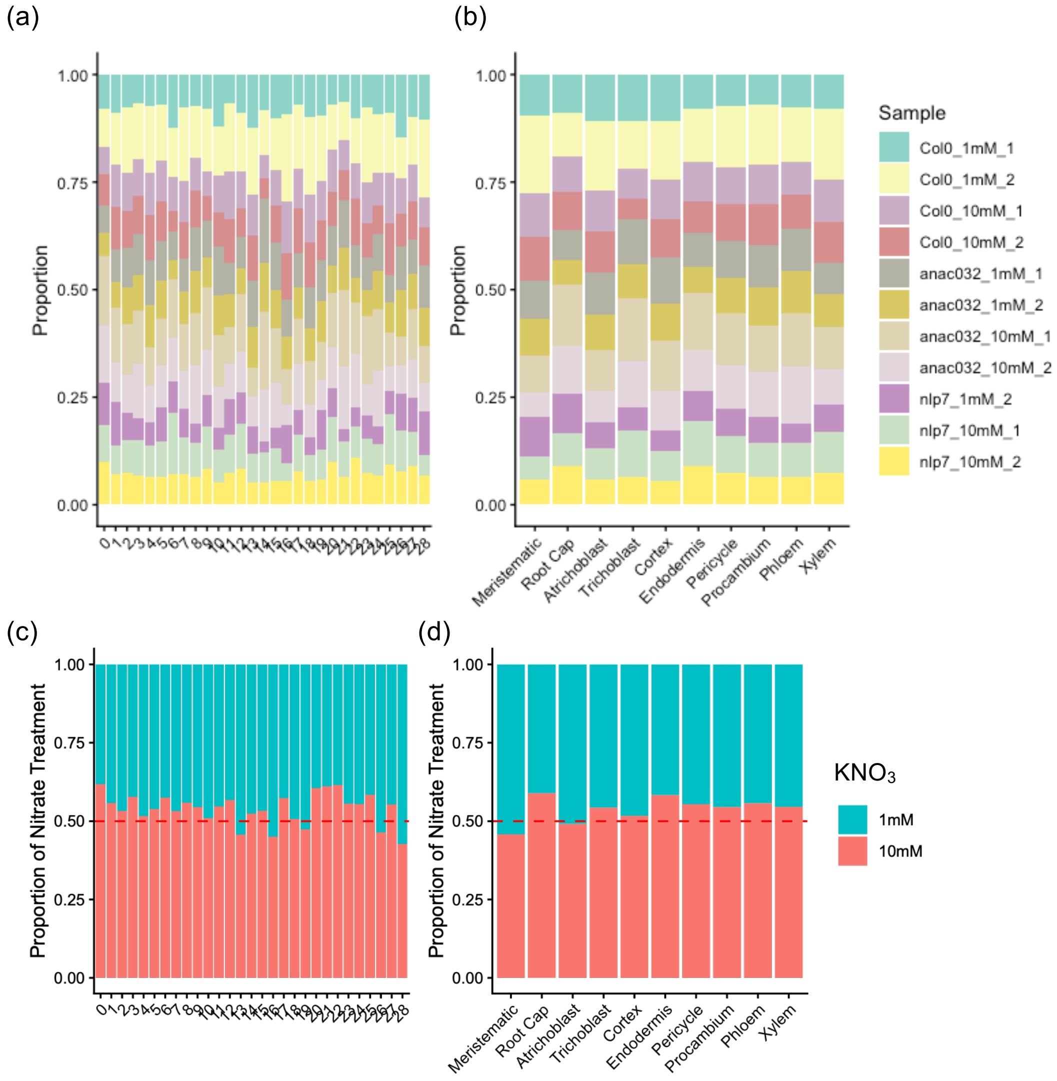


**Supplementary Figure S29. Distribution of samples in Arabidopsis scRNAseq clusters and annotated cell type. (a, b)** Distribution of samples in each Seurat cluster **(a)** and cell type **(b)**. **(c, d)** Distribution of samples from different $KNO_3$ conditions in each cluster **(c)** and cell type **(d)**.

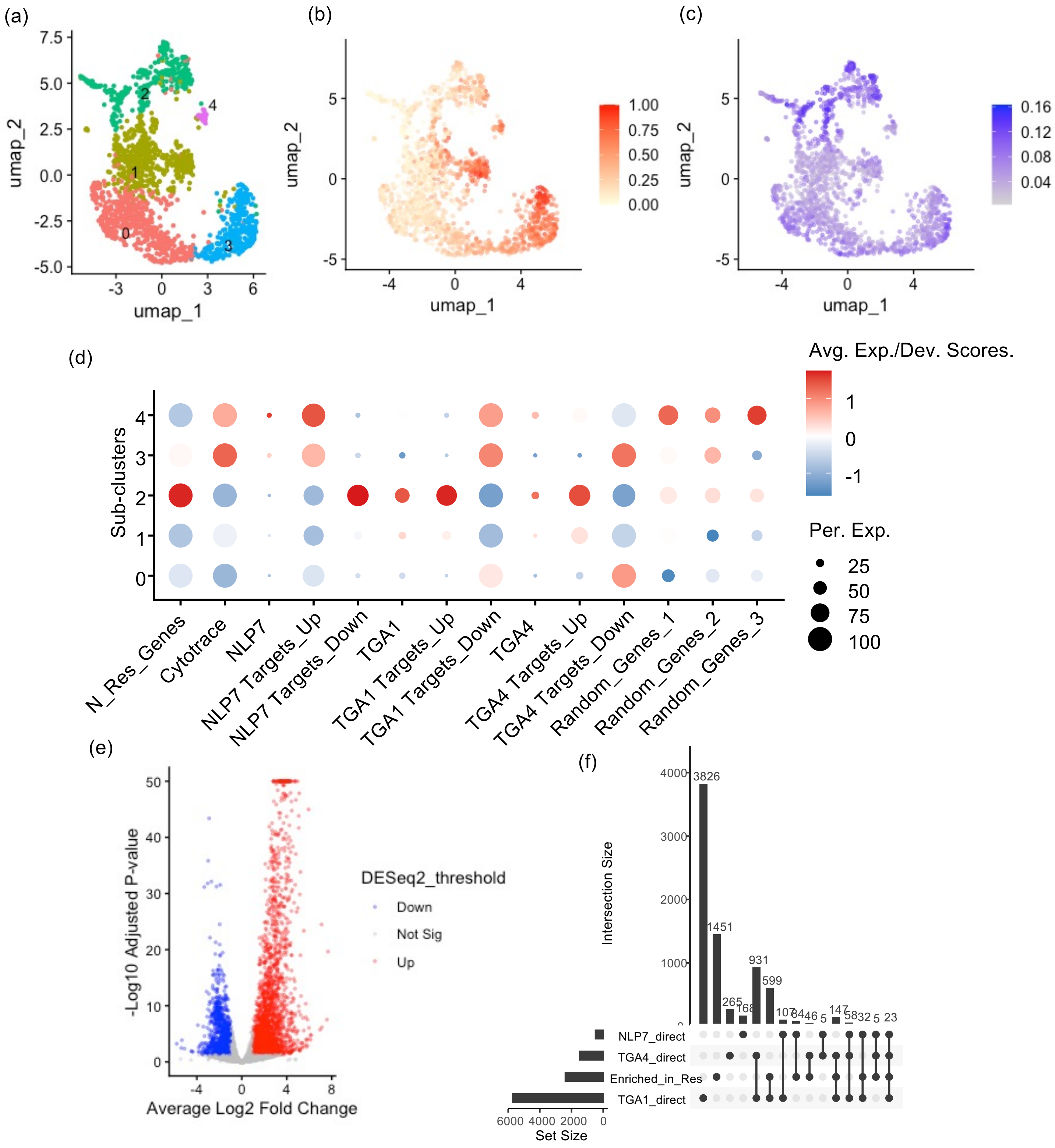


**Supplementary Figure S30. Sub-population analysis of Arabidopsis cortex in the *nlp7* mutant.** **(a)** UMAP visualisation of cortex sub-clusters. **(b)** CytoTRACE scores. **(c)** UMAP visualisation of the average expression of the N-responsive genes (Varala et al. 2018). **(d)** Scaled CytoTRACE scores and average expression of corresponding genes in subclusters. N_Res_Genes, N-responsive genes (Varala et al. 2018). NLP7 Targets_Up/Down, direct targets of AtNLP7 (Alvarez et al. 2020). TGA1/TGA4 Targets_Up/Down, direct targets of AtTGA1/TGA4 (Brooks et al. 2019). Random_Genes_1/2/3, three independent sets of 2652 randomly sampled genes. **(e)** Volcano plots of the differently expressed gene between responsive (sub-cluster 2) and non-responsive (sub-clusters 0 and 1) sub-clusters. Red dots indicate genes that are significantly up-regulated in the responsive non-responsive, mature (adjusted p_value < 0.05, log2 fold changes > 1) using DESeq2 on pseudobulked sub-populations. Blue dots for down-regulated genes. **(f)** Overlaps across NLP7/TGA1/TGA4 direct targets and genes that are up-regulated in the responsive sub-population.

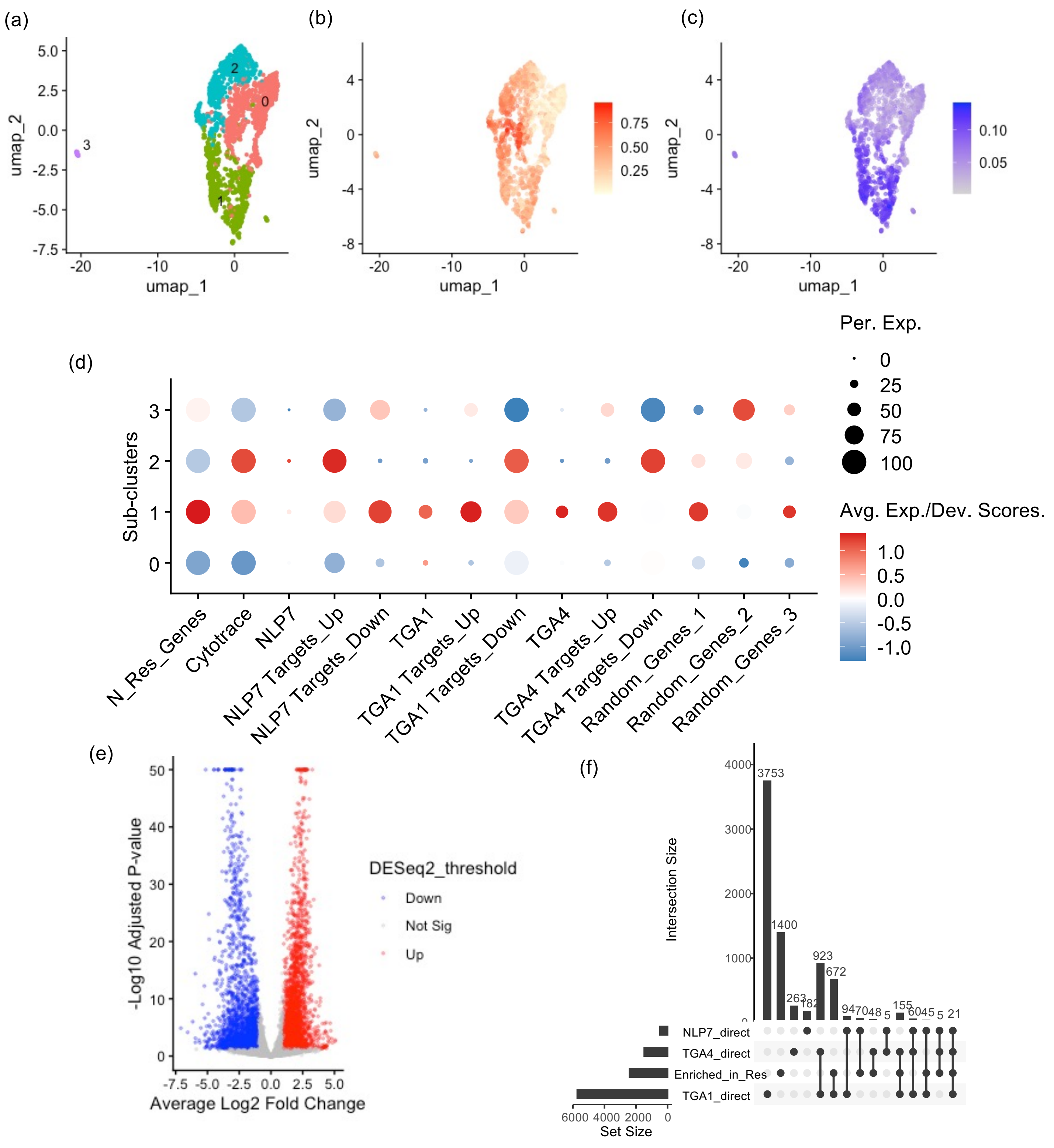


**Supplementary Figure S31. Sub-population analysis of Arabidopsis endodermis in the *nlp7* mutant. (a)** UMAP visualization of endodermis sub-clusters. **(b)** CytoTRACE scores. **(c)** UMAP visualization of the average expression of the N-responsive genes (Varala et al. 2018). **(d)** Scaled CytoTRACE scores and average expression of corresponding genes in subclusters. N_Res_Genes, N-responsive genes (Varala et al. 2018). NLP7 Targets_Up/Down, direct targets of AtNLP7 (Alvarez et al. 2020). TGA1/TGA4 Targets_Up/Down, direct targets of AtTGA1/TGA4 (Brooks et al. 2019). Random_Genes_1/2/3, three independent sets of 2652 randomly sampled genes. **(e)** Volcano plots of the differently expressed gene between responsive (sub-cluster 2) and non-responsive (sub-clusters 0 and 1) sub-clusters. Red dots indicate genes that are significantly up-regulated in the responsive non-responsive, mature (adjusted p_value < 0.05, log2 fold changes > 1) using DESeq2 on pseudobulked sub-populations. Blue dots for down-regulated genes. **(f)** Overlaps across NLP7/TGA1/TGA4 direct targets and genes that are up-regulated in the responsive sub-population.

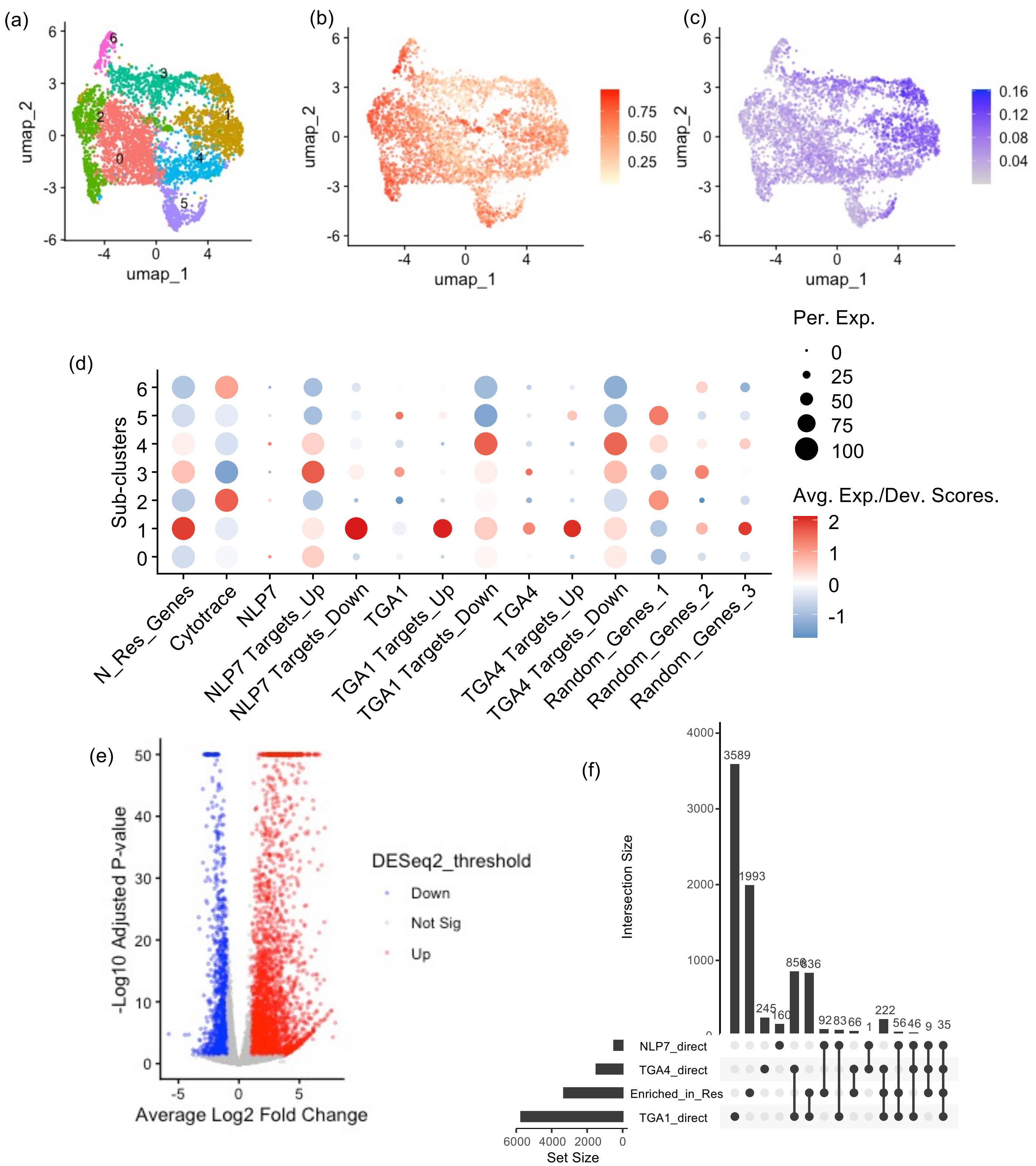


**Supplementary Figure S32. Sub-population analysis of Arabidopsis pericycle in the *nlp7* mutant. (a)** UMAP visualization of pericycle sub-clusters. **(b)** CytoTRACE scores. **(c)** UMAP visualization of the average expression of the N-responsive genes (Varala et al. 2018). **(d)** Scaled CytoTRACE scores and average expression of corresponding genes in subclusters. N_Res_Genes, N-responsive genes (Varala et al. 2018). NLP7 Targets_Up/Down, direct targets of AtNLP7 (Alvarez et al. 2020). TGA1/TGA4 Targets_Up/Down, direct targets of AtTGA1/TGA4 (Brooks et al. 2019). Random_Genes_1/2/3, three independent sets of 2652 randomly sampled genes. **(e)** Volcano plots of the differently expressed gene between responsive (sub-cluster 2) and non-responsive (sub-clusters 0 and 1) sub-clusters. Red dots indicate genes that are significantly up-regulated in the responsive non-responsive, mature (adjusted p_value < 0.05, log2 fold changes > 1) using DESeq2 on pseudobulked sub-populations. Blue dots for down-regulated genes. **(f)** Overlaps across NLP7/TGA1/TGA4 direct targets and genes that are up-regulated in the responsive sub-population.

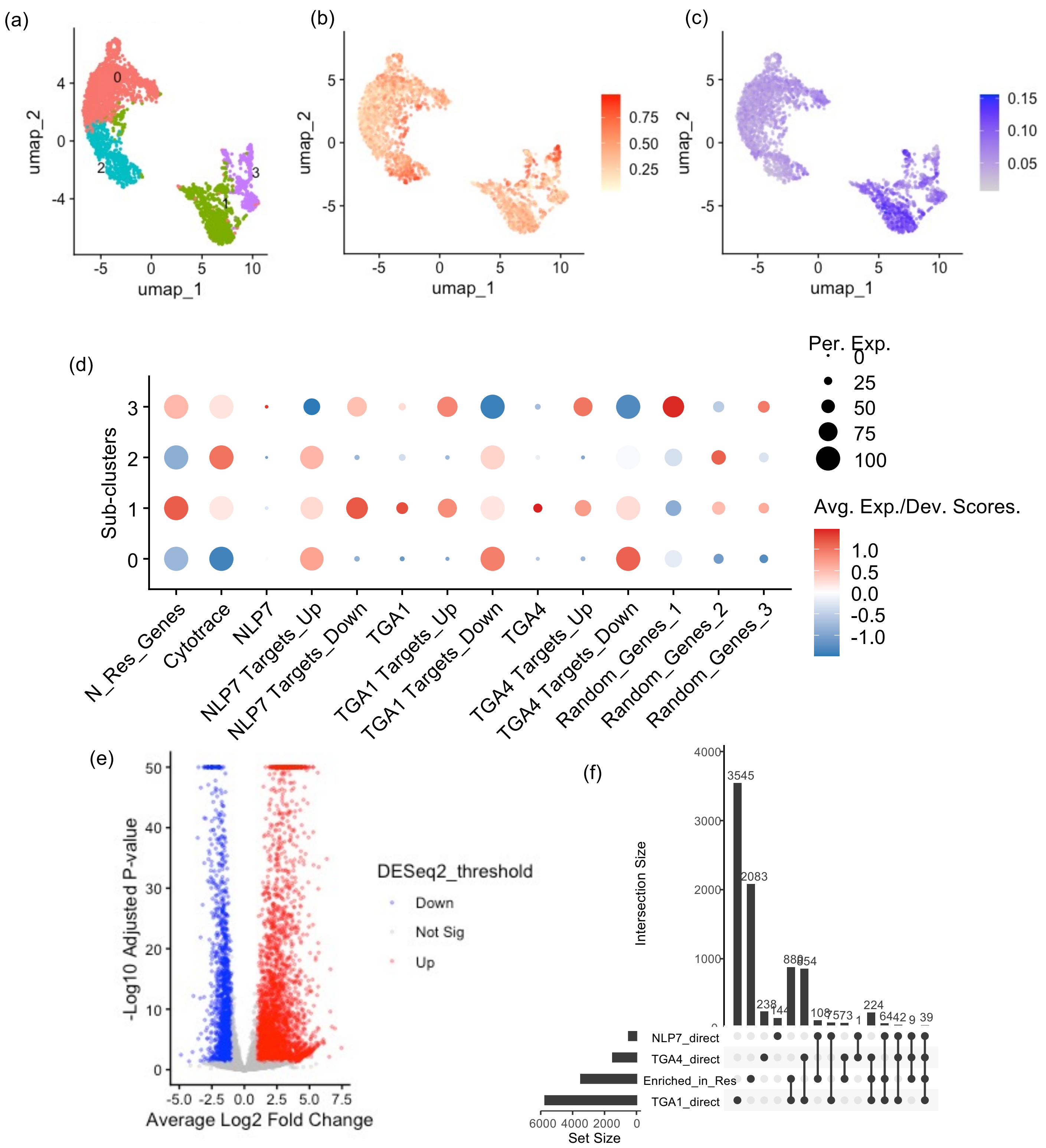


**Supplementary Figure S33. Sub-population analysis of Arabidopsis procambium in the *nlp7* mutant. (a)** UMAP visualization of procambium sub-clusters. **(b)** CytoTRACE scores. **(c)** UMAP visualization of the average expression of the N-responsive genes (Varala et al. 2018). **(d)** Scaled CytoTRACE scores and average expression of corresponding genes in subclusters. N_Res_Genes, N-responsive genes (Varala et al. 2018). NLP7 Targets_Up/Down, direct targets of AtNLP7 (Alvarez et al. 2020). TGA1/TGA4 Targets_Up/Down, direct targets of AtTGA1/TGA4 (Brooks et al. 2019). Random_Genes_1/2/3, three independent sets of 2652 randomly sampled genes. **(e)** Volcano plots of the differently expressed gene between responsive (sub-cluster 2) and non-responsive (sub-clusters 0 and 1) sub-clusters. Red dots indicate genes that are significantly up-regulated in the responsive non-responsive, mature (adjusted p_value < 0.05, log2 fold changes > 1) using DESeq2 on pseudobulked sub-populations. Blue dots for down-regulated genes. **(f)** Overlaps across NLP7/TGA1/TGA4 direct targets and genes that are up-regulated in the responsive sub-population.

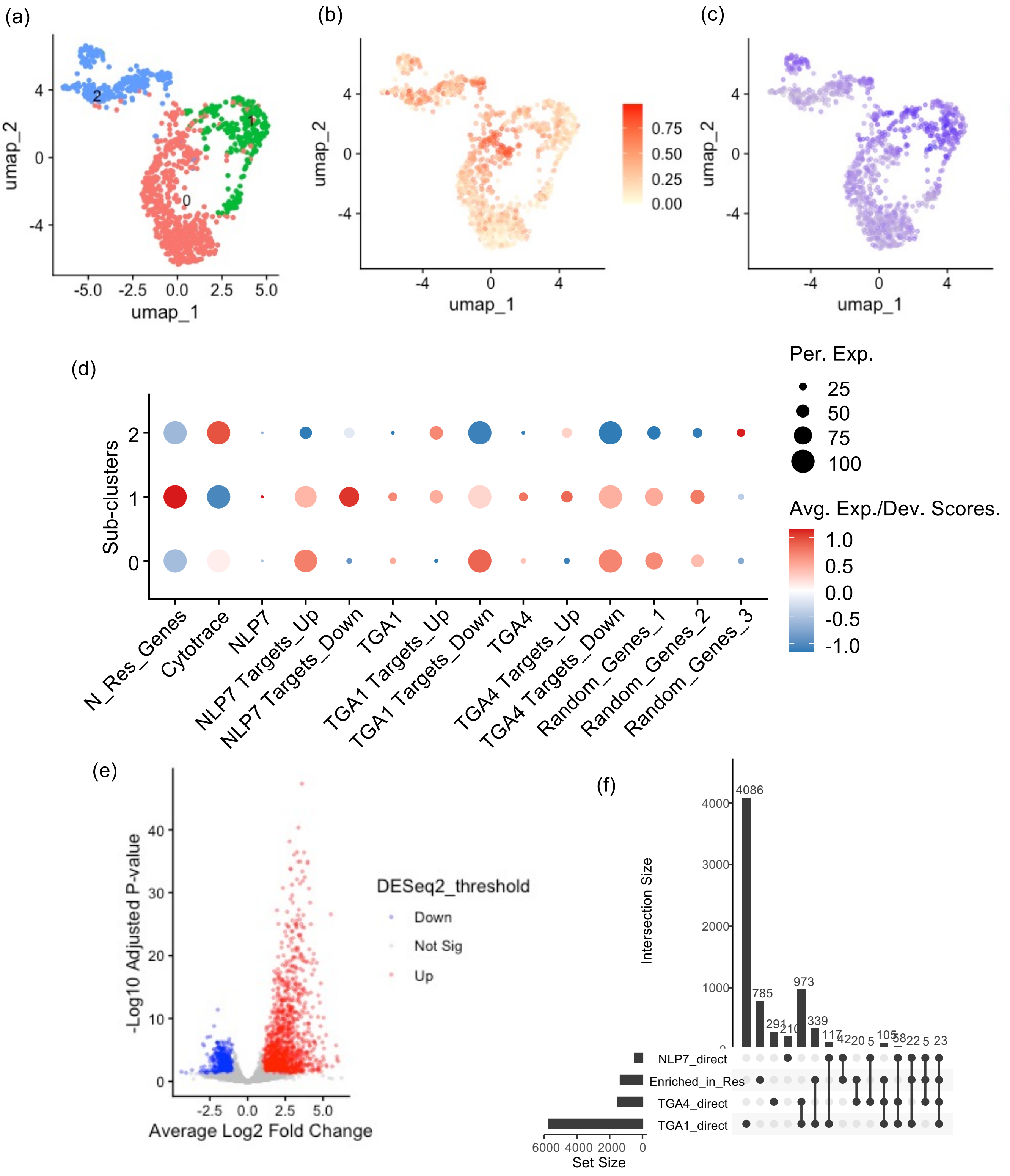


**Supplementary Figure S34. Sub-population analysis of Arabidopsis phloem in the *nlp7* mutant.** **(a)** UMAP visualization of phloem sub-clusters. **(b)** CytoTRACE scores. **(c)** UMAP visualization of the average expression of the N-responsive genes (Varala et al. 2018). **(d)** Scaled CytoTRACE scores and average expression of corresponding genes in subclusters. N_Res_Genes, N-responsive genes (Varala et al. 2018). NLP7 Targets_Up/Down, direct targets of AtNLP7 (Alvarez et al. 2020). TGA1/TGA4 Targets_Up/Down, direct targets of AtTGA1/TGA4 (Brooks et al. 2019). Random_Genes_1/2/3, three independent sets of 2652 randomly sampled genes. **(e)** Volcano plots of the differently expressed gene between responsive (sub-cluster 2) and non-responsive (sub-clusters 0 and 1) sub-clusters. Red dots indicate genes that are significantly up-regulated in the responsive non-responsive, mature (adjusted p_value < 0.05, log2 fold changes > 1) using DESeq2 on pseudobulked sub-populations. Blue dots for down-regulated genes. **(f)** Overlaps across NLP7/TGA1/TGA4 direct targets and genes that are up-regulated in the responsive sub-population.

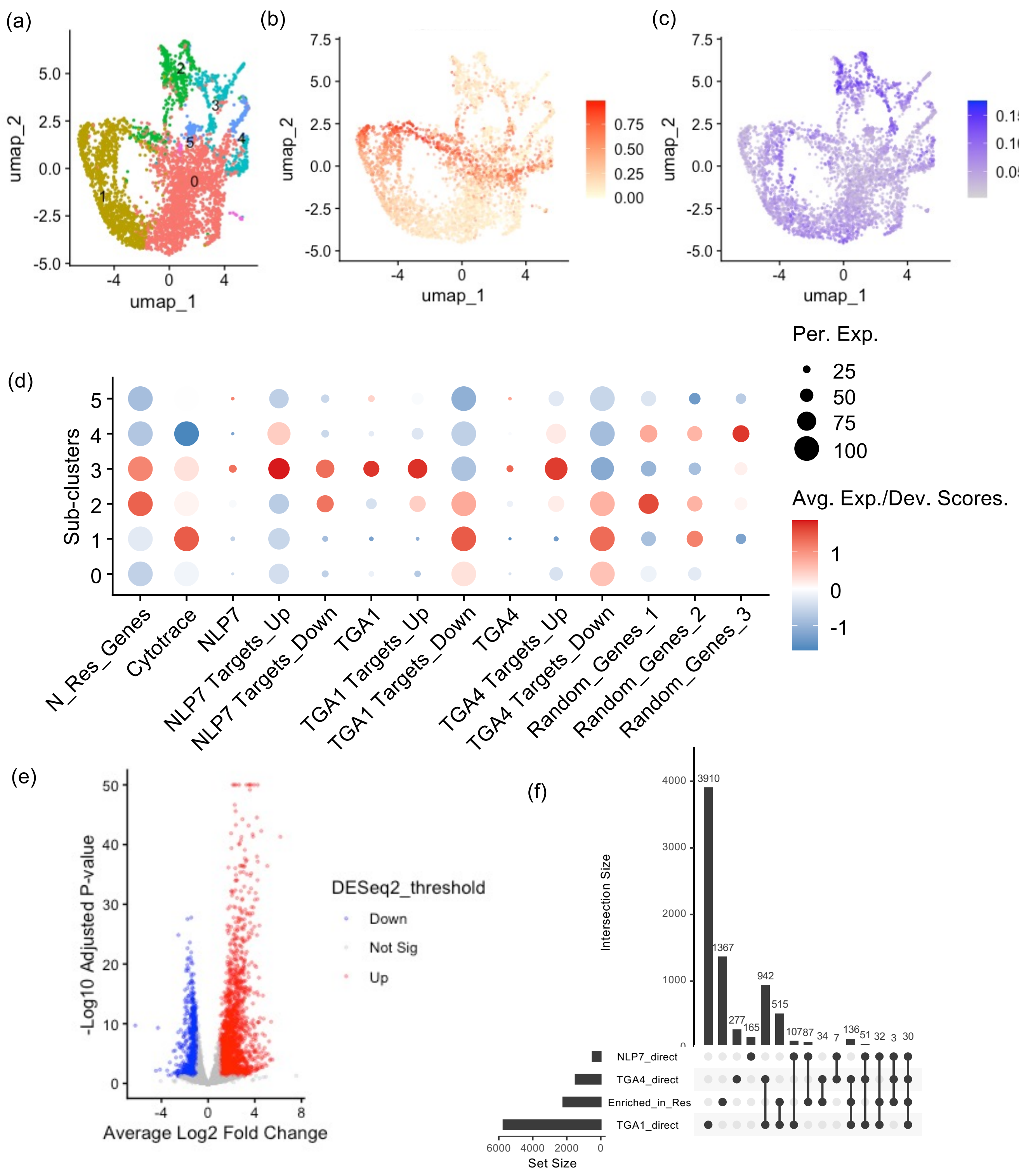


**Supplementary Figure S35. Sub-population analysis of Arabidopsis cortex in the *anac032* mutant. (a)** UMAP visualization of cortex sub-clusters. **(b)** CytoTRACE scores. **(c)** UMAP visualization of the average expression of the N-responsive genes (Varala et al. 2018). **(d)** Scaled CytoTRACE scores and average expression of corresponding genes in subclusters. N_Res_Genes, N-responsive genes (Varala et al. 2018). NLP7 Targets_Up/Down, direct targets of AtNLP7 (Alvarez et al. 2020). TGA1/TGA4 Targets_Up/Down, direct targets of AtTGA1/TGA4 (Brooks et al. 2019). Random_Genes_1/2/3, three independent sets of 2652 randomly sampled genes. **(e)** Volcano plots of the differently expressed gene between responsive (sub-cluster 2) and non-responsive (sub-clusters 0 and 1) sub-clusters. Red dots indicate genes that are significantly up-regulated in the responsive non-responsive, mature (adjusted p_value < 0.05, log2 fold changes > 1) using DESeq2 on pseudobulked sub-populations. Blue dots for down-regulated genes. **(f)** Overlaps across NLP7/TGA1/TGA4 direct targets and genes that are up-regulated in the responsive sub-population.

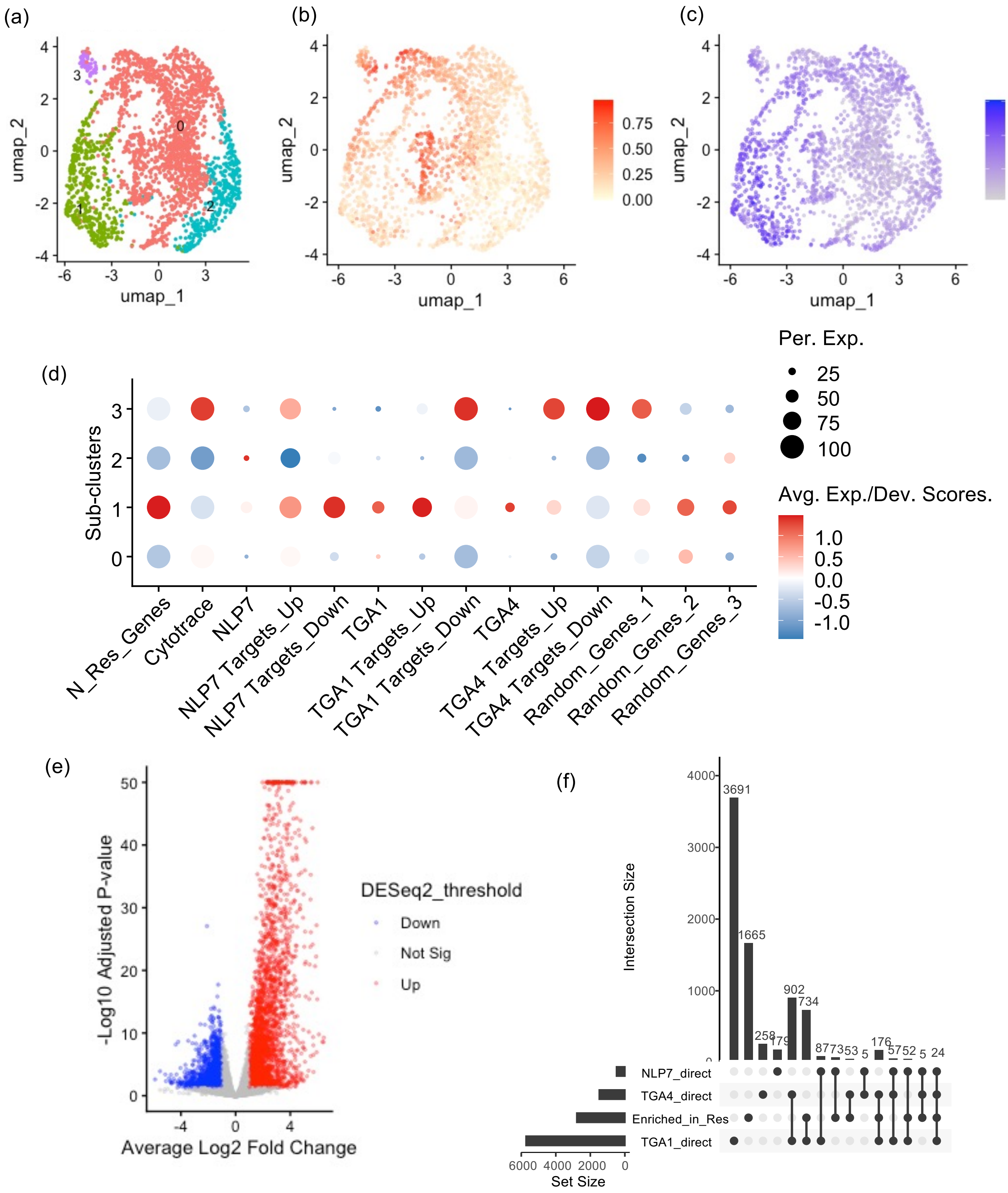


**Supplementary Figure S36. Sub-population analysis of Arabidopsis endodermis in the *anac032* mutant. (a)** UMAP visualization of endodermis sub-clusters. **(b)** CytoTRACE scores. **(c)** UMAP visualization of the average expression of the N-responsive genes (Varala et al. 2018). **(d)** Scaled CytoTRACE scores and average expression of corresponding genes in subclusters. N_Res_Genes, N-responsive genes (Varala et al. 2018). NLP7 Targets_Up/Down, direct targets of AtNLP7 (Alvarez et al. 2020). TGA1/TGA4 Targets_Up/Down, direct targets of AtTGA1/TGA4 (Brooks et al. 2019). Random_Genes_1/2/3, three independent sets of 2652 randomly sampled genes. **(e)** Volcano plots of the differently expressed gene between responsive (sub-cluster 2) and non-responsive (sub-clusters 0 and 1) sub-clusters. Red dots indicate genes that are significantly up-regulated in the responsive non-responsive, mature (adjusted p_value < 0.05, log2 fold changes > 1) using DESeq2 on pseudobulked sub-populations. Blue dots for down-regulated genes. **(f)** Overlaps across NLP7/TGA1/TGA4 direct targets and genes that are up-regulated in the responsive sub-population.

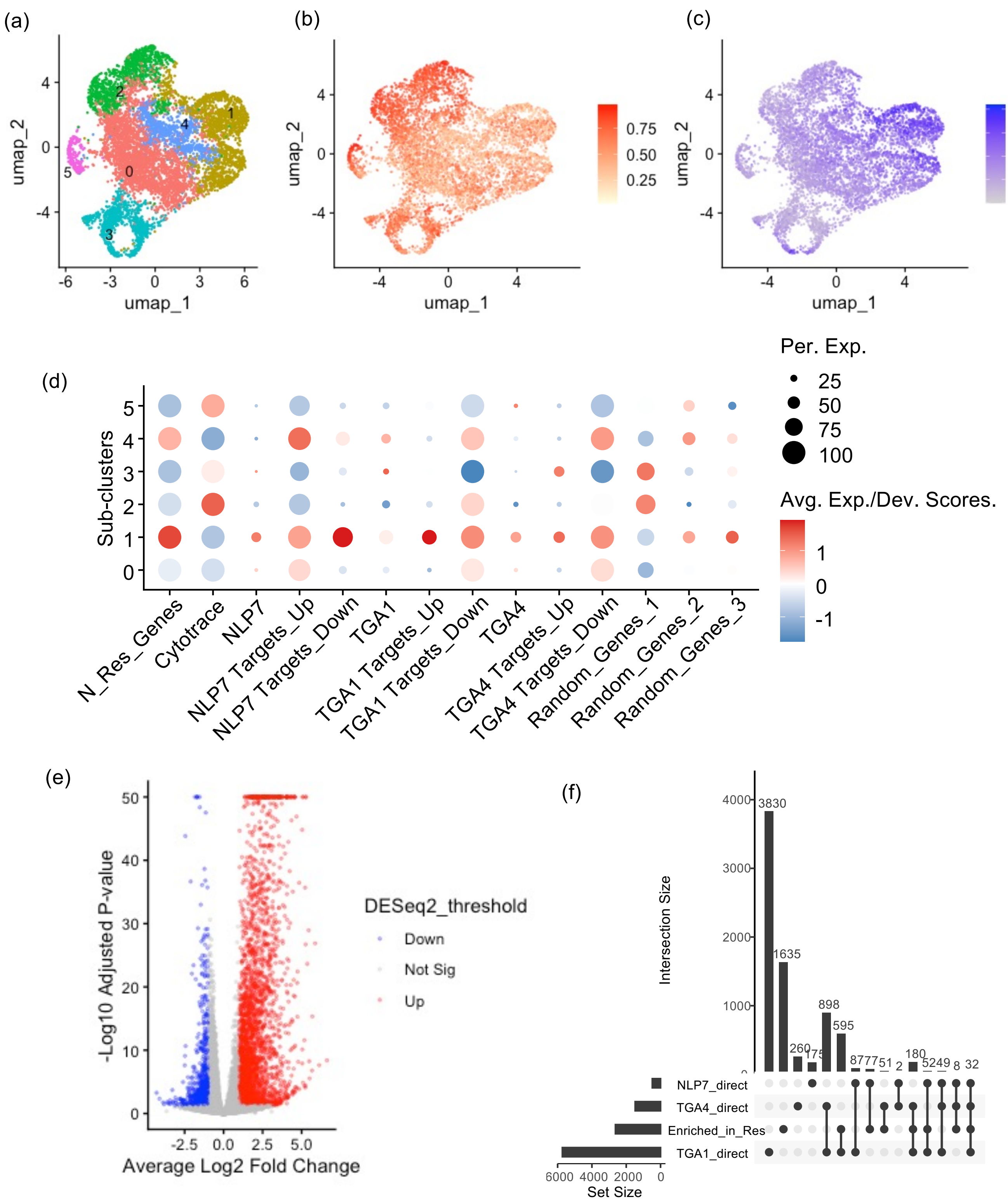

**Supplementary Figure S37. Sub-population analysis of Arabidopsis pericycle in the *anac032* mutant. (a)** UMAP visualization of pericycle sub-clusters. **(b)** CytoTRACE scores. **(c)** UMAP visualization of the average expression of the N-responsive genes (Varala et al. 2018). **(d)** Scaled CytoTRACE scores and average expression of corresponding genes in subclusters. N_Res_Genes, N-responsive genes (Varala et al. 2018). NLP7 Targets_Up/Down, direct targets of AtNLP7 (Alvarez et al. 2020). TGA1/TGA4 Targets_Up/Down, direct targets of AtTGA1/TGA4 (Brooks et al. 2019). Random_Genes_1/2/3, three independent sets of 2652 randomly sampled genes. **(e)** Volcano plots of the differently expressed gene between responsive (sub-cluster 2) and non-responsive (sub-clusters 0 and 1) sub-clusters. Red dots indicate genes that are significantly up-regulated in the responsive non-responsive, mature (adjusted p_value < 0.05, log2 fold changes > 1) using DESeq2 on pseudobulked sub-populations. Blue dots for down-regulated genes. **(f)** Overlaps across NLP7/TGA1/TGA4 direct targets and genes that are up-regulated in the responsive sub-population.

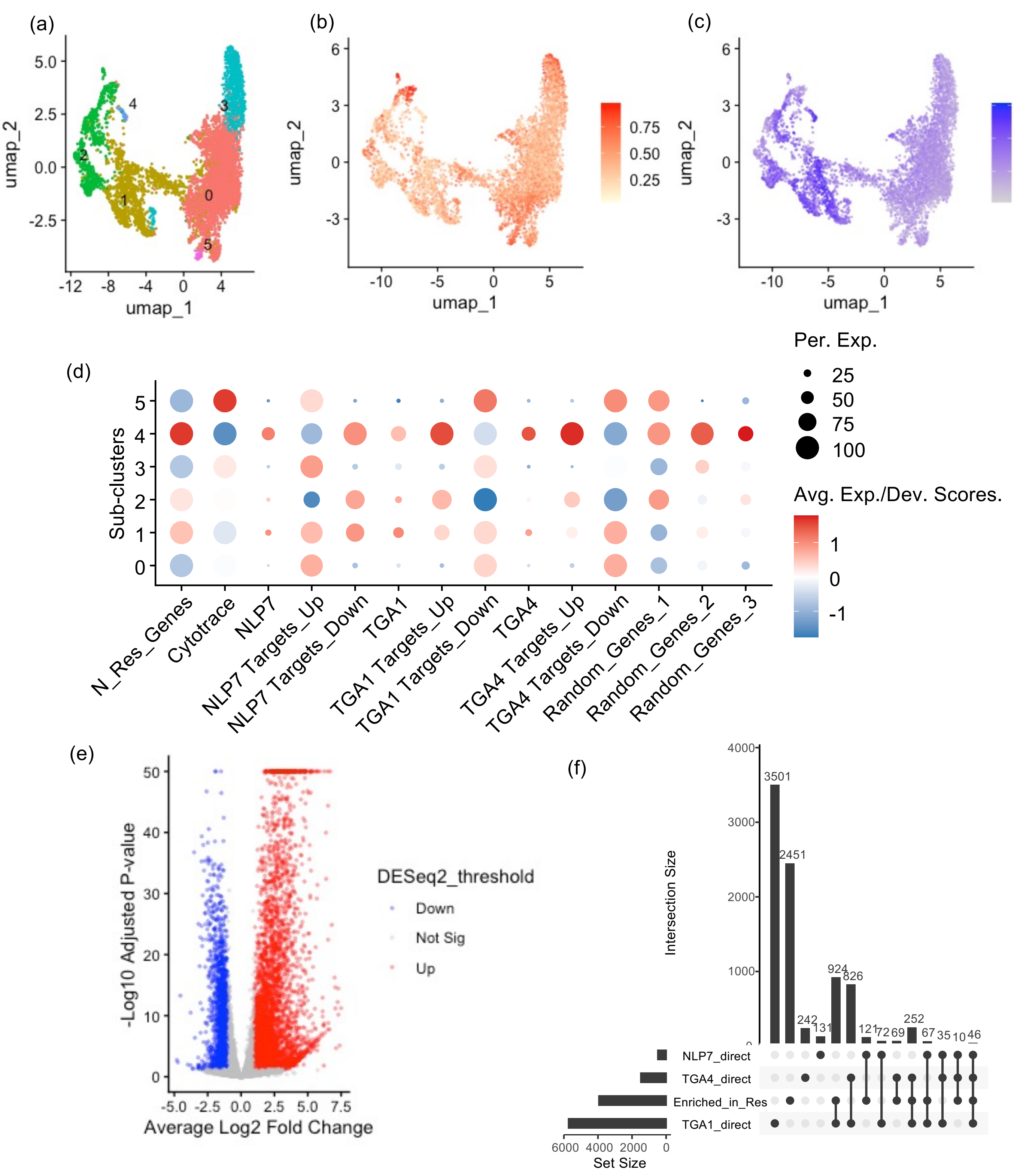


**Supplementary Figure S38. Sub-population analysis of Arabidopsis procambium in the *anac032* mutant. (a)** UMAP visualization of procambium sub-clusters. **(b)** CytoTRACE scores. **(c)** UMAP visualization of the average expression of the N-responsive genes (Varala et al. 2018). **(d)** Scaled CytoTRACE scores and average expression of corresponding genes in subclusters. N_Res_Genes, N-responsive genes (Varala et al. 2018). NLP7 Targets_Up/Down, direct targets of AtNLP7 (Alvarez et al. 2020). TGA1/TGA4 Targets_Up/Down, direct targets of AtTGA1/TGA4 (Brooks et al. 2019). Random_Genes_1/2/3, three independent sets of 2652 randomly sampled genes. **(e)** Volcano plots of the differently expressed gene between responsive (sub-cluster 2) and non-responsive (sub-clusters 0 and 1) sub-clusters. Red dots indicate genes that are significantly up-regulated in the responsive non-responsive, mature (adjusted p_value < 0.05, log2 fold changes > 1) using DESeq2 on pseudobulked sub-populations. Blue dots for down-regulated genes. **(f)** Overlaps across NLP7/TGA1/TGA4 direct targets and genes that are up-regulated in the responsive sub-population.

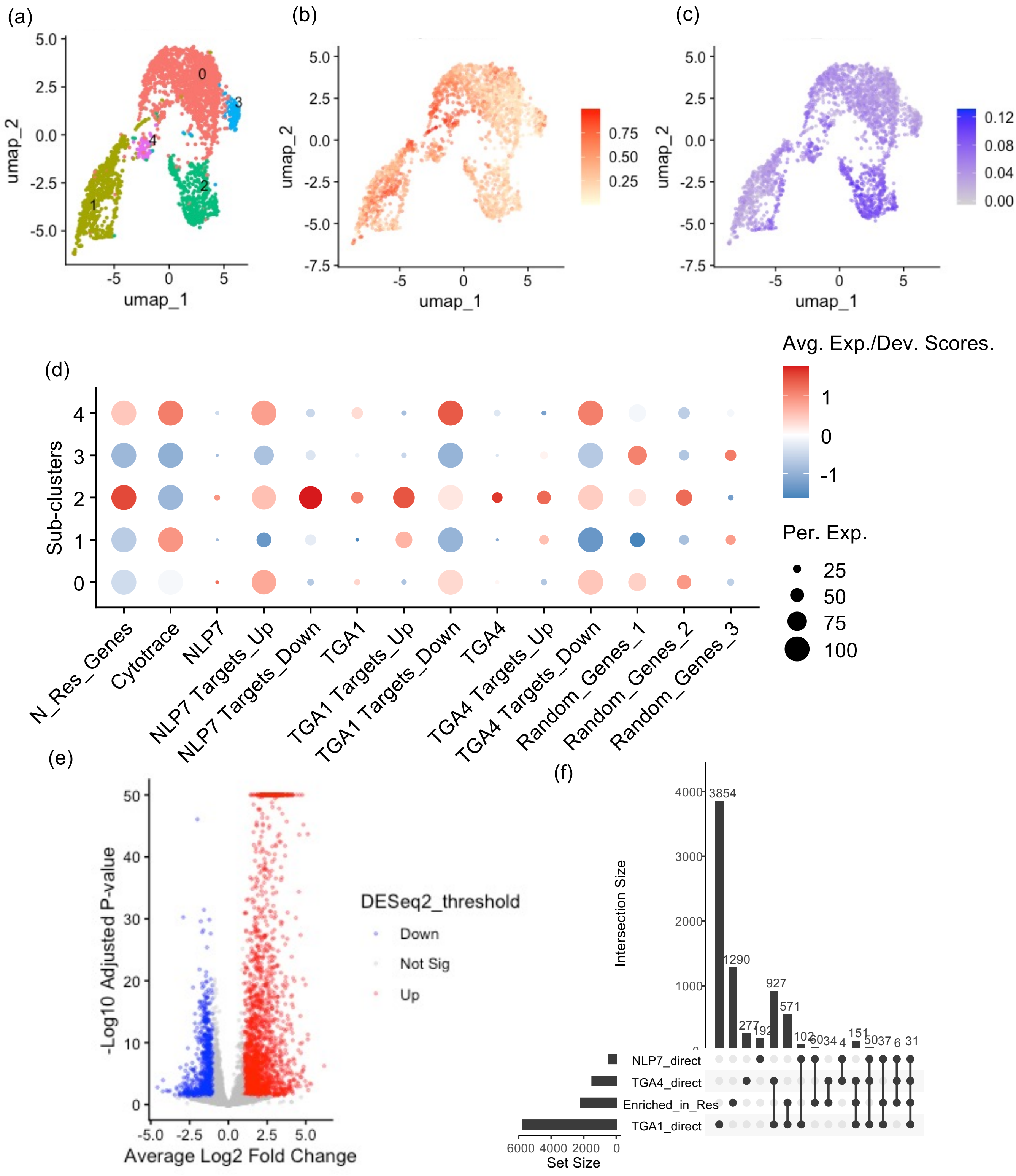


**Supplementary Figure S39. Sub-population analysis of Arabidopsis phloem in the *anac032* mutant.** (a) UMAP visualization of phloem sub-clusters. **(b)** CytoTRACE scores. **(c)** UMAP visualization of the average expression of the N-responsive genes (Varala et al. 2018). **(d)** Scaled CytoTRACE scores and average expression of corresponding genes in subclusters. N_Res_Genes, N-responsive genes (Varala et al. 2018). NLP7 Targets_Up/Down, direct targets of AtNLP7 (Alvarez et al. 2020). TGA1/TGA4 Targets_Up/Down, direct targets of AtTGA1/TGA4 (Brooks et al. 2019). Random_Genes_1/2/3, three independent sets of 2652 randomly sampled genes. **(e)** Volcano plots of the differently expressed gene between responsive (sub-cluster 2) and non-responsive (sub-clusters 0 and 1) sub-clusters. Red dots indicate genes that are significantly up-regulated in the responsive non-responsive, mature (adjusted p_value < 0.05, log2 fold changes > 1) using DESeq2 on pseudobulked sub-populations. Blue dots for down-regulated genes. **(f)** Overlaps across NLP7/TGA1/TGA4 direct targets and genes that are up-regulated in the responsive sub-population.

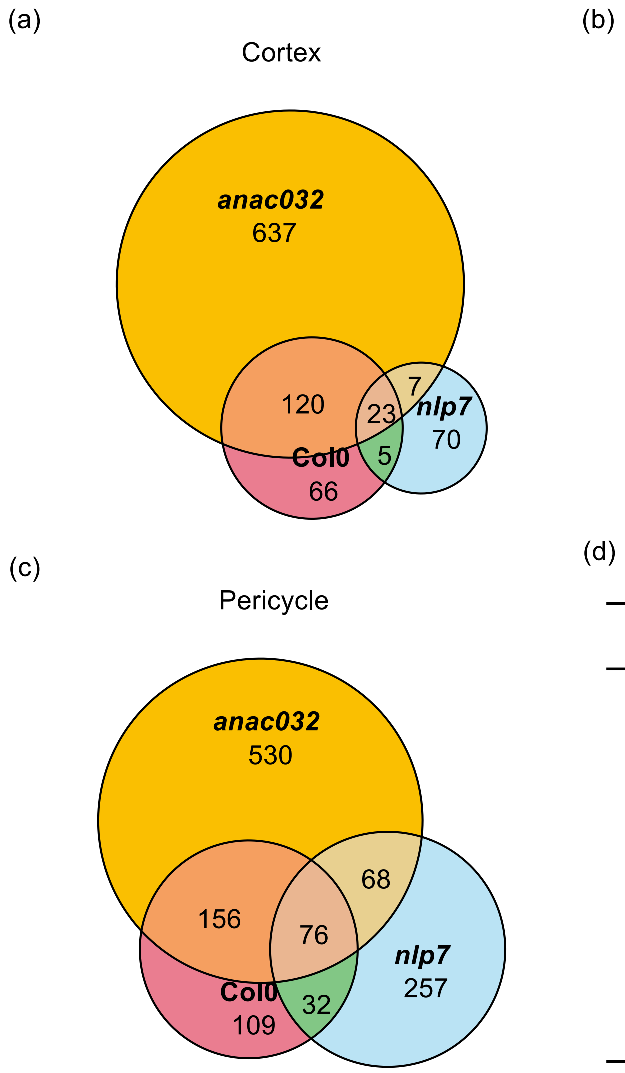


(d)

| Cell type | Comparison | p_value |
|---|---|---|
| Cortex | Col0 vs *nlp7* | $< 2.2*10^{-16}$ |
| Cortex | Col0 vs *anac032* | $< 2.2*10^{-16}$ |
| Endodermis | Col0 vs *nlp7* | $< 2.2*10^{-16}$ |
| Endodermis | Col0 vs *anac032* | $< 2.2*10^{-16}$ |
| Pericycle | Col0 vs *nlp7* | $< 2.2*10^{-16}$ |
| Pericycle | Col0 vs *anac032* | $< 2.2*10^{-16}$ |

**Supplementary Figure S40. Common DEGs were found in the cortex, endodermis and pericycle of Arabidopsis wild type and mutants. (a-c)** Euler plots showing genes up-regulated in the high $KNO_3$ condition in cortex (**a**), endodermis (**b**) and pericycle (**c**) of Col-0 wild-type and mutants (*anac032* and *nlp7*). **(d)** Overrepresentation of overlapping DEGs genes in each cell type between Col-0 and mutants using Fisher's exact test.

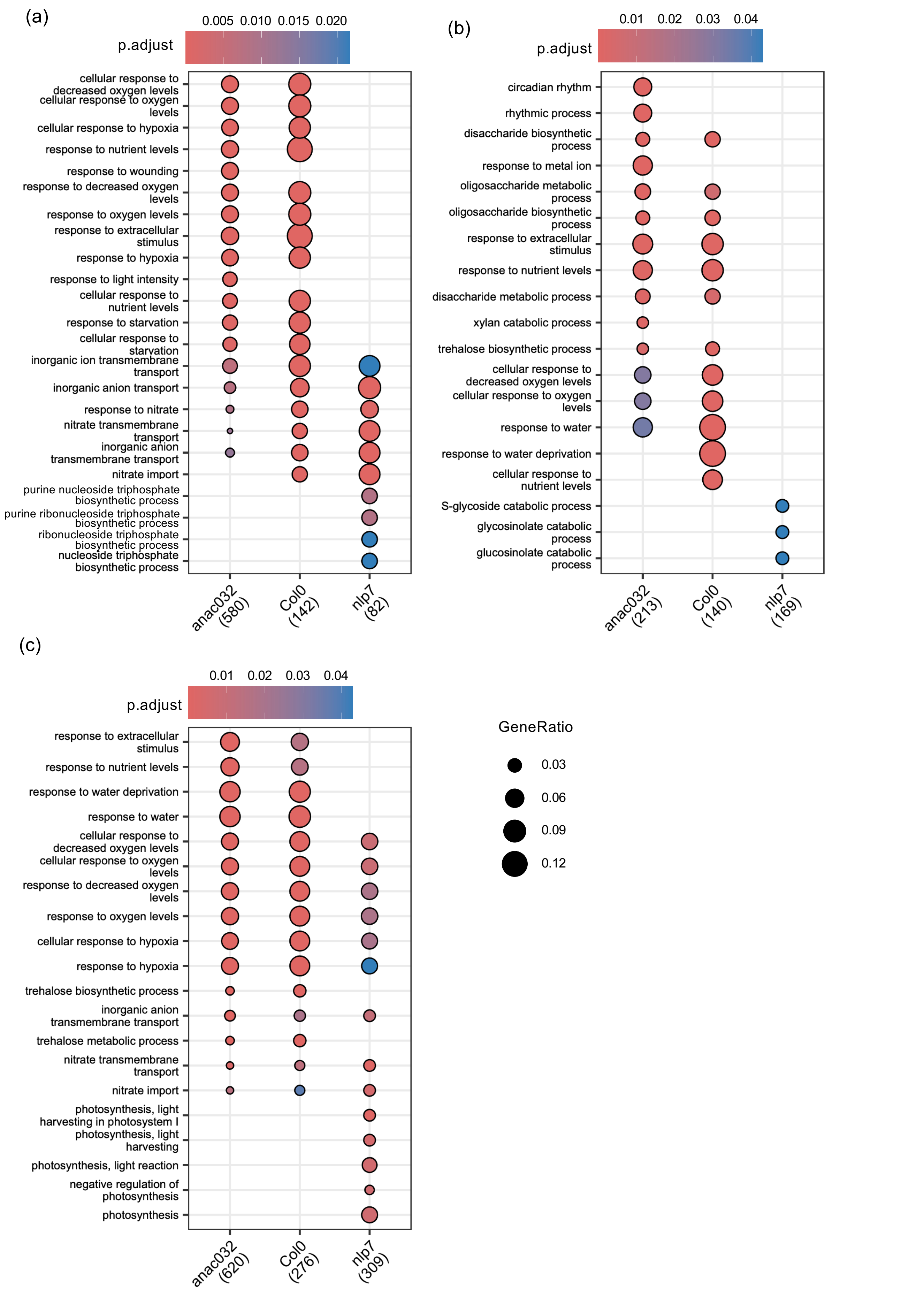


**Supplementary Figure S41. GO terms enriched in the DEGs in the cortex, endodermis and pericycle of Arabidopsis wild type and mutants. (a)** Cortex. **(b)** Endodermis. **(c)** Pericycle. Up to 10 top GO terms in each genotype were shown.

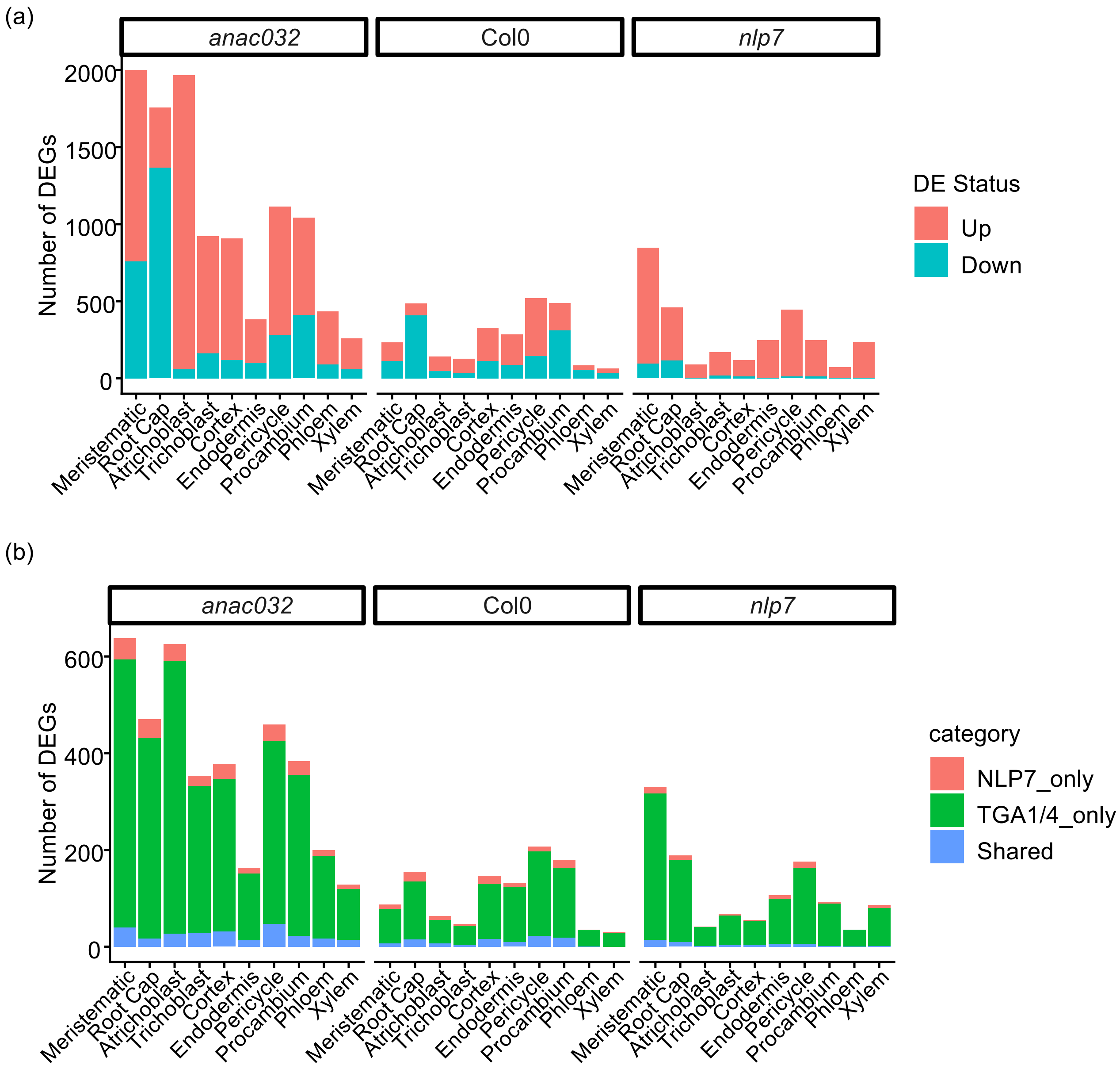


**Supplementary Figure S42. Summary of DEGs in Arabidopsis cell types. (a)** Number of DEGs in each genotype. DE status, 10 mM vs 1 mM $KNO_3$. **(b)** Number of DEGs that are targets of NLP7/TGA1/TGA4 in each genotype. NLP7_only, direct targets of only NLP7. TGA1/4_only, direct targets of only TGA1 or TGA4. Shared, direct targets of all of NLP7, TGA1 and TGA4.

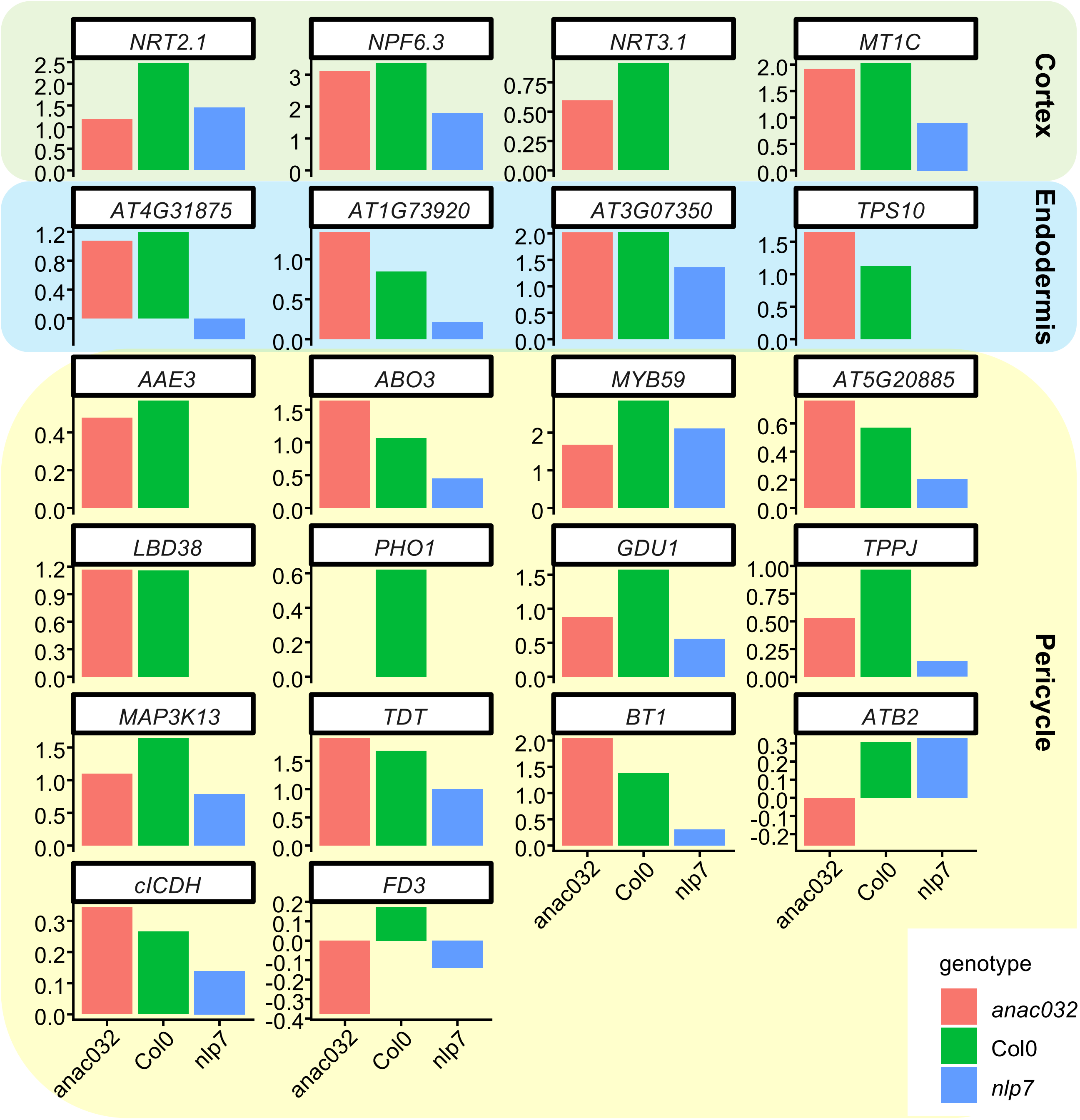


**Supplementary Figure S43. Nitrate responses of AtNLP7 targets across Arabidopsis genotypes.** Log 2 fold changes (10 vs 1mM $KNO_3$) of AtNLP7 direct targets in Fig4c in cell types where each gene was enriched and N-responsive.

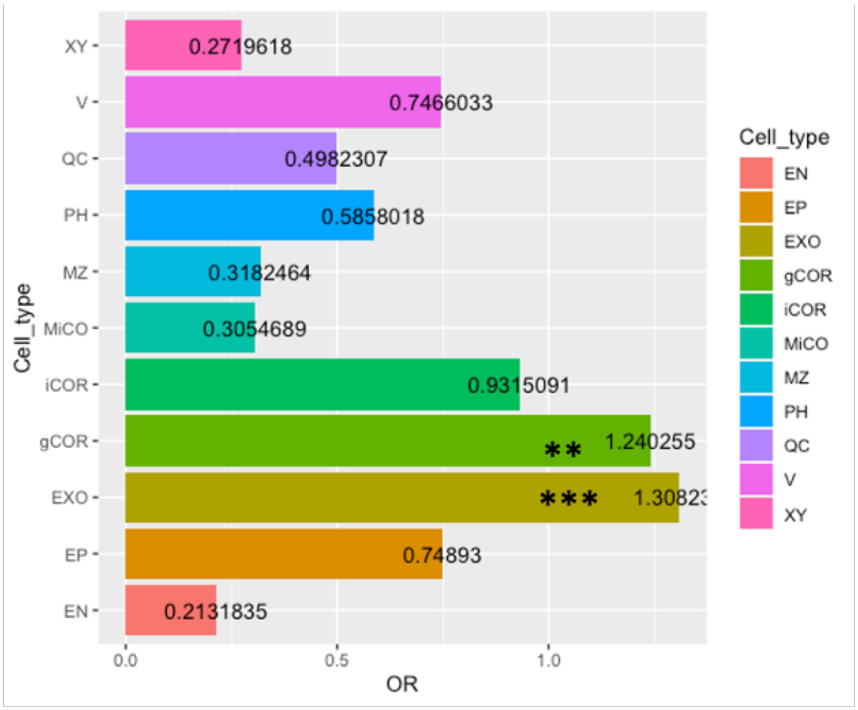


**Supplementary Figure S44. Over-representation of cell type-enriched ribosome-associated transcripts (Kajala et al., 2021) within N concentration-dependent differentially expressed genes in the tomato whole root (Bian et al., 2025).** Odd's ratio (OR) are presented for each cell type dataset. XY, xylem. V, vasculature. QC, quiescent centre. PH, phloem. MZ, meristematic zone. MiCO, meristematic inner cortex. iCOR, inner cortex. gCOR, general cortex. EXO, exodermis. EP, epidermis. EN, endodermis; *** = p<.001; ** = p<.01.

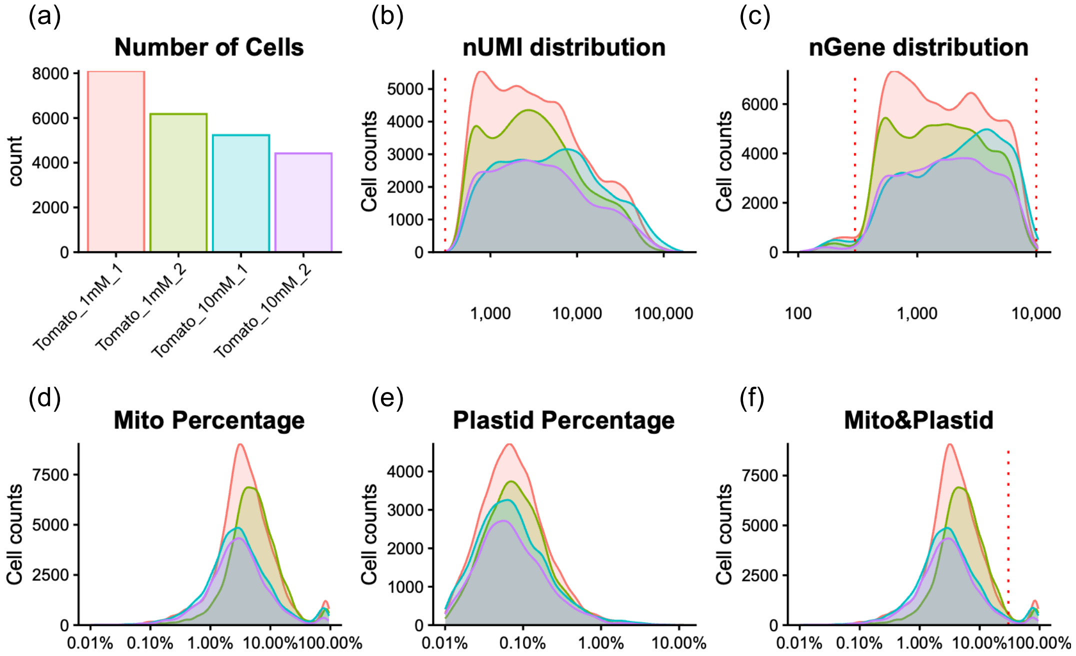


**Supplementary Figure S45. Cell and read distributions of tomato scRNAseq samples from different $KNO_3$ conditions and filtering cutoffs. (a)** Number of cells in each sample before filtering. (**b-f**) Distribution of the number of nUMI (unique molecular identifier) (b), number of genes (c), percentage of mitochondrial transcripts (d), percentage of chloroplast transcripts (e), and percentage of the sum of mitochondrial and chloroplast transcripts (f) in each sample. Filtering thresholds are indicated by red dash lines (UMIs > 300, detected genes between 300 to 10000, and organellar (mitochondrial and chloroplast) transcripts <30%).

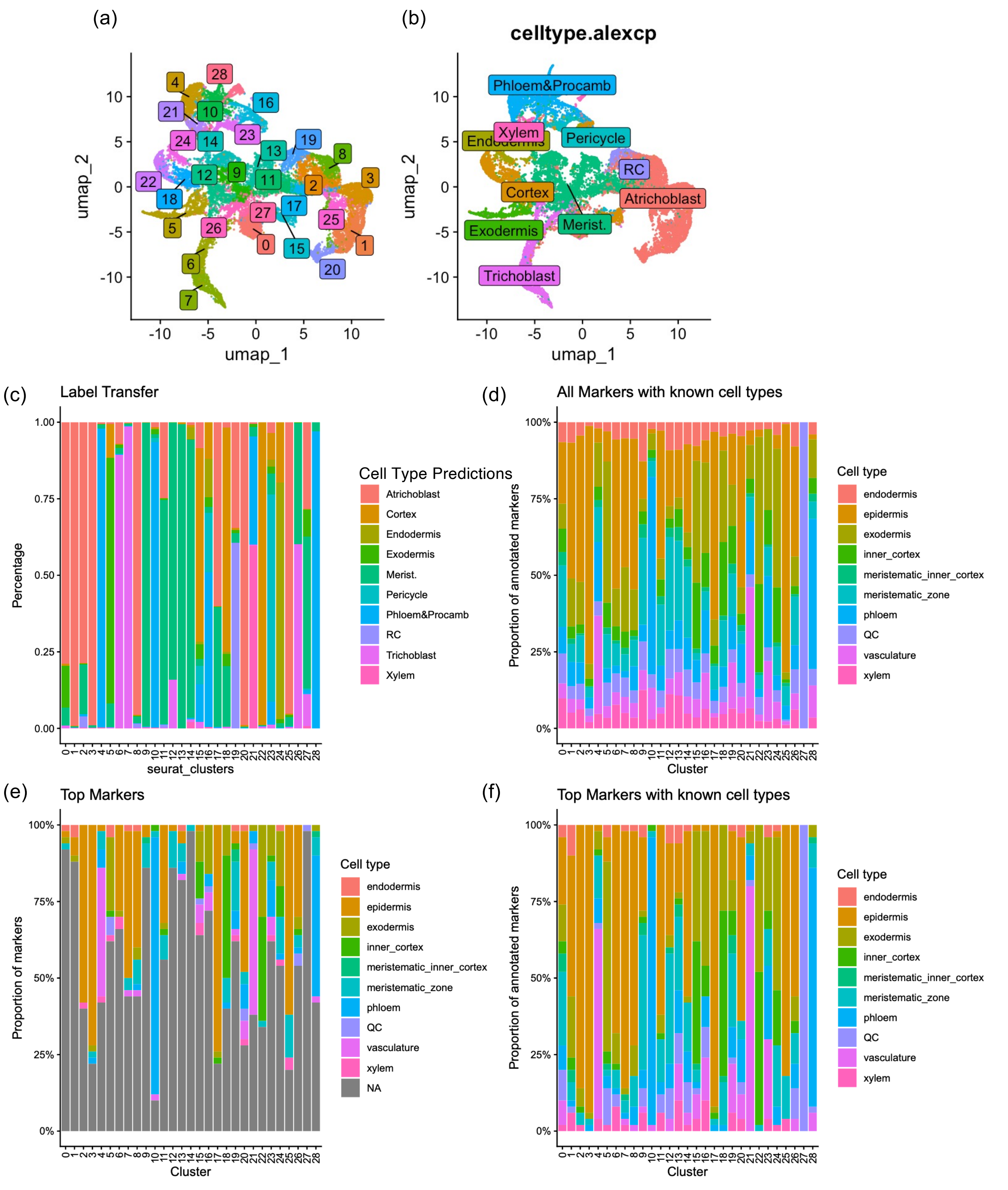


**Supplementary Figure S46. Cell type prediction of tomato scRNAseq dataset from different $KNO_3$ conditions. (a)** UMAP visualization of Seurat clusters. **(b)** Cell type predictions by label transfer method using the annotated tomato root atlas from Cantó-Pastor et al. (2024) as the reference. **(c)** Percentage of predicted cell type labels in each cluster using the label transfer method. **(d)** Percentage of cell types where markers with known expression patterns in each cluster are expressed. **(e)** Percentage of cell types where top 50 markers in each cluster are expressed. **(f)** Percentage of cell types where top 50 markers with known expression patterns in each cluster are expressed. Cell type expression patterns of marker genes are from the cell-type-enriched transcriptomes generated by FACS sorting (Kajala et al. 2021).

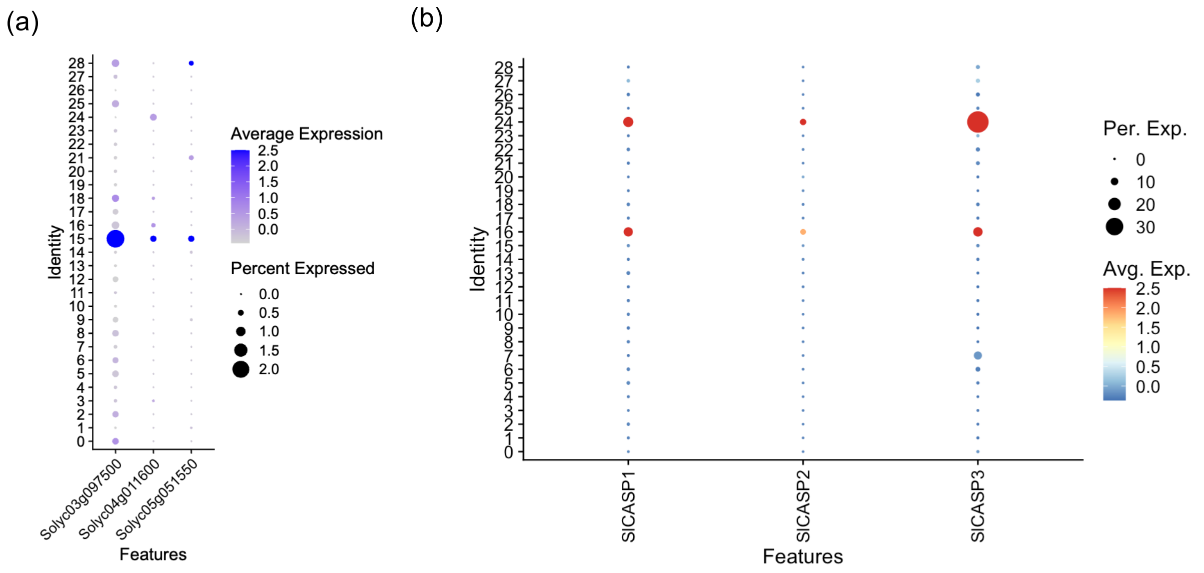


**Supplementary Figure S47. Refinement of annotation of tomato scRNAseq clusters from different $KNO_3$ conditions. (a)** Three genes known to be enriched in the mature exodermal cells were mostly expressed in cluster 15. **(b)** *SlCASP1-3,* which are known to be expressed in endodermal cells, were expressed in cluster 16 and a clearly identified endodermis cluster 24.

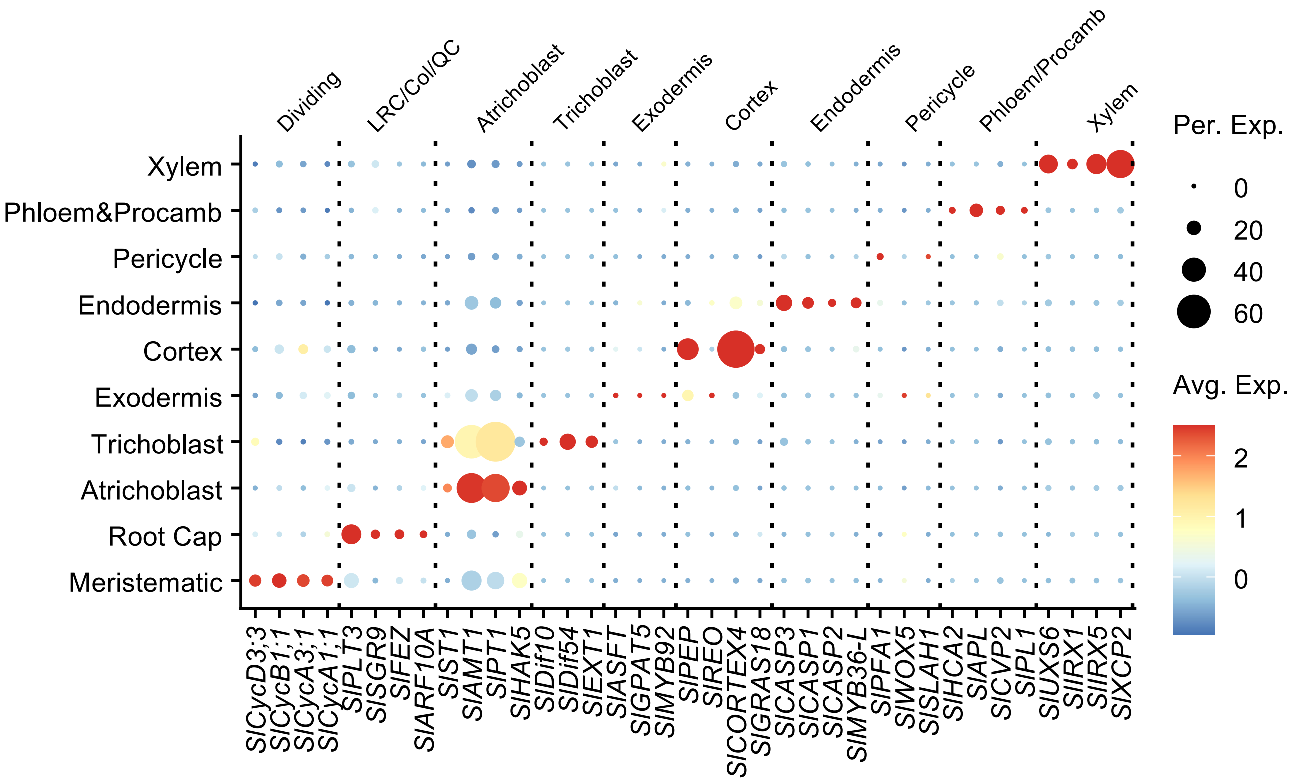


**Supplementary Figure S48. Validation of cell type annotation of tomato scRNAseq from different $KNO_3$ conditions using marker genes with known expression patterns.** LRC, lateral root cap. Col, columella. QC, quiescent center. Procamb, procambium. Per. Exp., percentage of cells expressing the gene. Avg. Exp., scaled average expression level.

(a) (b)

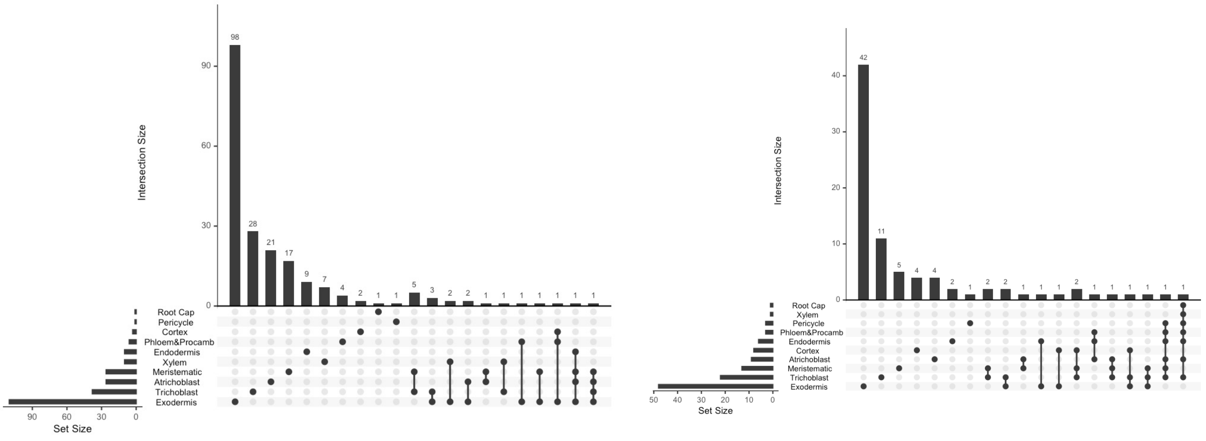


**Supplementary Figure S49. Overlaps of DEGs between different $KNO_3$ conditions in tomato root cell types**. **(a)** Genes up-regulated at high $KNO_3$ conditions. **(d)** Genes down-regulated at high $KNO_3$ conditions.

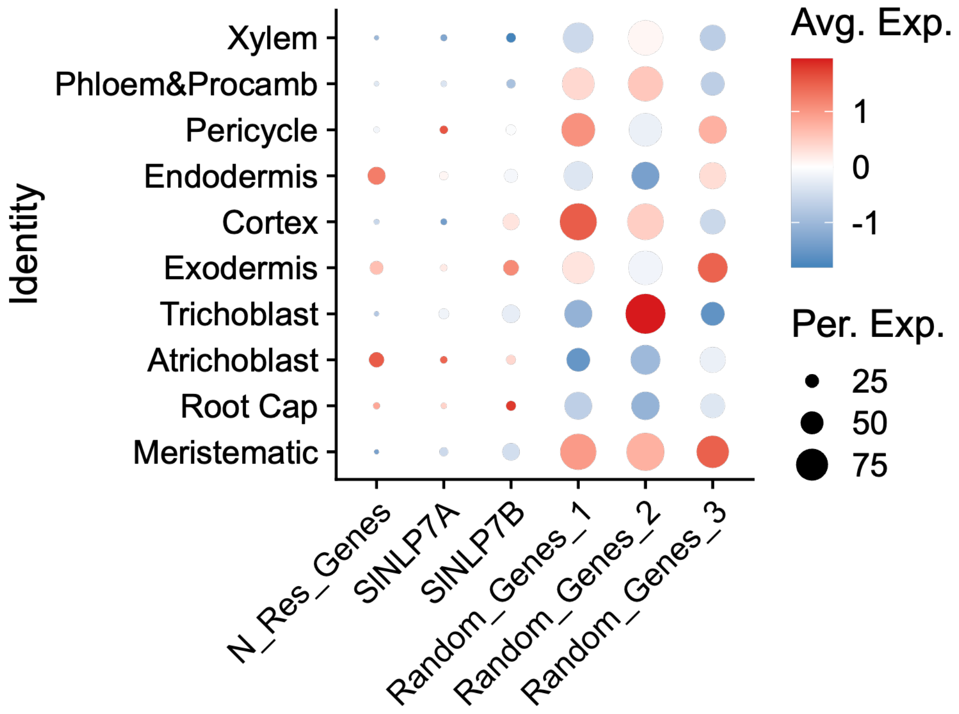


**Supplementary Figure S50. Expression of N-responsive genes in tomato root cell types.** N_Res_Genes, 1199 genes that were significantly differentially expressed in the bulk RNAseq of roots from tomato plants grown under the same high and low $KNO_3$ conditions as scRNAseq was performed. Expression of *SlNLP7A* and *SlNLP7B* were also included. Random_Genes_1/2/3, three independent sets of 636 randomly sampled genes used as controls in the later sub-clustering analysis.

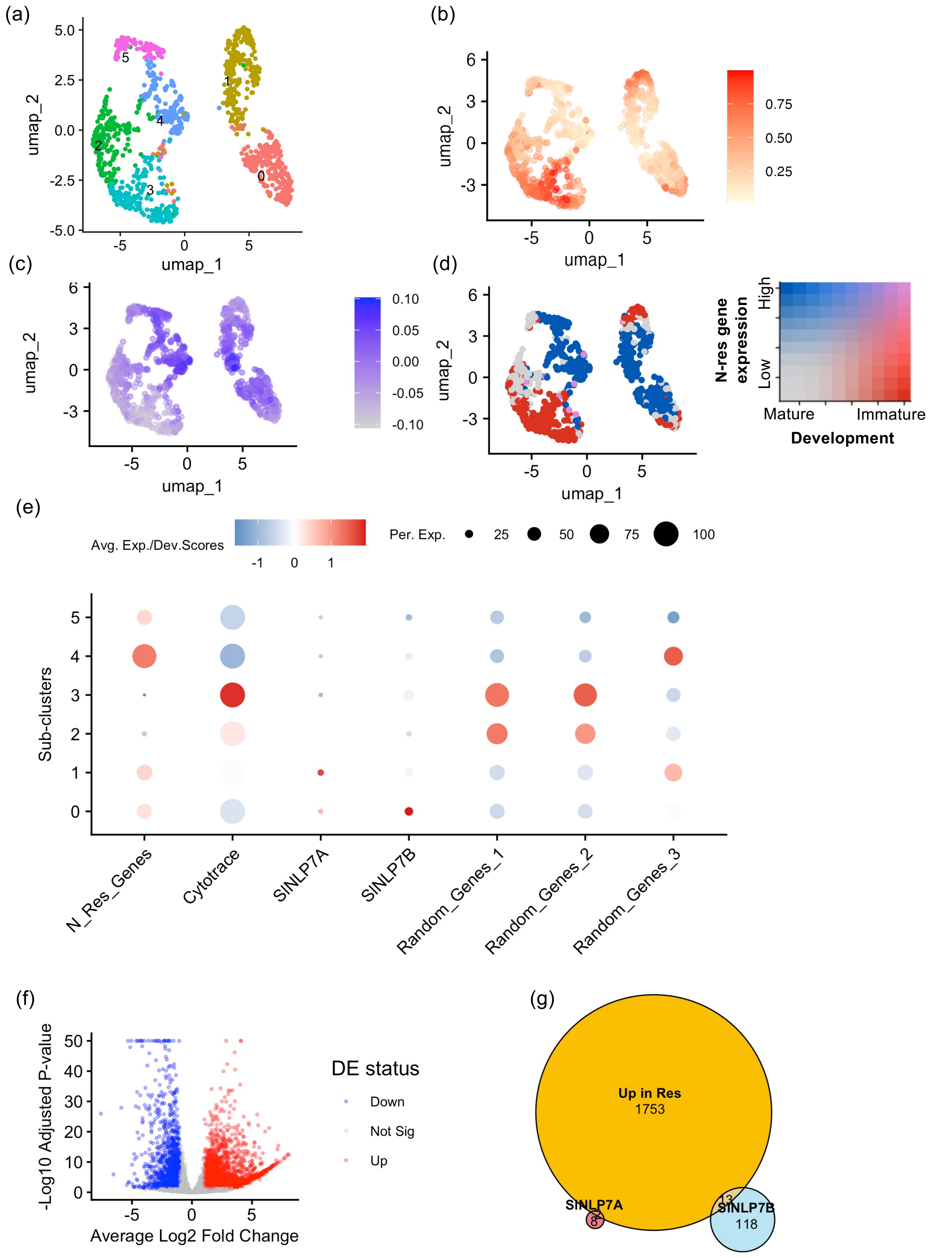


**Supplementary Figure S51. Sub-population analysis of tomato endodermis.** (a) UMAP visualisation of endodermis sub-clusters. **(b)** CytoTRACE scores. **(c)** Average expression of the N-responsive genes (from bulk RNAseq in the same growth conditions). **(d)** Correlation of N-responsive genes with maturity in sub-clusters. **(e)** Scaled CytoTRACE scores and average expression of corresponding genes in subclusters. N_Res_Genes, N-responsive genes from bulk RNAseq. Random_Genes_1/2/3, three independent sets of 1199 randomly sampled genes. **(f)** Volcano plots of the differently expressed gene between responsive (sub-cluster 1) and non-responsive, mature (sub-cluster 3, 4, 5, 6) sub-clusters. Red dots indicate genes that are significantly up-regulated in the responsive sub-population (adjusted p_value < 0.05, log2 fold changes > 1) using DESeq2 on pseudobulked sub-populations. Blue dots for down-regulated genes. **(g)** Overlaps between genes that are up-regulated in the responsive sub-population and cell-type-specific targets that are activated by SlNLP7A or SlNLP7B in this cell type.

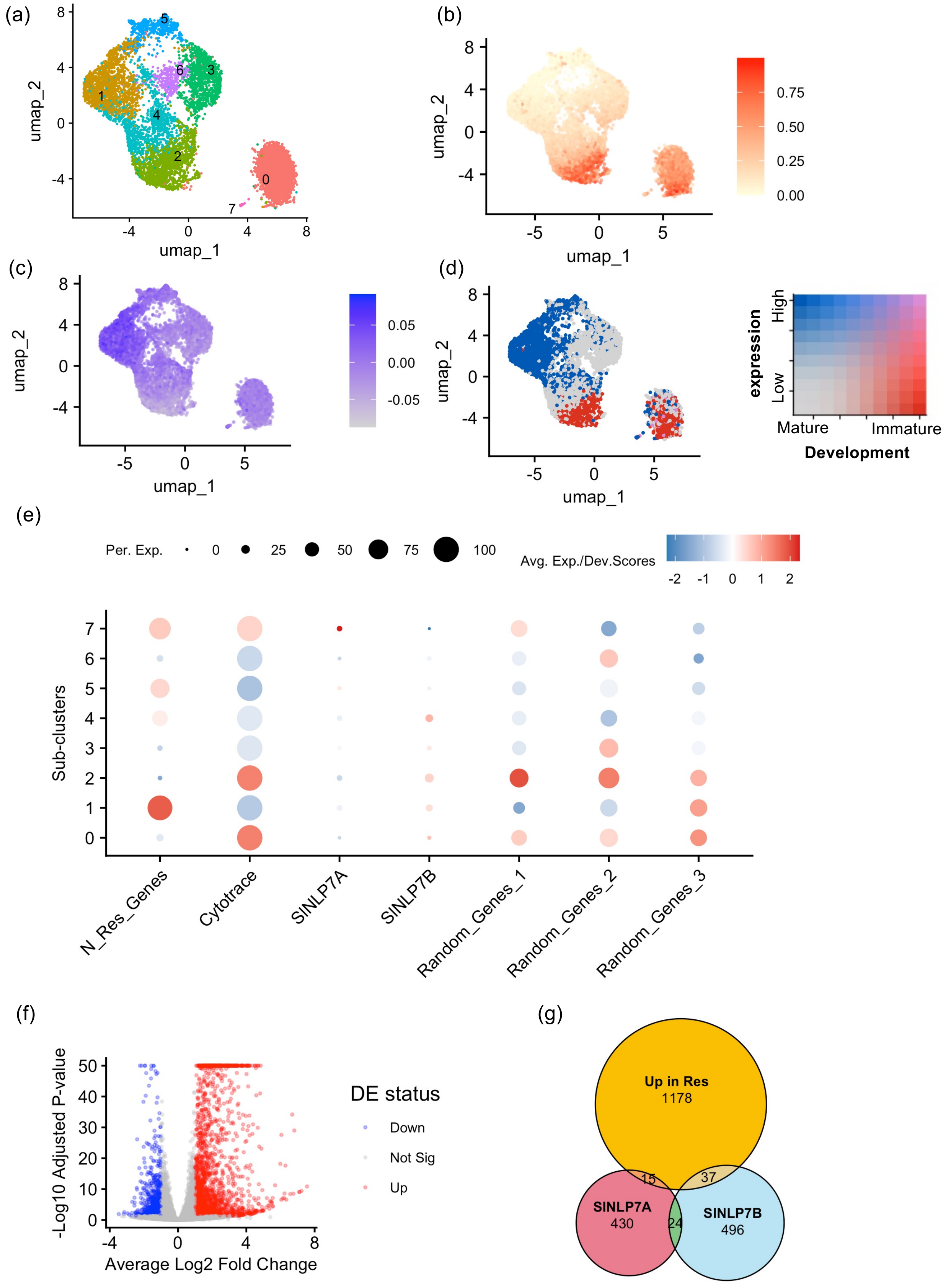


**Supplementary Figure S52. Sub-population analysis of tomato atrichoblast. (a)** UMAP visualisation of atrichoblast sub-clusters. **(b)** CytoTRACE scores. **(c)** Average expression of the N-responsive genes (from bulk RNAseq in the same growth conditions). **(d)** Correlation of N-responsive genes with maturity in sub-clusters. **(e)** Scaled CytoTRACE scores and average expression of corresponding genes in subclusters. N_Res_Genes, N-responsive genes from bulk RNAseq. Random_Genes_1/2/3, three independent sets of 1199 randomly sampled genes. **(f)** Volcano plots of the differently expressed gene between responsive (sub-clusters 1 and 5) and non-responsive, mature (sub-cluster 3, 4, 6) sub-clusters. Red dots indicate genes that are significantly up-regulated in the responsive sub-population (adjusted p_value < 0.05, log2 fold changes > 1) using DESeq2 on pseudobulked sub-populations. Blue dots for down-regulated genes. **(g)** Overlaps between genes that are up-regulated in the responsive sub-population and cell-type-specific targets that are activated by SlNLP7A or SlNLP7B in this cell type.

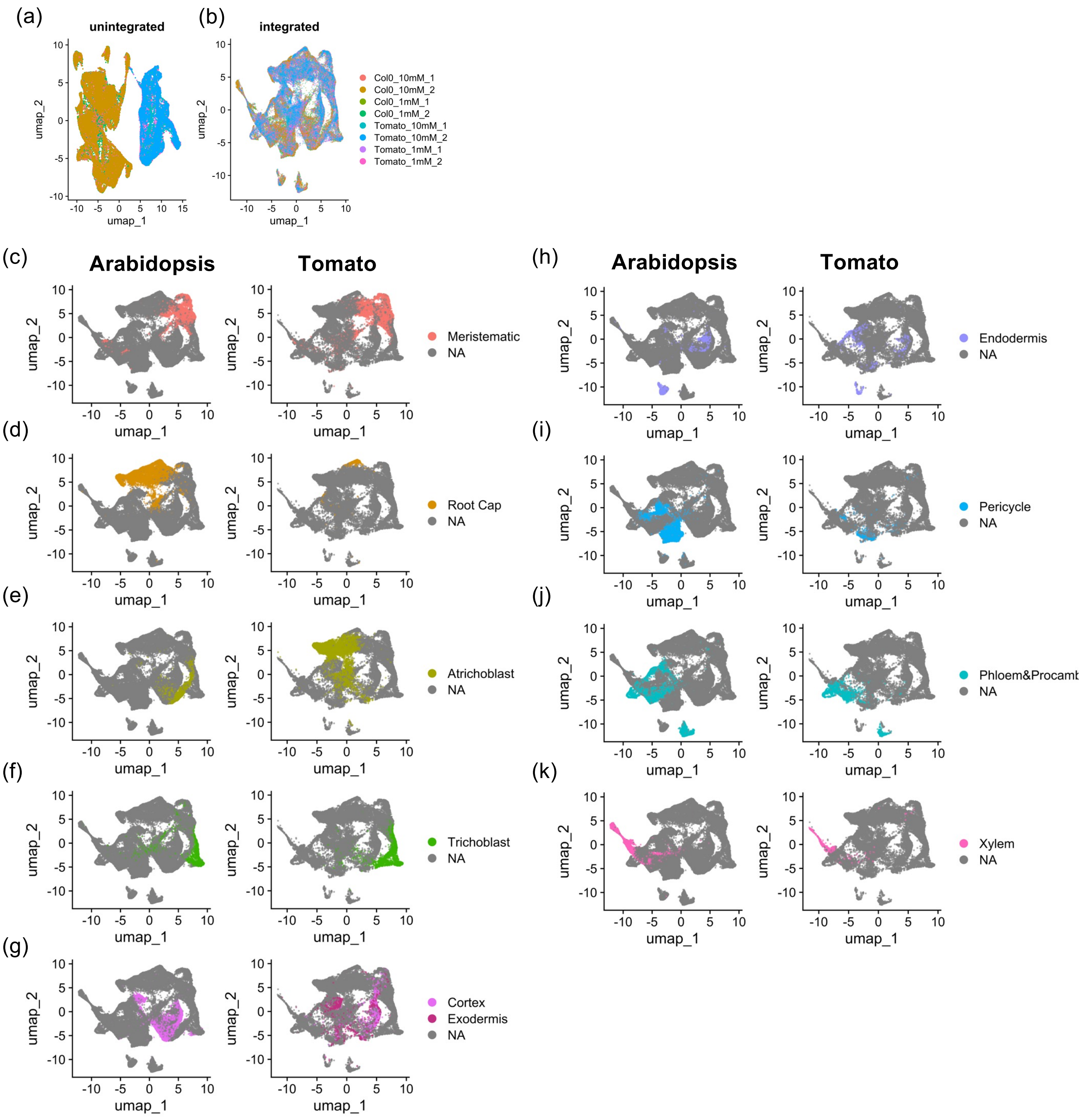


**Supplementary Figure S53. Integration of Arabidopsis (wild-type) and tomato scRNAseq datasets by combining transcripts of all genes in a homology group based on Kajala et al. (2021) expressologs**. (**a**,**b**) UMAPs of Arabidopsis and tomato cells before (**a**) and after (**b**) integration. (**c-k**) UMAPs highlighting the same groups of cell types in Arabidopsis (left) and tomato (right).

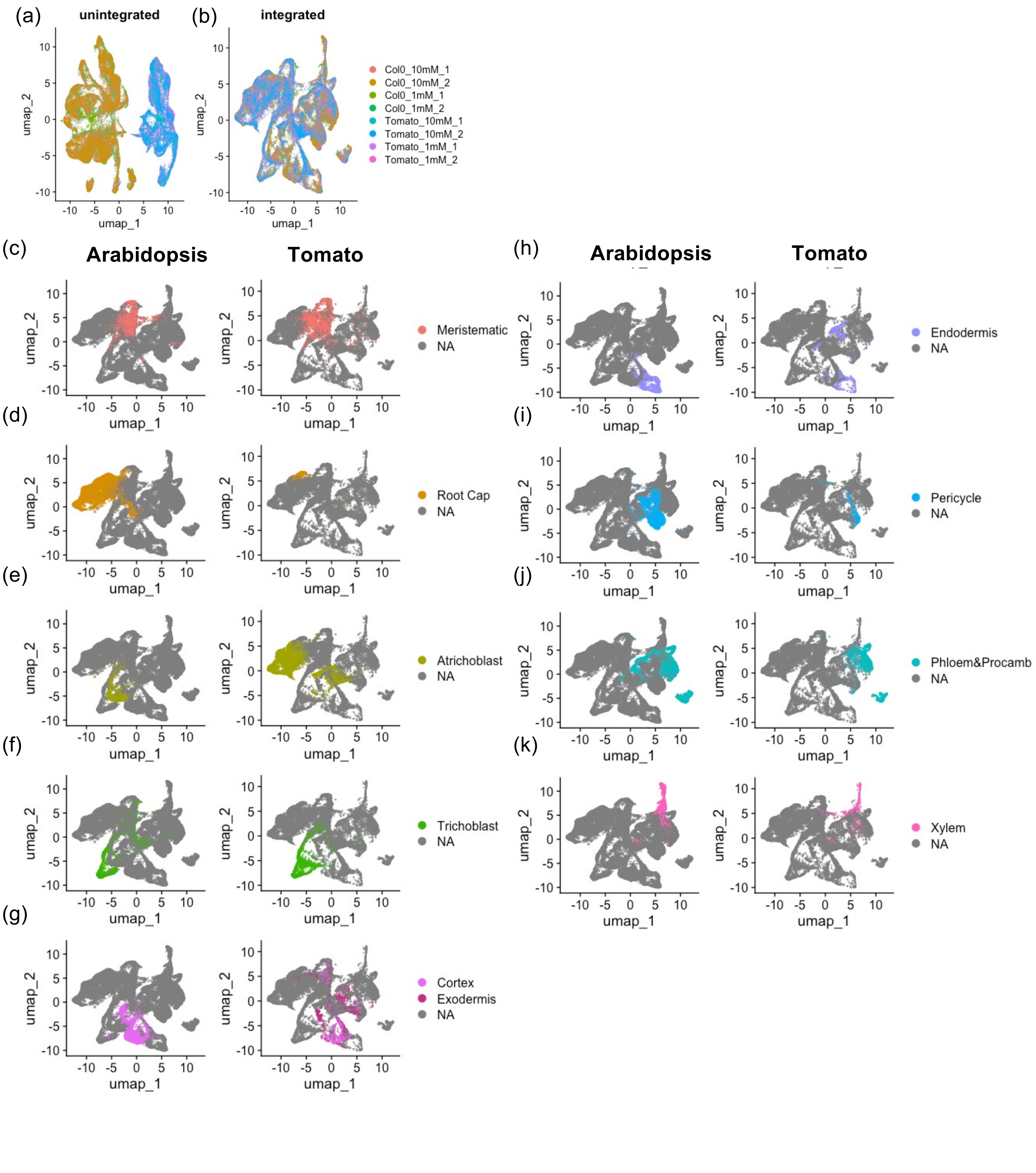


**Supplementary Figure S54. Integration of Arabidopsis (wild-type) and tomato scRNAseq datasets by combining transcripts of all genes in a homology group based on EnsemblPlant homology database**. (**a,b**) UMAPs of Arabidopsis and tomato cells before (**a**) and after (**b**) integration. (**c-k**) UMAPs highlighting the same groups of cell types in Arabidopsis (left) and tomato (right).

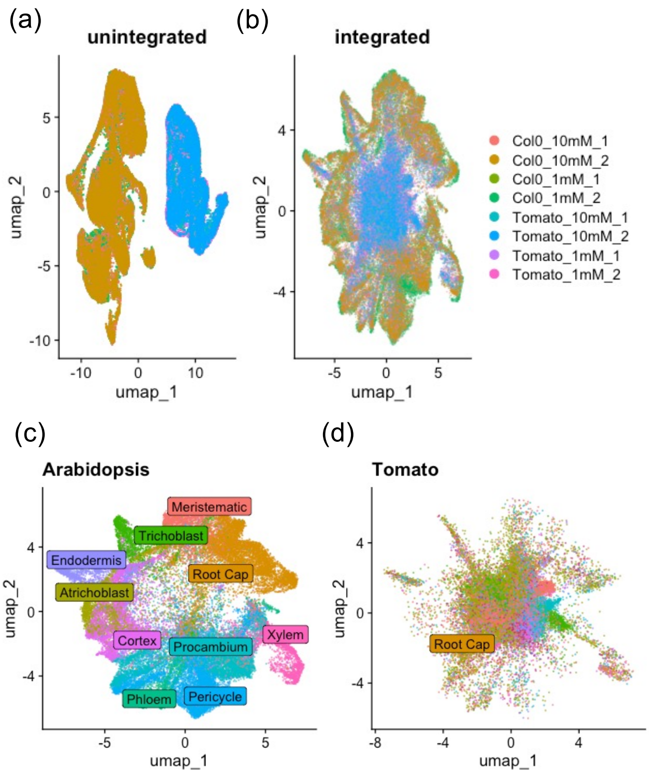


**Supplementary Figure S55. Integration of Arabidopsis (wild-type) and tomato scRNAseq datasets by taking averages of all genes in a homology group based on Kajala et al. (2021) expressologs.** (**a,b**) UMAPs of Arabidopsis and tomato cells before (**a**) and after (**b**) integration. (**c-d**) UMAP representation of Arabidopsis (**c**) and tomato (**d**) cell type annotations.

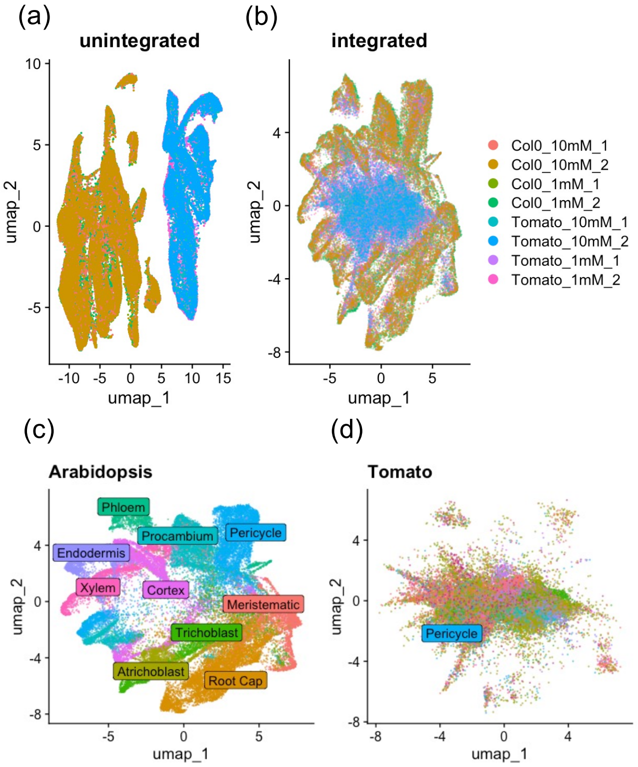


**Supplementary Figure S56. Integration of Arabidopsis (wild-type) and tomato scRNAseq datasets by taking averages of all genes in a homology group based on EnsemblPlant homology database.** (**a**,**b**) UMAPs of Arabidopsis and tomato cells before (**a**) and after (**b**) integration. (**c-d**) UMAP representation of Arabidopsis (**c**) and tomato (**d**) cell type annotations.

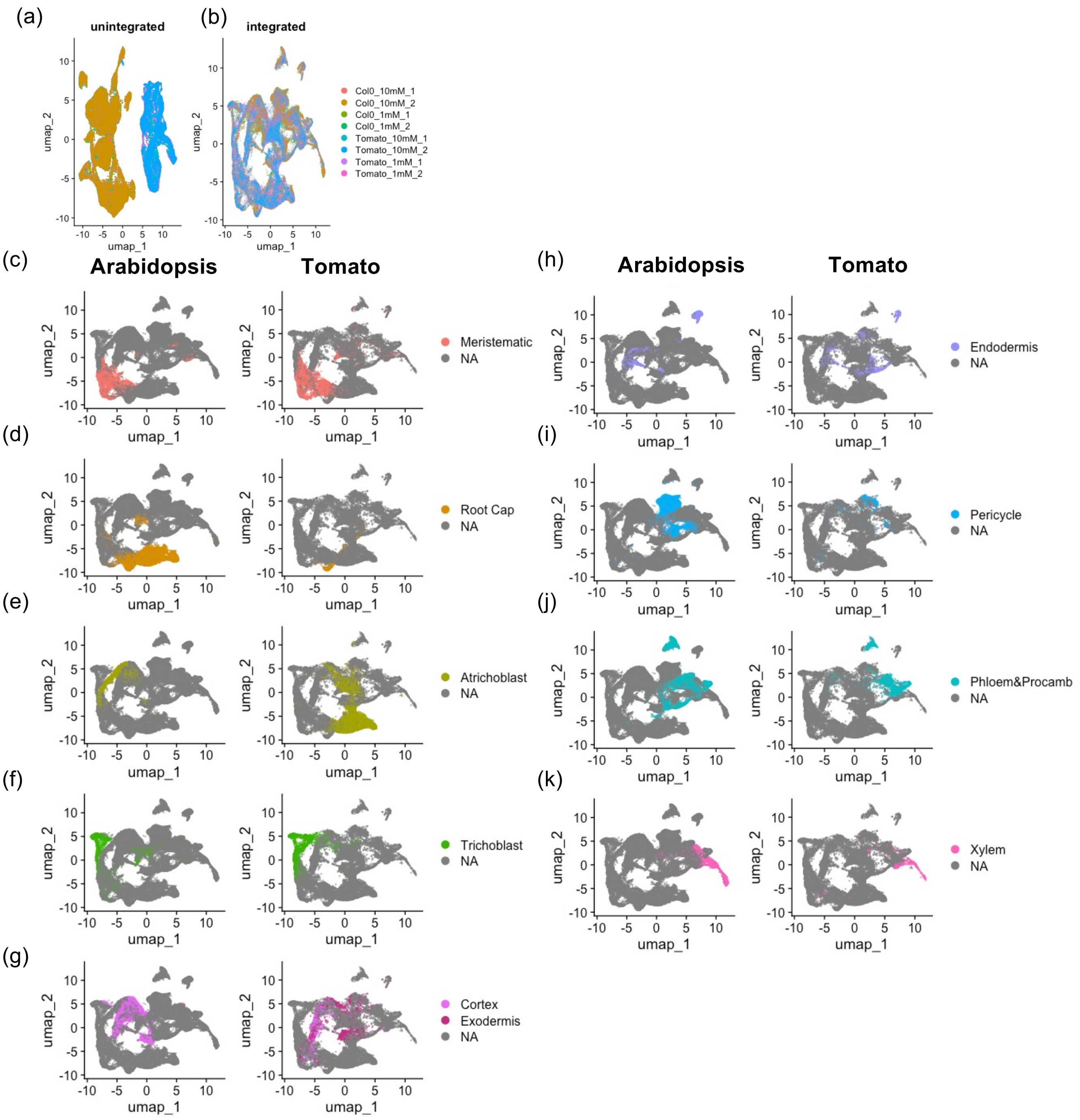


**Supplementary Figure S57. Integration of Arabidopsis (wild-type) and tomato scRNAseq datasets by using the most abundant gene in a homology group based on Kajala et al. (2021) expressologs.** (**a**,**b**) UMAPs of Arabidopsis and tomato cells before (**a**) and after (**b**) integration. (**c-k**) UMAPs highlighting the same groups of cell types in Arabidopsis (left) and tomato (right).

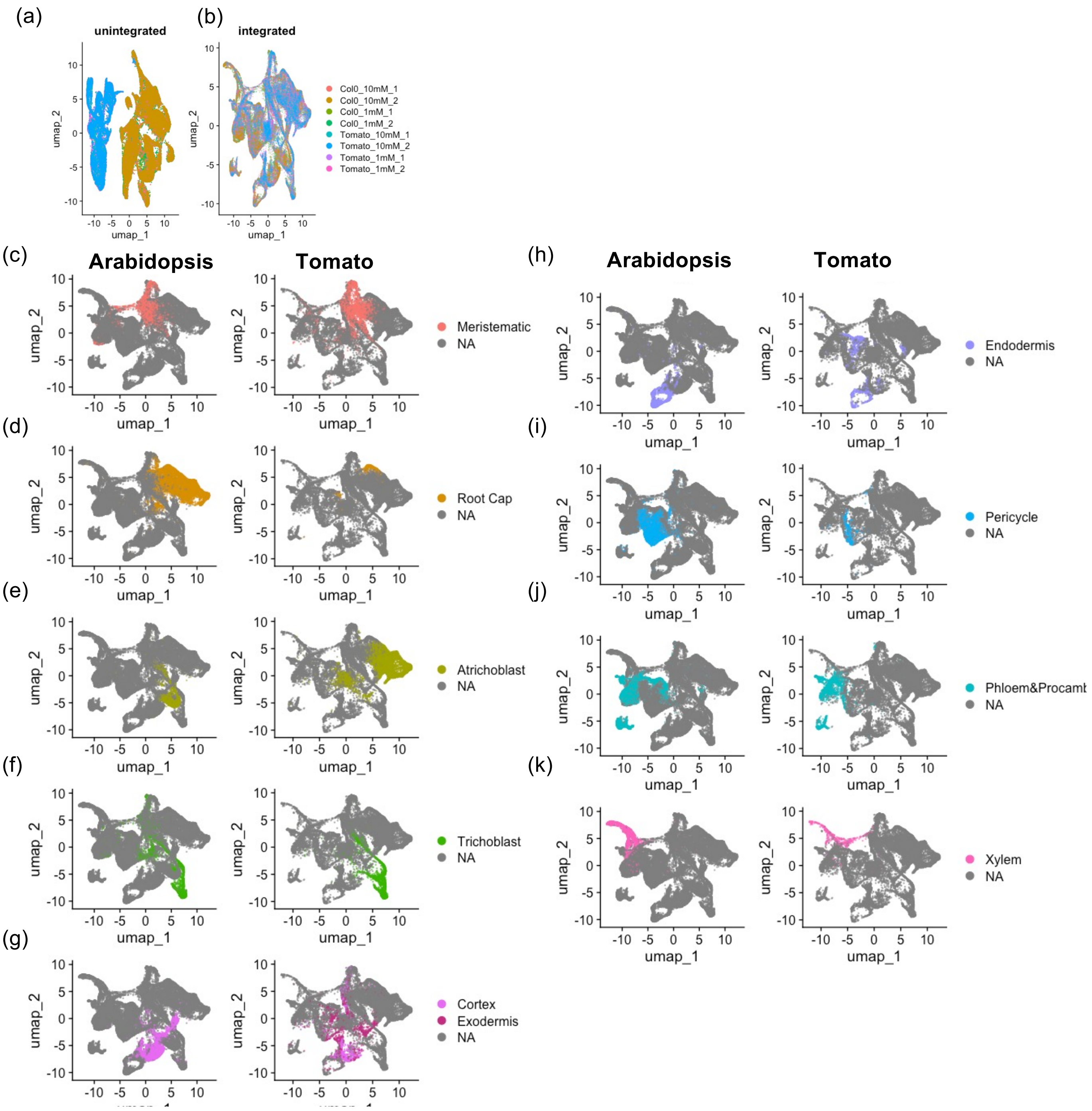


**Supplementary Figure S58. Integration of Arabidopsis (wild-type) and tomato scRNAseq datasets by using the most abundant gene in a homology group based on EnsemblPlant homology database.** (**a,b**) UMAPs of Arabidopsis and tomato cells before (**a**) and after (**b**) integration. (**c-k**) UMAPs highlighting the same groups of cell types in Arabidopsis (left) and tomato (right).

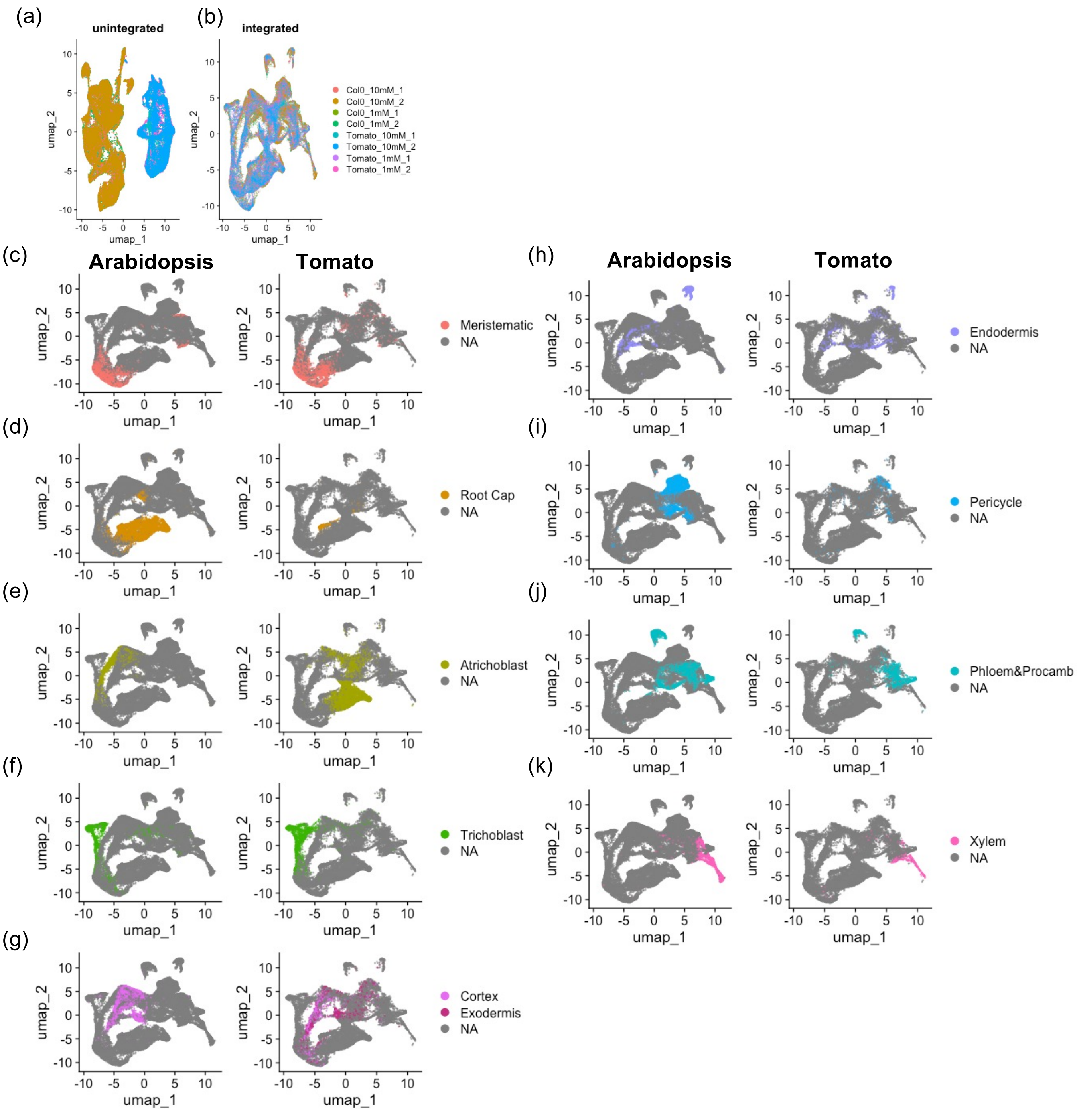


**Supplementary Figure S59. Integration of Arabidopsis (wild-type) and tomato scRNAseq datasets by one-to-one orthologs based on Kajala et al. (2021) expressologs.** (**a,b**) UMAPs of Arabidopsis and tomato cells before (**a**) and after (**b**) integration. (**c-k**) UMAPs highlighting the same groups of cell types in Arabidopsis (left) and tomato (right).

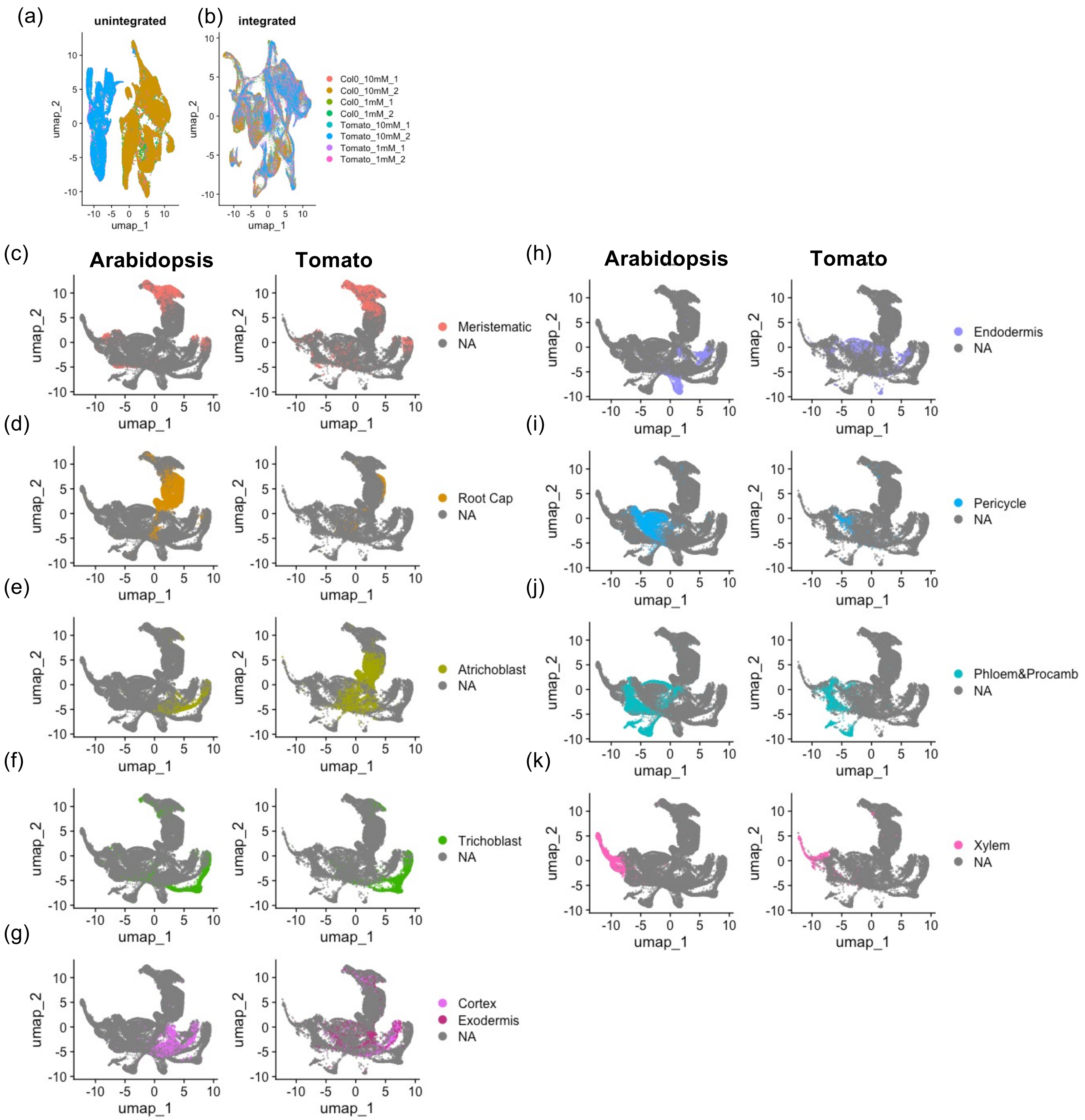


**Supplementary Figure S60. Integration of Arabidopsis (wild-type) and tomato scRNAseq datasets by one-to-one orthologs based on EnsemblPlant homology database.** (**a**,**b**) UMAPs of Arabidopsis and tomato cells before (**a**) and after (**b**) integration. (**c-k**) UMAPs highlighting the same groups of cell types in Arabidopsis (left) and tomato (right).

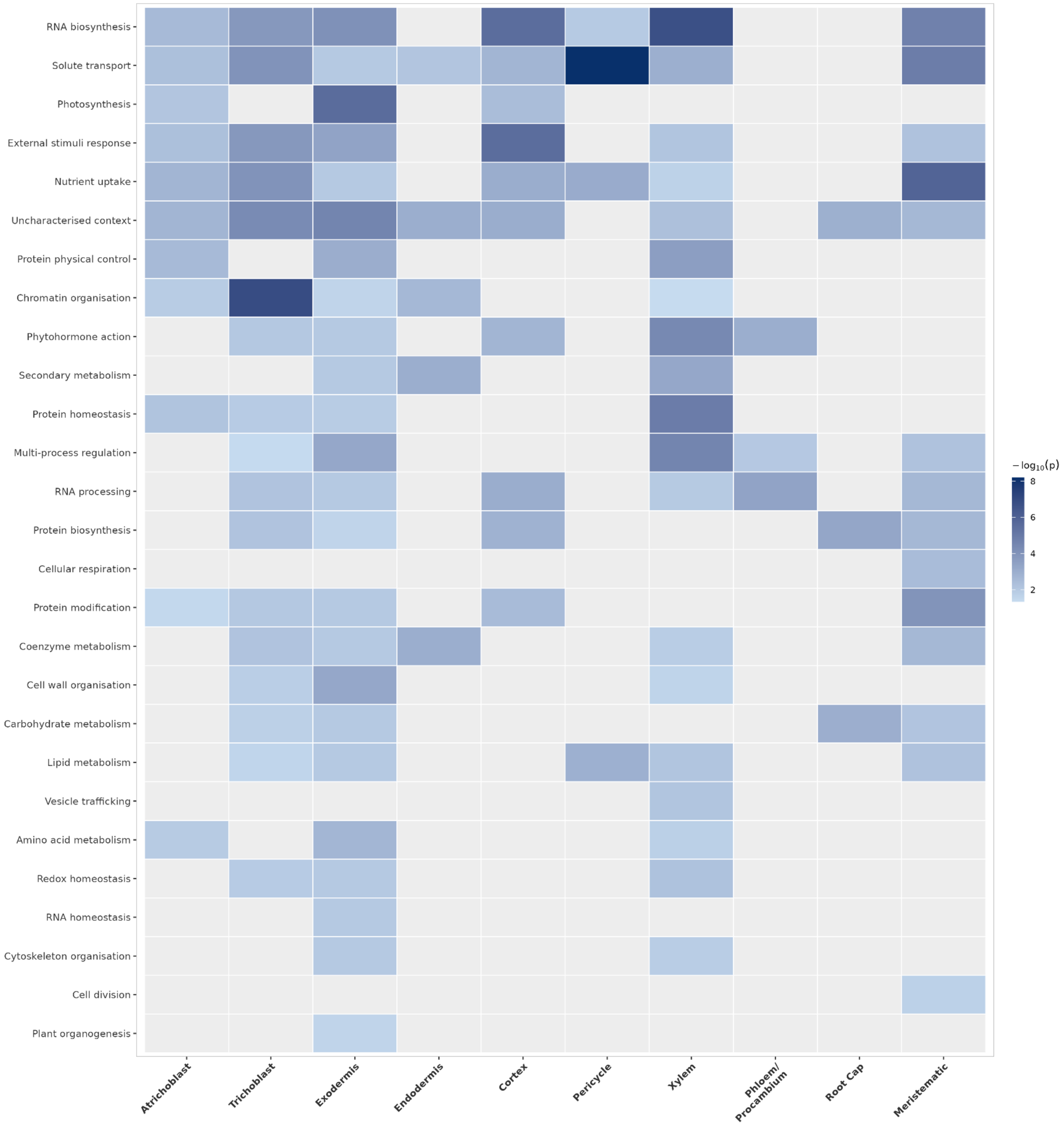


**Supplementary Figure S61. MapMan broad category enrichment of nitrogen-responsive genes across tomato root cell types.** N-responsive differentially expressed genes scRNA-seq data were tested for overrepresentation of MapMan broad functional categories using a one-sided Fisher's exact test against the full MapMan-annotated tomato genome. Up- and down-regulated genes were analysed together per cell type. Tile colour indicates enrichment significance ($-\log_{10}(p)$, light blue to dark navy); grey tiles indicate no significant enrichment ($p \geq 0.05$).

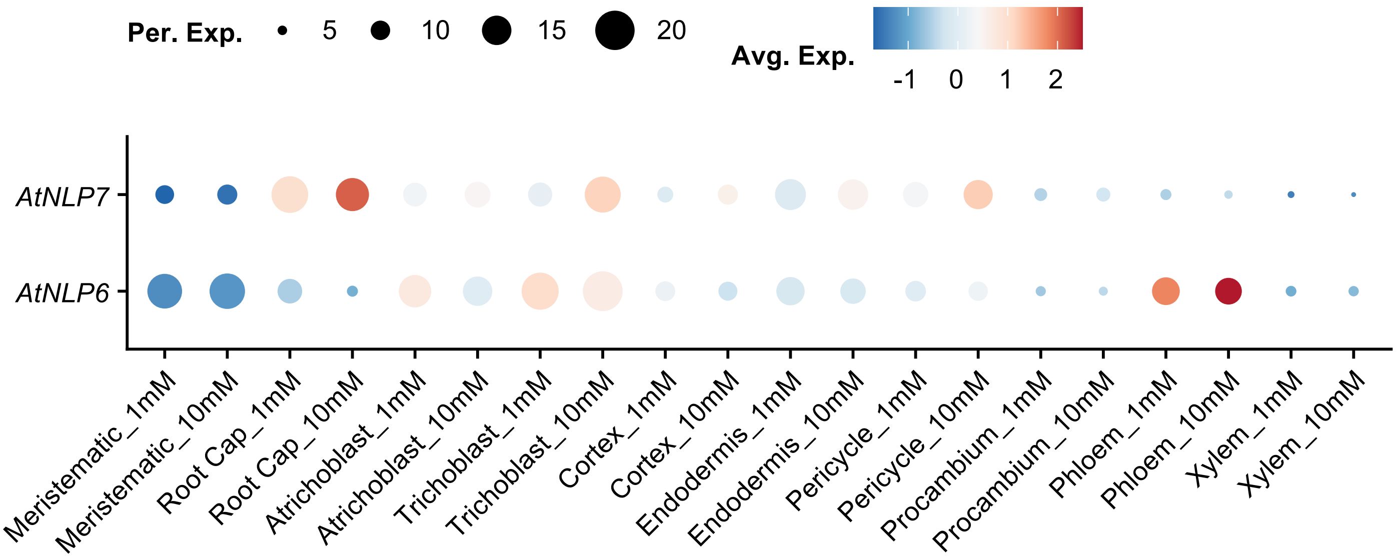


**Supplementary Figure S62. Cell type expression profiles of *AtNLP6* and *AtNLP7* in low (1 mM) vs high (10 mM) $KNO_3$.**

(a)

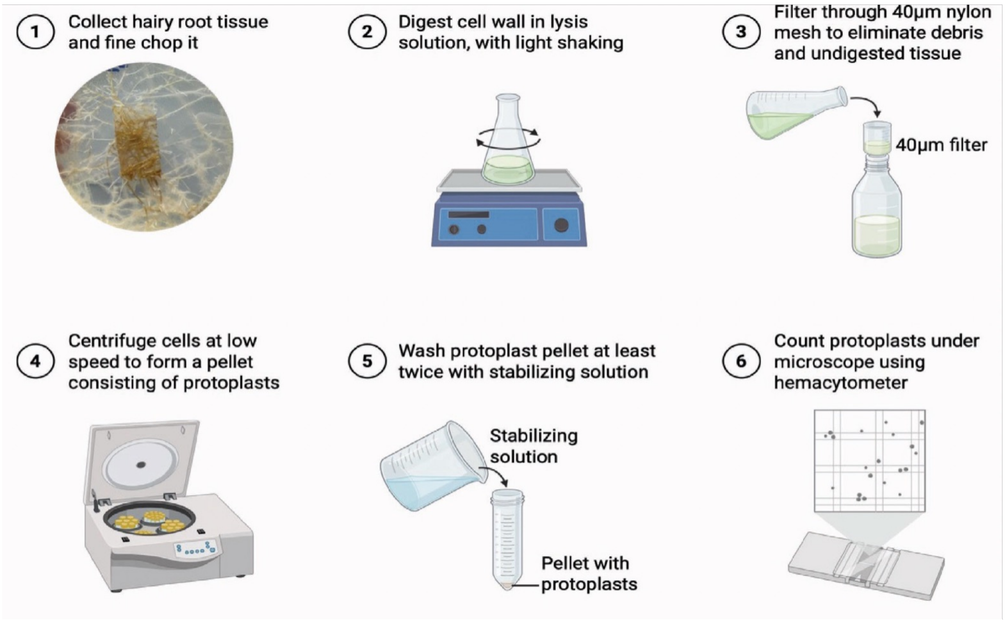


(b)

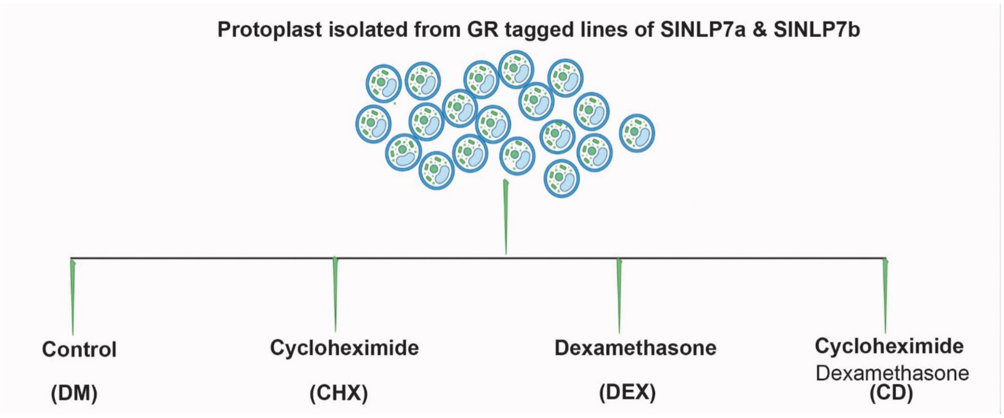


(c)

| Species | Treatment | SlNLP7a-GR | SlNLP7b-GR |
|---|---|---|---|
| *Solanum lycopersicum* | DMSO control (DM) | 1 | 1 |
| *Solanum lycopersicum* | Cycloheximide (CHX) | 1 | 1 |
| *Solanum lycopersicum* | Dexamethasone (DEX) | 1 | 1 |
| *Solanum lycopersicum* | CHX + DEX | 1 | 1 |

**Supplementary Figure S63. Experimental workflow for scRNA-seq of the modified TARGET assay. (a)** Protoplast isolation from hairy roots. Hairy root tissue from GR-tagged SlNLP7a and SlNLP7b lines was chopped, digested, filtered and centrifuged to isolate protoplasts. Protoplasts were counted and divided into four treatment conditions **(b)**: DMSO control (DM), cycloheximide (CHX), dexamethasone (DEX), and cycloheximide + dexamethasone (CD), for single-cell RNA sequencing. **(c)** Sample number for each treatment and hairy root line.

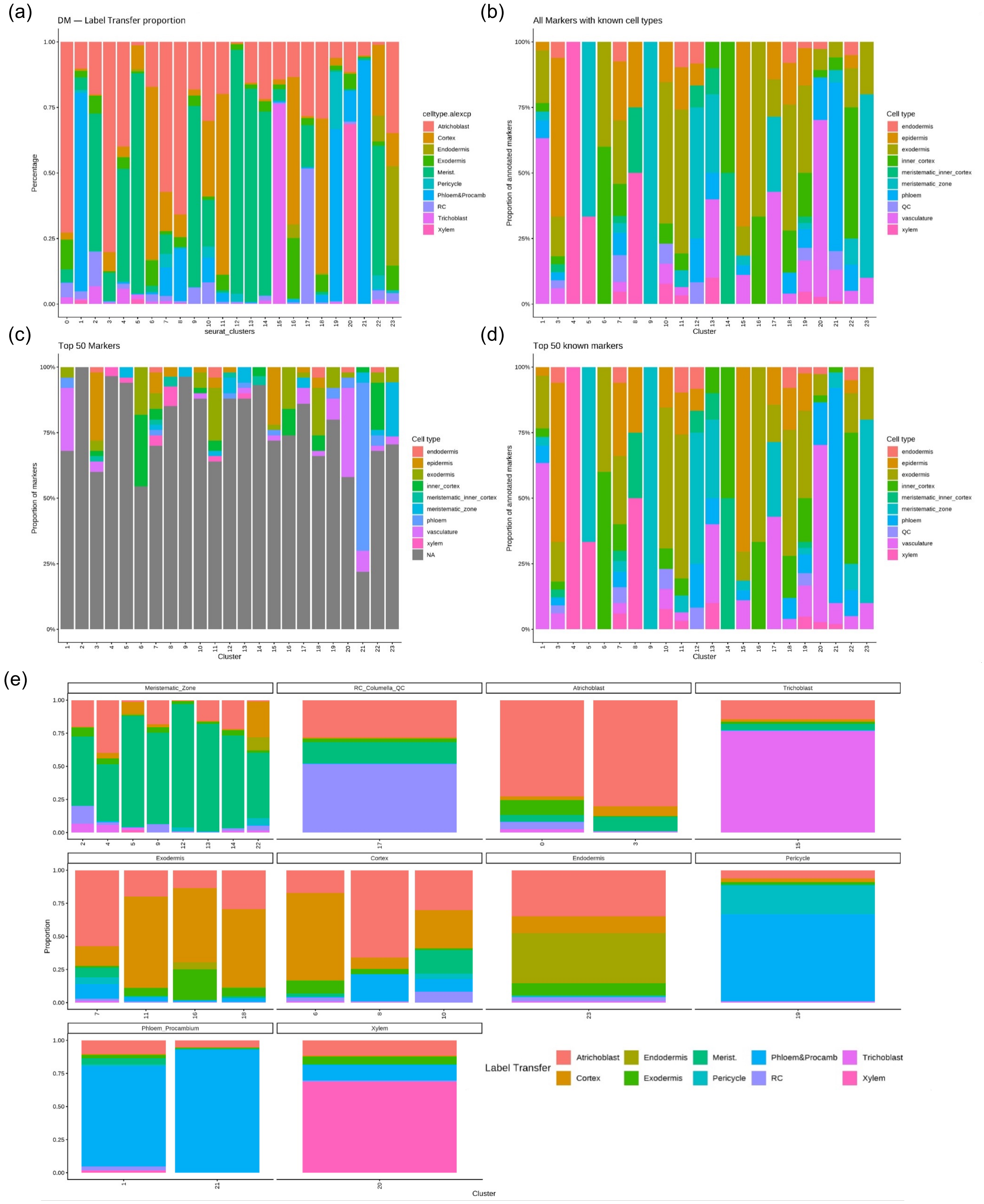


**Supplementary Figure S64. Cell type prediction of tomato root scRNA-seq from the modified TARGET assay. (a)** Percentage of predicted cell type labels in each cluster by label transfer method using the annotated tomato root atlas from Cantó-Pastor et al. (2024) as the reference. **(b)** Percentage of cell types where markers with known expression patterns in each cluster are expressed. **(c)** Percentage of cell types where top 50 markers in each cluster are expressed. **(d)** Percentage of cell types where top 50 markers with known expression patterns in each cluster are expressed. **(e)** Label transfer composition in each annotated cluster.

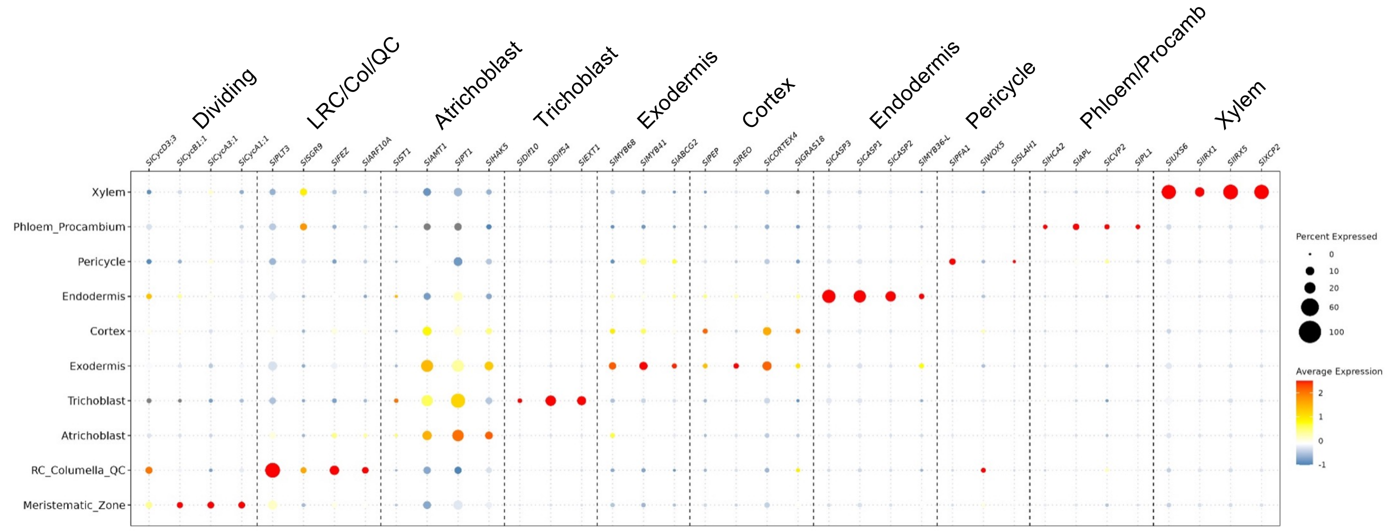


**Supplementary Figure S65. Validation of cell type annotation of tomato scRNAseq dataset from the modified TARGET assay.** Expression of marker genes with known expression patterns in each annotated cell type is shown. RC, root cap. LRC, lateral root cap. Col, columella. QC, quiescent center. Procamb, procambium.

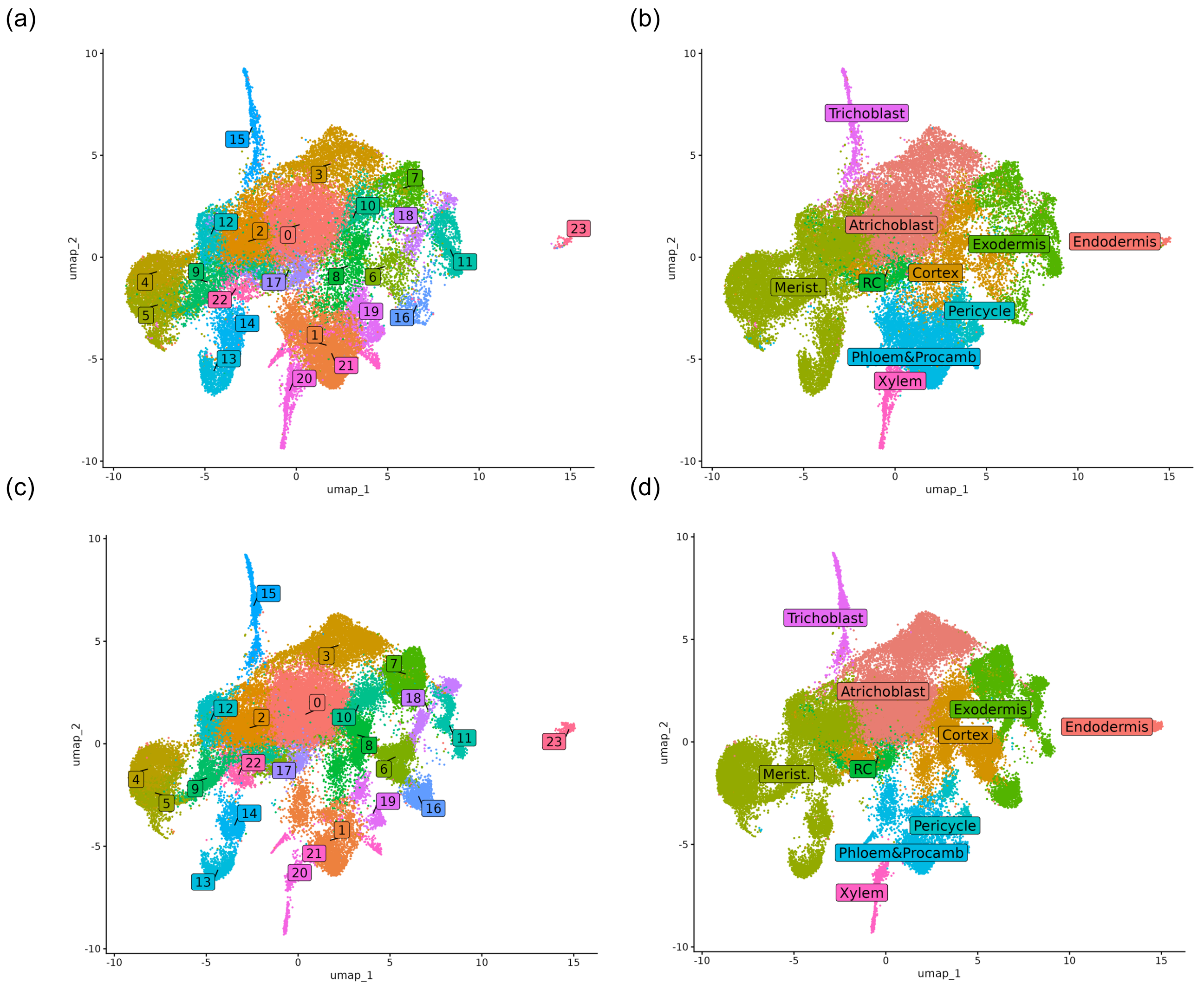


**Supplementary Figure S66. UMAP visualisation of cell type annotations and Seurat clusters of tomato scRNAseq samples from the modified TARGET assay. (a, b)** Seurat clusters (**a**) and cell type annotations (**b**) from SlNLP7a hairy root TARGET samples. (**c, d**) Seurat clusters (**c**) and cell type annotations (**d**) from SlNLP7b hairy root TARGET samples.

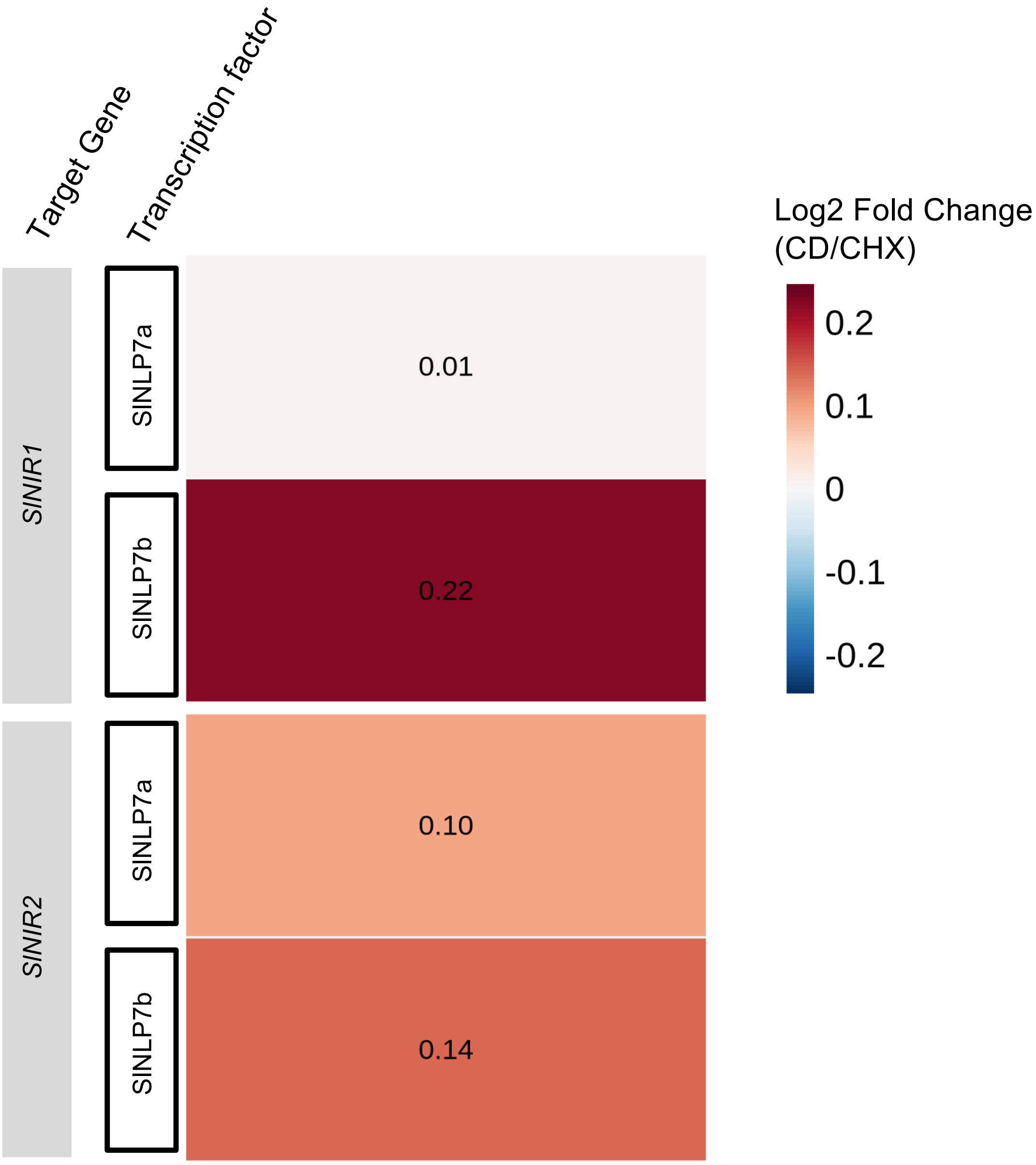


**Supplementary Figure S67. *SlNIR1* and *SlNIR2* were up-regulated in response to CD compared to CHX treatment**. Log2 fold changes (CD/CHX) in the expression of two known SlNLP7a/b targets, *SlNIR1* and *SlNIR2*, calculated from pseudobulked tomato TARGET scRNAseq samples. CD, cycloheximide (CHX) and dexamethasone (DEX) treatment.

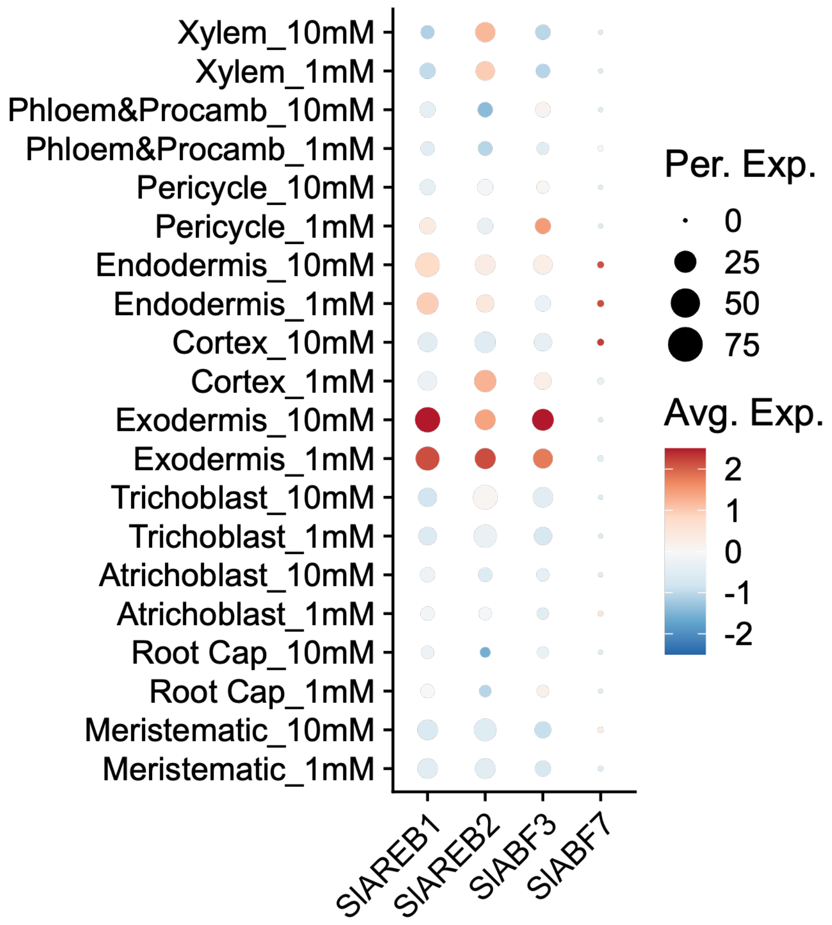


**Supplementary Figure S68. Expression of putative tomato orthologs of *AtABFs* in tomato root cell types at different $KNO_3$ conditions.**
